# Comparative Profiling of Cytosolic Accumulation of Cell-Penetrating Peptides in Mammalian and Bacterial Cells

**DOI:** 10.64898/2026.11.13.766535

**Authors:** Sobika Bhandari, Sarah E. Newkirk, Karl Ocius, Zichen Liu, Mohammad Sharifia, Ananya Naick, Carson Starnes, Rachita Dash, Joey J. Kelly, Yuchen He, George Ongwae, Noah Lucas, Caitlin Fitzgerald, Hobie Bursum, Ryan Kenyon, Marcos M. Pires

## Abstract

Delivering molecules to the cytosol remains a major bottleneck in antibacterial therapy, as both the Gram-negative bacterial envelope and host-cell membranes restrict intracellular access. Cell-penetrating peptides (CPPs) offer a potential strategy for overcoming these barriers. However, selecting CPPs for conjugate development remains largely empirical, in part because conventional fluorescence-based uptake measurements can misidnentify cellular association as cytosolic delivery. Here, we used our recently developed Chloroalkane Azide-tagged Membrane Permeation (CHAMP) assay, a HaloTag-based flow cytometry platform that directly reports cytosolic arrival, to systematically compare 40 CPPs in mammalian cells and *Escherichia coli* (*E. coli*). In mammalian cells, cytosolic accumulation varied substantially within each peptide class, indicating that uptake depends on individual sequence features rather than class membership alone. Among peptides with measurable accumulation in both systems, none exhibited greater apparent accumulation in mammalian cells than in *E. coli*. Complementary assays showed that several peptides reached the bacterial cytosol without detectable membrane damage, whereas others accumulated in association with compromised membrane integrity. Together, these findings provide a quantitative comparison of CPP-mediated cytosolic access in mammalian and bacterial cells and establish a framework for selecting peptide carriers for antibiotic conjugates.

## INTRODUCTION

Antibiotic resistance is among the leading threats to human health. A global analysis estimated that bacterial antimicrobial resistance was associated with 4.71 million deaths in 2021, including 1.14 million deaths directly attributable to resistance, and projected 39.1 million attributable deaths between 2025 and 2050.^1^ One major challenge in developing effective antibiotics is achieving sufficient intracellular accumulation, in addition to engaging the intended target. Whereas mammalian cells are enclosed by a plasma membrane, Gram-negative bacteria possess a multilayered envelopes comprising an inner membrane, a peptidoglycan layer, and an outer membrane (**Fig. 1a**). Together, these structures impose additional barriers to molecular permeation.^2^ Consequently, relatively few compounds achieve substantial intracellular accumulation in Gram-negative bacteria.^3^ This challenge is further compounded when pathogens reside within host cells, because effective treatment requires compounds to cross both the host-cell membrane and the bacterial envelope. Many antibiotics, including aminoglycosides, exhibit limited permeation across eukaryotic membranes, further restricting their access to intracellular pathogens.^4^ More broadly, membrane permeability remains a major obstacle to intracellular drug delivery, particularly for large, hydrophilic molecules such as peptides and proteins.^5,6^ Thus, delivering therapeutics across both bacterial and mammalian membranes remains a fundamental challenge, motivating considerable efforts to understand the physicochemical properties that govern membrane permeation.

**Figure 1.**
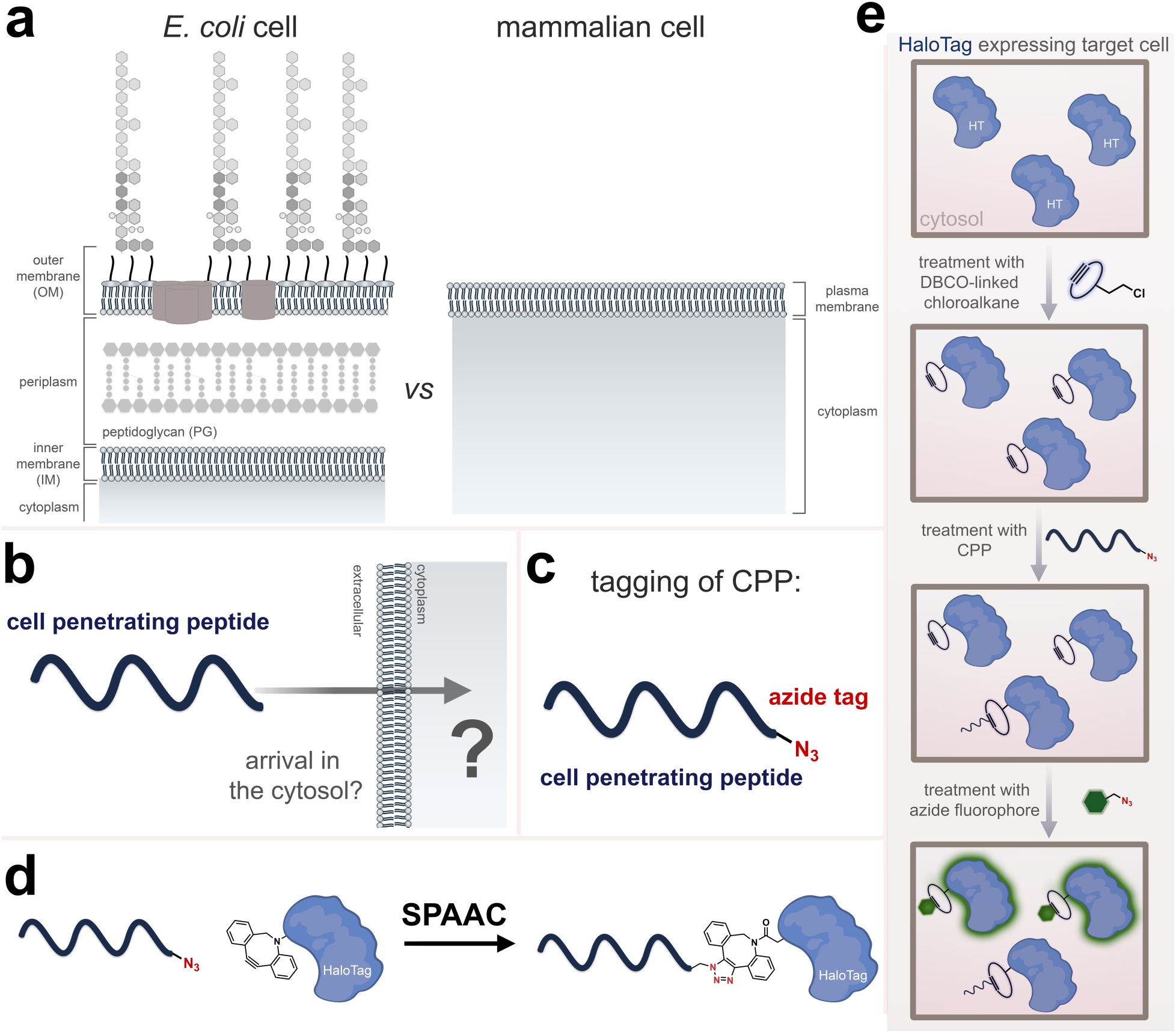
**a** Schematic representation of Barriers to cytosolic delivery of CPPs in bacterial and mammalian cells. **b**. General Scheme of Principle of CHAMP. Cells express HaloTag (HT) in the cytosol (top). Treatment with a DBCO-linked chloroalkane anchors a strained alkyne to HaloTag (middle). Azide-tagged test molecules that reach the cytosol react with the anchored DBCO by strain-promoted azide–alkyne cycloaddition and block it, and a subsequent azide fluorophore (green) labels the remaining DBCO sites (bottom). Fluorescence therefore decreases as cytosolic accumulation increases.

Among the strategies explored to overcome these barriers, cell-penetrating peptides (CPPs) have emerged as promising molecular delivery vehicles. Following the discovery that HIV-1 Tat protein can enter cells^7,8^ and that its truncated basic domain can accumulate in the nucleus,^9^ numerous CPPs with related properties have been identified or rationally designed.^10,11^ CPPs are short peptides commonly classified as cationic, amphipathic, or hydrophobic according to their physicochemical properties.^12–14^ Their relatively small size and ability to facilitate cellular uptake have made them attractive delivery vehicles.^15^ Despite their widespread use, however, the mechanisms that govern CPP internalization and cytosolic delivery remain incompletely understood. This uncertainty complicates CPP classification and limits our ability to predict their performance across different cell types.^16^

In antibacterial research, CPPs have been investigated as delivery vehicles to enhance antibiotic uptake through coadministration or covalent conjugation.^17^ For example, a cleavable conjugate of kanamycin and the antibacterial CPP P14LRR demonstrated greater activity against intracellular *Mycobacterium tuberculosis* in macrophages and reduced *Salmonella* burden in a *Caenorhabditis elegans* infection model compared with its noncleavable counterpart.^4^ This conjugate was subsequently evaluated against biofilms and ESKAPE pathogens.^18^ CPPs have also been used to facilitate the intracellular delivery of antisense agents into bacteria. For instance, conjugation of (KFF)3K to peptide nucleic acids improved their antisense activity against *E. coli.*^19^

Despite substantial effort, it remains unclear which CPPs most effectively reach the cytosol of a given target cell (**Fig. 1b**).^20^ Many studies have examined CPP internalization in mammalian cells, including the mechanisms of uptake and intracellular localization.^21,22^ A central challenge is that most internalization assays measure total cell-associated signal rather than the fraction that has escaped endosomal compartments. Consequently, a peptide trapped in vesicles may appear indistinguishable from one that has reached the cytosol.^23^ Although several studies have demonstrated cytosolic delivery by CPPs in mammalian cells, few have directly compared CPP accumulation in bacterial and mammalian cells.^24,25^ These cell types present fundamentally different barriers. In mammalian cells, CPP uptake often involves endocytosis, followed by the need for endosomal escape. In Gram-negative bacteria, by contrast, a peptide must traverse the outer membrane, periplasm, and inner membrane without vesicular trafficking. It therefore remains unclear whether the ability of a CPP to accumulate in the mammalian cytosol predicts its accumulation in bacteria.

Current methods for assessing membrane permeability and intracellular accumulation include direct compound quantification by liquid chromatography-tandem mass spectrometry (LC-MS/MS),^26^ as well as standardized permeability assays such as the parallel artificial membrane permeability assay (PAMPA), typically quantified by UV spectrophotometry,^27^ and cell-based assays such as Caco-2.^28^ Mass-spectrometry-based approaches, however, are generally low-throughput and may not readily distinguish membrane-associated compounds from those that have reached the cytosol.^29,30^ Fluorescent labeling provides an alternative for visualizing cellular uptake and intracellular localization,^30^ but fluorescence measurements alone generally cannot distinguish membrane-associated or endosomally entrapped peptides from cytosolic peptides.^31^ In addition, attaching a fluorophore can substantially alter a molecule’s physicochemical properties and may affect its biological activity, localization or intracellular trafficking. Methods that directly quantify cytosolic accumulation while minimally perturbing the test molecule are therefore needed.

One strategy for confirming cytosolic arrival is to make signal generation dependent on a covalent capture event that occurs only within the cytosol. The HaloTag-based chloroalkane penetration assay (CAPA) directly reports the cytosolic arrival of molecules in mammalian cells.^31,32^ Our group subsequently extended this approach to various bacterial species, including *E. coli* and mycobacteria, through BaCAPA, and to bacteria residing inside mammalian host cells.^2^ Building on this platform, we developed the Chloroalkane Azide-tagged Membrane Permeation (CHAMP) assay, which replaces the chloroalkane tag on a test molecule with a substantially smaller azide group to minimize the tag’s influence on permeability (**Fig. 1c**).^33^ In CHAMP, HaloTag-expressing cells are first treated with a chloroalkane-modified strained alkyne. The azide-tagged test molecule is then added and, if it reaches the cytosol, undergoes strain-promoted azide–alkyne cycloaddition (SPAAC) with the alkyne-functionalized HaloTag (**Fig. 1d**). In a subsequent chase step, a fluorophore-conjugated azide labels the remaining unoccupied alkyne groups through a selective, rapid, and essentially irreversible reaction.^33,34^ Fluorescence intensity therefore decreases as cytosolic accumulation of the test molecule increases, enabling high-throughput analysis by flow cytometry (**Fig. 1e**).^32^ We have used this assay to assess molecular accumulation in mycobacteria,^35^ *E. coli*,^36^ and mammalian cells.^37^ More recently, we demonstrated that fatty acid acylation can have markedly different effects on small molecule accumulation in bacterial and mammalian cells.^38^

In this work, we used CHAMP to compare the cytosolic accumulation of 40 CPPs spanning cationic, amphipathic, and hydrophobic classes in mammalian cells and in *E. coli.* To distinguish cytosolic delivery from apparent accumulation associated with membrane damage, we paired these measurements with cell viability and membrane integrity assays. This combined approach allowed us to identify CPPs that reached the cytosol with little evidence of membrane damage, as well as peptides whose accumulation coincided with compromised membrane integrity. Our findings provide a quantitative framework for evaluating CPPs as potential carriers for antibiotic conjugates.

## RESULTS AND DISCUSSION

CPPs are not a mechanistically uniform class: cationic, amphipathic, and hydrophobic sequences can interact with membranes through distinct mechanisms, and no single peptide is representative of the entire group. Identifying the features that govern cytosolic entry in different cell types therefore requires a panel that samples a broad range of sequence and charge properties. Mammalian and bacterial cells present fundamentally different barriers to peptide entry (**Fig 1a**). Mammalian cells are enclosed by a single plasma membrane enriched in cholesterol and sphingolipids and characterized by relatively low surface-charge density.^39^ CPPs can enter mammalian cells through endocytic pathways,^12,40^ after which they must escape endosomal compartments to reach the cytosol. In contrast, Gram-negative bacteria lack endocytic pathways. Peptides must instead cross the outer membrane, including its lipopolysaccharide (LPS) layer, the peptidoglycan layer, and the inner membrane to reach the cytoplasm.^3^ These differences raise the question of whether CPPs that accumulate efficienctly in mammalian cells behave similarly in bacteria. To address this question, we evaluated a library of 40 CPPs in HaloTag-expressing HeLa cells and *E. coli* using the CHAMP assay (**Fig 1b**).

### Cytosolic accumulation of a diverse CPP library in mammalian cells

To compare cytosolic accumulation across CPP classes, we synthesized a library of 40 CPPs comprising 13 cationic (C1–C13), 21 amphipathic (A1–A21), and 6 hydrophobic (H1–H6) CPPs (**Fig. 2**). Each CPP carried an *N*-terminal azide tag. The library was designed following the approach reported by the Numata laboratory, which screened a CPP library in plants cells.^41^ It included representatives widely studied delivery peptides, including Tat-derived peptides, polyarginine derivatives, Transportan, PolyP3(SAP), and crotamine-derived peptides.^42–44^ We first assessed the cytosolic accumulation of each peptide in HaloTag-expressing mammalian cells using the CHAMP assay at two time points (**Fig 3a**).^45^ Accumulation varied substantially among peptides within each class, indicating that cytosolic uptake was strongly sequence-dependent. Most peptides produced a detectable response after 2 h. Apparent accumulation increased between 2 and 4 h for **C1**, **C13**, **A18** and **H6**, whereas **A1** changed little, and **H1**, **A16** and **A19** showed decreased apparent accumulation (**Fig. S1**). Thus, these findings highlight that both peptide sequence and incubation time influence the measured extent of cytosolic accumulation.

**Figure 2.**
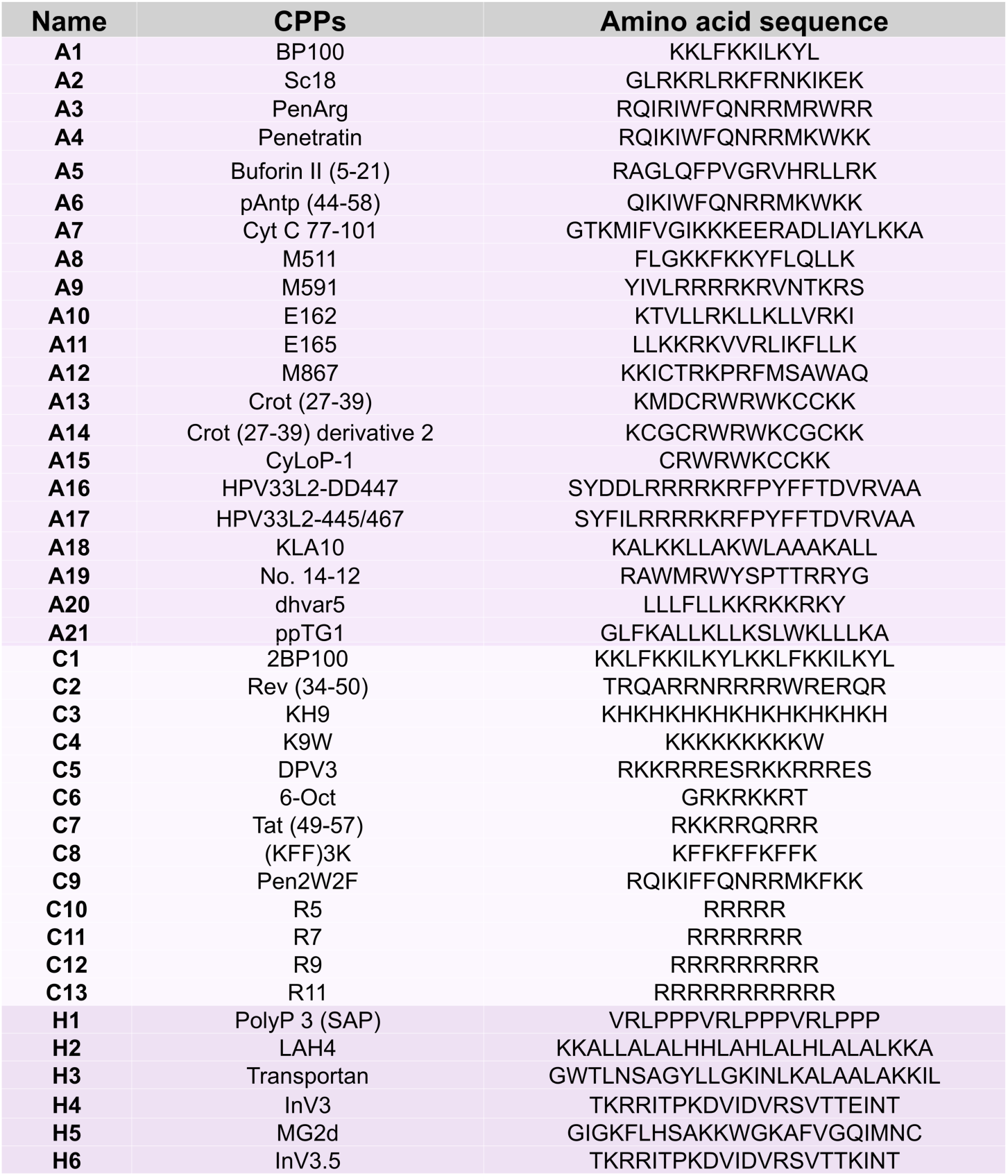
**a**. Chemical structures of the amphipathic (**A1**–**A21**), cationic (**C1**–**C13**) and hydrophobic (**H1**–**H6**) CPPs. The N-terminal azide tag is [highlighted in red]. **b**. Codes, names and amino acid sequences (one-letter code, N to C). Peptides were synthesized by solid-phase peptide synthesis [with N-terminus azide].

**Figure 3.**
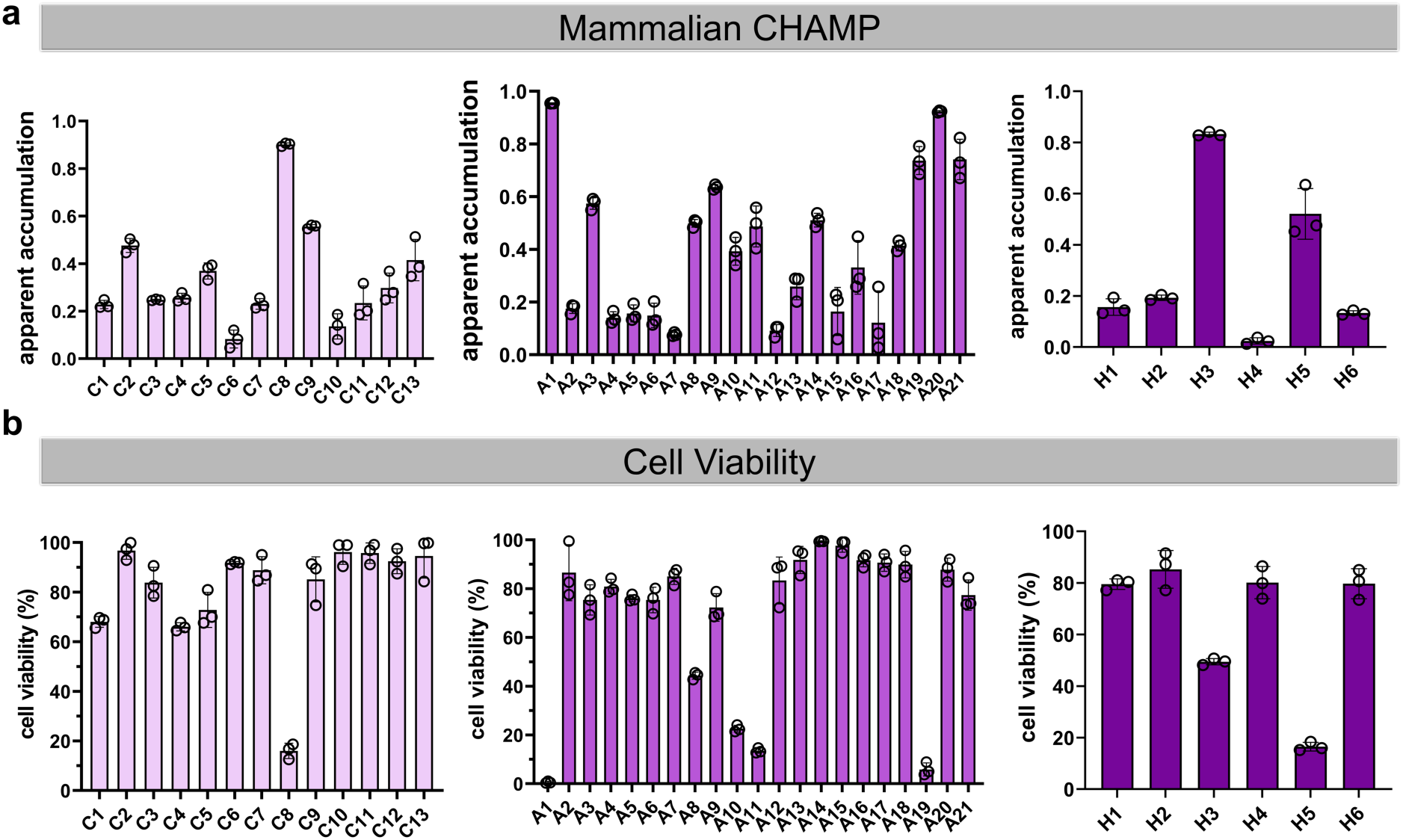
**a**. Apparent accumulation measured by CHAMP in HaloTag-expressing HeLa cells for cationic (**C1**–C13), amphipathic (**A1**–**A21**) and hydrophobic (**H1**–**H6**) CPPs. HaloTag were treated with 10 μM of DBCOcl for 15 minutes, pulsed with 50 μM of azide-tagged CPPs for 1 h, chased with 50 μ TMRaz for 1 h. **b**. Viability of HeLa cells after treatment with each peptide at 50 μM for 2h, measured by MTT assay and expressed as a percentage of untreated cells.

Within the cationic class, **C8** showed the greatest apparent accumulation at both time points, whereas **C6** and **C10** showed the least. Increasing the number of positively charged residues, as in **C2**, **C3**, **C5**, and **C9**, did not uniformly increase accumulation. Across the full set, neither net charge nor arginine content alone explained the observed differences. A clearer pattern emerged when aromatic content was considered. **C1**, **C8**, and **C9**, the three peptides containing at least three aromatic residues, ranked among the four strongest accumulators in this class despite having net charges ranging from +4 to +10. Aromatic content may therefore contribute substantially to accumulation across the cationic panel. Among the remaining peptides, which contained no more than one aromatic residue, arginine content and net charge appeared to be more informative. These observations suggest that aromatic side chains and cationic residues may each contribute to cytosolic accumulation, although the relative importance of these features depends on the sequence context.

The contribution of cationic residues also appeared to depend on amino acid identity rather than formal charge alone. At a matched net charge of +9, the polyarginine peptide **C12** accumulated 1.5-to 1.9-fold more than the lysine-based peptides **C3** and **C4,** while the arginine-rich **C2** accumulated to an even greater extent. Within the homologous polyarginine series, apparent accumulation increased with chain length at both time points, from **C10** through **C11** and **C12** to **C13**. This trend is consistent with reports that arginine-rich peptides interact with membranes through multidentate hydrogen bonding between guanidinium groups and membrane phosphates.^46^ The time dependence also differed between peptides: **C8** appeared to reach a plateau by the earlier time point, whereas the polyarginines continued to accumulate, with the magnitude of the increase scaling with chain length. These kinetic differences may reflect distinct uptake processes, although the present data do not establish the underlying mechanisms.

Within the amphipathic class, several CPPs demonstrated high apparent cytosolic accumulation, although substantial differences were also evidence across the panel. **A1** is a short cationic antimicrobial peptide derived from a hybrid of cecropin A, an antimicrobial peptide from the moth *Hyalophora cecropia*, and melittin M, a membrane-active component of bee venom.^47^ **A1** has reported activity against Gram-negative bacteria, raising the possibility that some features associated with bacterial activity may also influence accumulation in mammalian cells. Its derivative, **C1**, showed comparatively low cytosolic accumulation in HeLa cells. Several previously reported CPPs accumulated poorly, including **A5**, **A6**, **A7**, **A12**, **A15**, and **A17**, whereas engineered amphipathic peptides such as **A9** and **A10** accumulated strongly. Although these peptides differ in sequence, their amphipathic character may facilitate interactions with membrane phospholipids while supporting insertion of hydrophobic residues into the lipid bilayer. Such properties have been associated with membrane translocation and intracellular delivery.^48,49^

Peptides **A13** and **A14** differ in cysteine content and net charge. **A14** showed approximately twofold greater apparent cytosolic accumulation than **A13** in HeLa cells after 2 h. One possible explanation is that cysteine residues can promote uptake through thiol-disulfide exchange with cell-surface thiols. The increased net charge of **A14** may also strengthen interactions with anionic cell-surface components, including proteoglycans and phospholipids, thereby increasing the local peptide concentration at the membrane.^50^ **A16** and **A17**, which differ in hydrophobicity, also showed different accumulation profiles: the more hydrophobic peptide, **A17**, had lower apparent accumulation than **A16**. This observation suggests that increasing hydrophobicity does not necessarily enhance cytosolic accumulation and may instead favor membrane association or partitioning into the lipid bilayer without efficient translocation.^51–53^ Together, these comparisons suggest that efficient cytosolic accumulation may require a balance between hydrophobicity and polarity rather than simply increased lipophilicity.

Within the hydrophobic class, accumulation also varied substantially among peptides. **H6**, a derivative of **H4** in which a glutamic acid residue is replaced by lysine, showed approximately threefold greater apparent cytosolic accumulation after 2 h. This difference is consistent with the possibility that additional cationic charge promotes interactions with anionic cell-surface components, including phosphatidylserine, sialic acids, and heparan sulfate proteoglycans, thereby increasing the local peptide concentration at the cell surface.^50,54^ **H3**, a 27-amino acid chimeric peptide previously reported to transport small proteins and nucleotide oligomers through direct membrane interactions, showed the greatest apparent accumulation in this class.^55^

Although peptides from all three classes reached the cytosol, accumulation varied more strongly among individual sequences than among classes. To determine whether the low apparent accumulation of selected peptides reflected limited cytosolic access rather than an inability to react with cytosolic HaloTag, we permeabilized the plasma membrane with digitonin, which complexes membrane cholesterol and has been used to facilitate intracellular entry.^33,56^ We selected one poorly accumulating peptide from each class: **C6**, **A7**, and **H4**. Digitonin treatment increased their apparent accumulation to approximately 50% of the maximum signal (**Fig. S2**). This result indicates that the azide-tagged peptides could reach and react with cytosolic HaloTag when membrane access was facilitated. Their low signals under untreated conditions are therefore consistent with limited cytosolic accumulation.

Because membrane-active peptides can disrupt cellular membranes and reduce cell viability, we next assessed whether differences in apparent accumulation could associated with peptide-induced toxicity using an MTT assay (**Fig. 3b**). The CPP library produced a broad range of effects on HeLa cell viability, from little or no detectable loss reductions in viability below 20% viable. Several peptides, including **A1**, **A11**, **A19**, **C8**, and **H5**, reduced cell viability to below 20%, whereas most peptides had comparatively modest effects. These results indicate that cytotoxicity is not determined solely by net cationic charge and may also depend on amino acid composition, hydrophobicity, amphipathicity, secondary structure, and peptide conformation.^57^ Importantly, high apparent cytosolic accumulation did not consistently correspond to increased cytotoxicity. For example, **C9**, **A3**, and **A20** showed efficient accumulation while retaining relatively high cell viability, whereas **A11** and **H6** exhibited substantial toxicity without ranking among the strongest accumulators. This lack of a direct relationship suggests that cytosolic accumulation and loss of cell viability are not equivalent readouts. More broadly, total cellular uptake, endosomal uptake, and cytosolic delivery represent distinct processes; efficient internalization does not necessarily entail plasma-membrane disruption or cell death.^58,59^

### Cytosolic accumulation of a diverse CPP library in bacterial cells

To determine whether CPPs that accumulated efficiently in mammalian cells behaved similarly in bacteria, we evaluated the same panel of 40 peptides using CHAMP in HaloTag-expressing *E. coli* (**Fig. 4a**).^60^ Molecules entering bacterial cells must overcome multiple permeability barriers to reach the cytoplasm, and bacteria lack the endocytic pathways available to mammalian cells.^61^ We therefore anticipated that the determinants of accumulation might differ between the two systems. Most peptides showed substantial apparent accumulation in *E. coli*: 17 of 21 amphipathic, 10 of 13 cationic, and 3 of 6 hydrophobic peptides produced more than a sevenfold increase in apparent accumulation. Hydrophobic peptides showed the greatest variability. The principal exceptions were **C4**, **C6**, **A7**, **H1**, and **H4**. Notably, **C7** showed markedly greater apparent accumulation in bacteria than in mammalian cells. Several amphipathic peptides accumulated strongly in both systems, suggesting that amphipathic character may be favorable across these distinct biological contexts. By contrast, the hydrophobic peptides showed greater variability than the cationic and amphipathic groups: **H3** accumulated efficiently, whereas **H1**, **H4,** and several other members of this class did not. These results indicate that CPP performance cannot be assumed to transfer across biological membranes. CPP performance should therefore be evaluated in the biological system relevant to the intended application rather than inferred from mammalian cell uptake alone.

**Figure 4.**
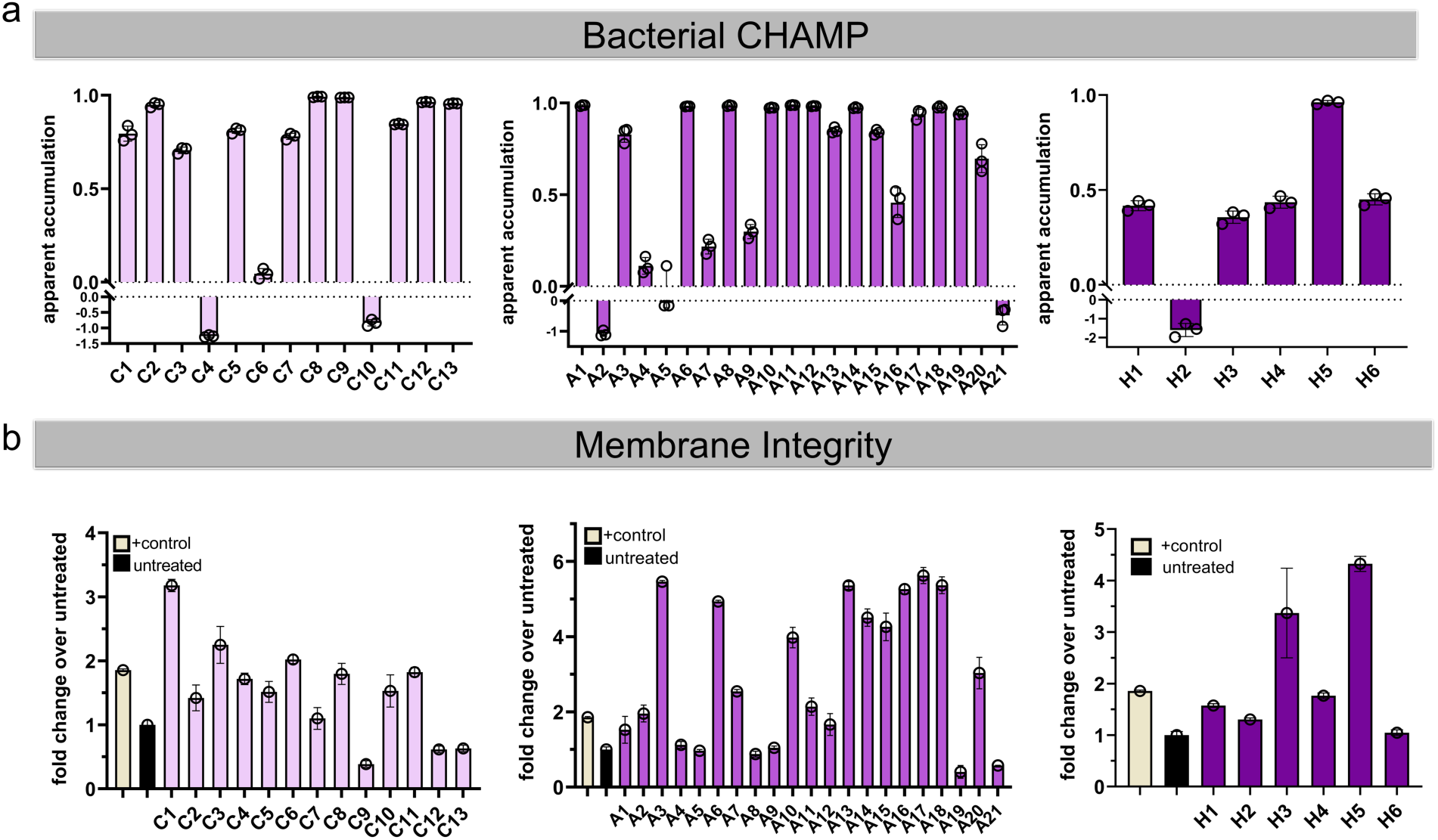
**a**. Measurement of cytosolic accumulation in *E.coli*. Apparent accumulation measured by CHAMP in HaloTag-expressing *E. coli* for cationic (**C1**–**C13**), amphipathic (**A1**–**A21**) and hydrophobic (**H1**–**H6**) CPPs. Cells were treated with 50 µM chloroalkane– DBCO (30 min), pulsed with 50 μM azide-tagged CPPs (1 h) and chased with 50 μM TMRaz (1 h). **b**. Membrane integrity assessed by SYTO9 fluorescence of peptide-treated cells at 50μM and dye at 10 μM for 1h shown as fold change over untreated cells. Polymyxin B. is used as positive control

We next asked whether the bacterial outer membrane (OM) limits the accumulation of selected CPPs. Cells were pretreated with polymyxin B nonapeptide (PMBN), which permeabilizes the OM without substantially killing cells.^62^ PMBN increased the apparent accumulation of **C1**, **A7** and **H3** (**Fig. S3**), consistent with the OM limiting the entry of these peptides under the conditions tested. We also compared **R9** (the all L-enantiomer) with its D-enantiomer, **r9**. In our previous study, **R9** entered mammalian cells more efficiently than **r9**.^33^ In *E. coli*, however, **r9** showed approximately 1.8-fold greater apparent accumulation than **R9** (**Fig. S4**). D-oligoarginines are resistant to proteolysis and have been reported to show enhanced uptake relative to **R9** in mammalian cells.^63^ The difference observed in *E. coli* is therefore consistent with proteolytic degradation limiting the accumulation of **R9**, although this possibility was not directly tested here.

To determine whether bacterial accumulation was accompanied by membrane damage, we stained peptide-treated *E. coli* with SYTO9 and propidium iodine (PI) (**Fig. 4b and Fig. S5**).^64^ SYTO9 is a cell-permeant nucleic acid stain that labels bacteria, whereas PI preferentially enters cells with compromised membranes and can reduce SYTO9 fluorescence through dye displacement. The two signals must therefore be interpreted together when assessing membrane integrity.^65^ Several peptides that accumulated efficiently produced fluorescence signals for both dyes that remained near untreated-cell levels, including **C7**, **A4**, **A5**, **A8**, **A9**, **C2** and **C5**. For **A5**, this observation is consistent with previous reports that it can enter *E. coli* without causing extensive membrane lysis.^66^ In contrast, **A10**, **A13**, **A15**, **A17**, **A20** and **H5**, produced signals for both dyes, consistent with membrane perturbation accompanying peptide accumulation. **C12** and **C13** decreased SYTO9 fluorescence while increasing signal from PI fluorescence, a pattern consistent with compromised membrane integrity and with reports that oligoarginines can damage *E. coli* membranes during cytoplasmic entry.^67^ A further group (**C1**, **C3**, **A3**, **A6**, **A11**, **A14**, **A16**, **A18** and **H3**) increased SYTO9 fluorescence without a comparable increase in PI signal. This pattern may reflect changes in dye accessibility or partial envelope perturbation, but does not by itself establish the extent or location of membrane damage. Overall, several peptides accumulated without detectable changes in these membrane-integrity readouts, whereas others showed evidence of membrane perturbation. Conversely, **A7**, **C6**, **H1** and **H4** increased SYTO9 fluorescence despite showing relatively low accumulation, indicating that changes in dye signal do not necessarily predict cytosolic access. Because many peptides approached the upper range of the apparent accumulation response, the dye signals did not quantitatively track cytosolic accumulation. These observations, together with previous reports that eukaryotic CPPs enter bacteria and exert antimicrobial effects,^68^ underscore the importance of measuring accumulation alongside membrane integrity.

To assess OM permeabilization more directly, we performed an N-phenyl-1-naphthylamine (NPN) uptake assay.^69^ NPN uptake profiles (**Fig. S6**) showed time- and peptide-dependent responses. Polymyxin B, included as a positive control, produced the strongest permeabilization response relative to untreated cells. Several CPPs induced measurable OM permeabilization, although their responses were generally lower than that of polymyxin B and varied substantially across the peptide panel. Collectively, these findings suggest that OM permeabilization does not necessarily predict cytosolic accumulation. Bacterial entry therefore appears to depend on additional peptide-specific properties, highlighting the importance of evaluating OM permeabilization, overall membrane integrity, and cytosolic accumulation as distinct, complementary readouts.

## CONCLUSIONS

Delivering molecules to the cytosol remains a major challenge in antibacterial therapy, and CPPs are attractive potential carriers whose cytosolic access has rarely been measured directly. Here, we used CHAMP to quantify the apparent cytosolic accumulation of 40 CPPs in HeLa cells and *E. coli*. In both systems, accumulation varied substantially among individual peptide sequences, demonstrating that CPP class alone is insufficient to predict performance. Accumulation in mammalian cells did not reliably predict accumulation in bacteria; indeed, most peptides showed greater apparent accumulation in *E. coli* under the conditions tested. The measured accumulation was also not uniformly associated with membrane damage. Several peptides reached the bacterial cytosol without detectable changes in the membrane-integrity assays, whereas others accumulated alongside evidence of membrane perturbation. In addition, OM permeabilization did not consistently predict cytosolic arrival. These findings support evaluating CPPs in the intended target system and pairing cytosolic accumulation measurements with independent assessments of membrane integrity. Peptides that accumulate efficiently in *E. coli* without substantial membrane damage may be promising starting points for antibiotic conjugate development. Future studies will be needed to determine whether conjugation preserves their accumulation profiles and improves antibacterial activity.

## ACKNOWLEDGEMENTS

This study was supported by NIH grants 1R01AI178975-01 (M.M.P.), R35GM124893 (M.M.P.), and R01AI179080-01 (M.M.P.).

