## Supplementary Information for "Comparative Profiling of Cytosolic Accumulation of Cell-Penetrating Peptides in Mammalian and Bacterial Cells"

### TABLE OF CONTENTS


| Supplementary Figures | |
|---|---|
| Cytosolic Accumulation of CPPs in HT cells for 4h | **P3-P4** |
| Digitonin Permeabilization Uptake of CPPs | **P4** |
| Outer Membrane Permeabilization using PMBN | **P5** |
| Effect of Stereochemistry on CPP accumulation in *E.coli.* | **P5** |
| Membrane integrity assessed via Propidium Iodide | **P6-P7** |
| OM permeabilization of *E. coli* by CPPs measured by NPN uptake. | **P8** |
| **Materials and Methods** | |
| Materials | **P9** |
| **Experimental methods** | |
| Mammalian cell culture | **P9** |
| Mammalian CHAMP assay | **P9** |
| Digitonin assay | **P9** |
| MTT cell viability assay | **P10** |
| Expression of HaloTag protein and Bacterial cell culture | **P10** |
| Bacterial CHAMP Assay | **P10** |
| Bacterial Assay with PMBN | **P11** |
| Membrane Integrity Assay via SYTO9 | **P11** |
| Propidium Iodide Assay | **P11** |
| NPN uptake Assay | **P12** |
| **Synthesis, Characterization, and Structure of Test Molecules** | |
| Synthesis of peptides via SPPS | **P12** |
| Analytical HPLC Chromatogram and MALDI-TOF | **P13-P53** |
| Structure of libraries of CPPs | **P54-P64** |
| **References** | **P65** |

#### Supplementary Figures

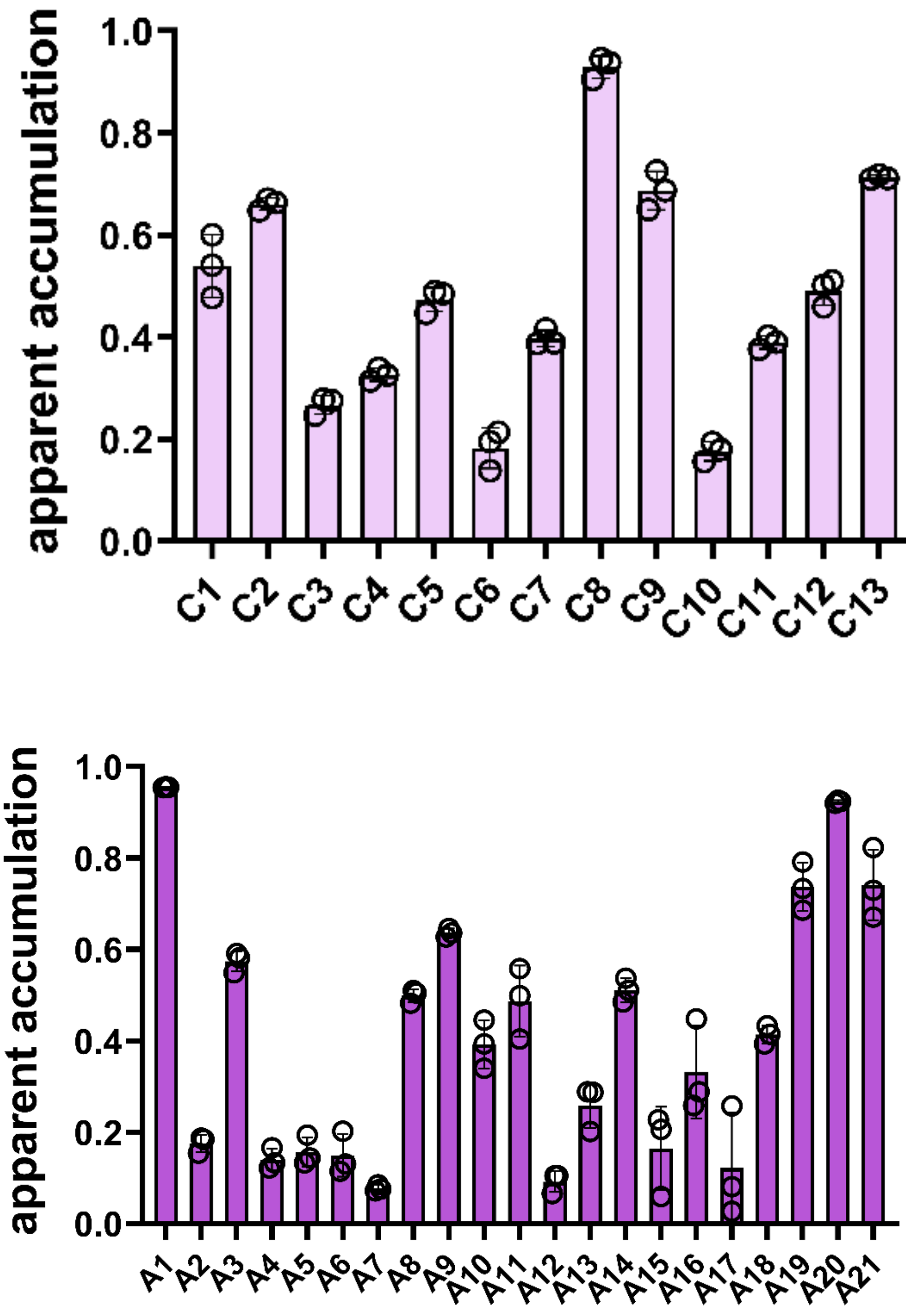


**Figure S1a.** Measurement of apparent accumulation of cationic (top) and amphipathic cell penetrating peptides (bottom) (CPPs) using Mammalian CHAMP assay. Cell expressing HaloTag were treated with 10 μM of DBCOcl for 15 min, pulsed with 50 μM of azide-tagged CPPs for 4 h, chased with 50 μM TMRaz for 15 min. Data are represented as mean ± SD (n=3).

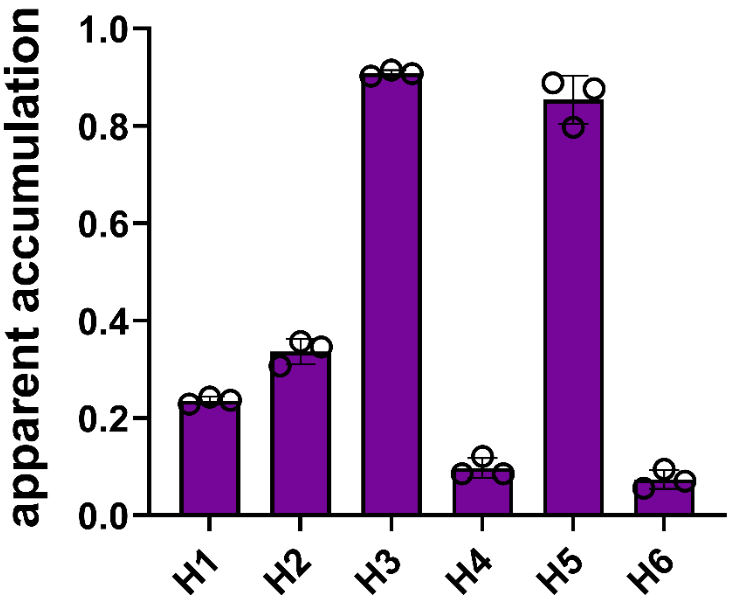


**Figure. S1b:** Measurement of apparent accumulation of hydrophobic cell penetrating peptides (CPPs) using Mammalian CHAMP assay. Cell expressing HaloTag were treated with 10 μM of DBCOcl for 15 min, pulsed with 50 μM of azide-tagged CPPs for 4 h, chased with 50 μM TMRaz for 15 min. Data are represented as mean ± SD (n=3).

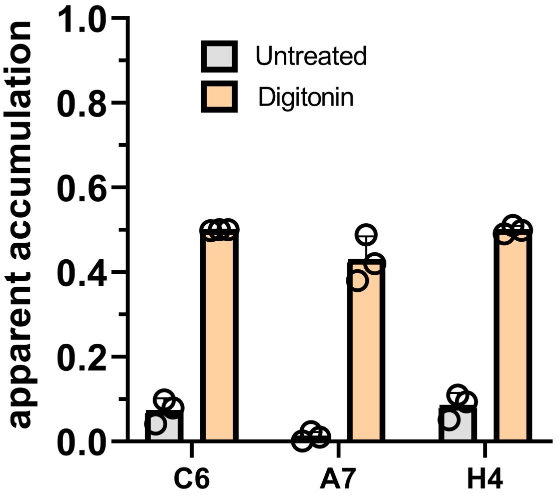


**Figure S2:** Digitonin permeabilization uptake of CPPs. Apparent accumulation of poor accumulator peptides with HT cells treated and untreated with digitonin. HT Cells were co-incubated with 40 μg/mL of digitonin for 2 h. Data are represented as mean ± SD (n=3).

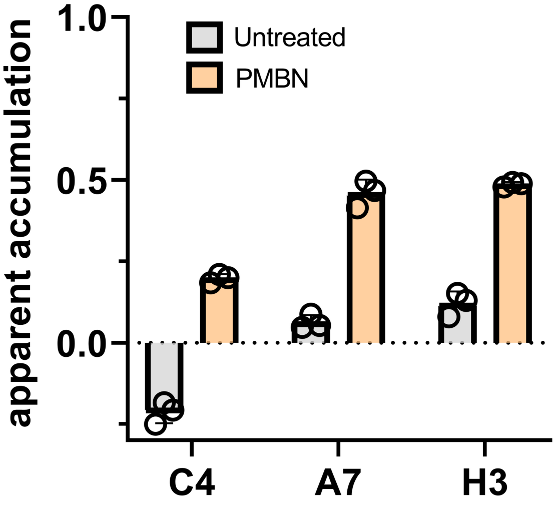


**Figure S3:** Outer Membrane Permeabilization using PMBN. Apparent accumulation of poor accumulator peptides with *E.coli* cells treated and untreated with PMBN. Cells were co-incubated with 10 μg/mL of PMBN for 1 h. Data are represented as mean ± SD (n=3).

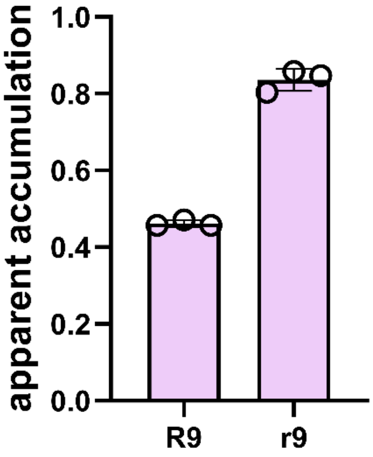


**Figure S4:** Effect of Stereochemistry on CPP accumulation in *E.coli.* Apparent accumulation of L-nonaarginine (R9) and its D-enantiomer (r9) in HaloTag-expressing *E. coli* measured by CHAMP. Data are represented as mean ± SD (n=3).

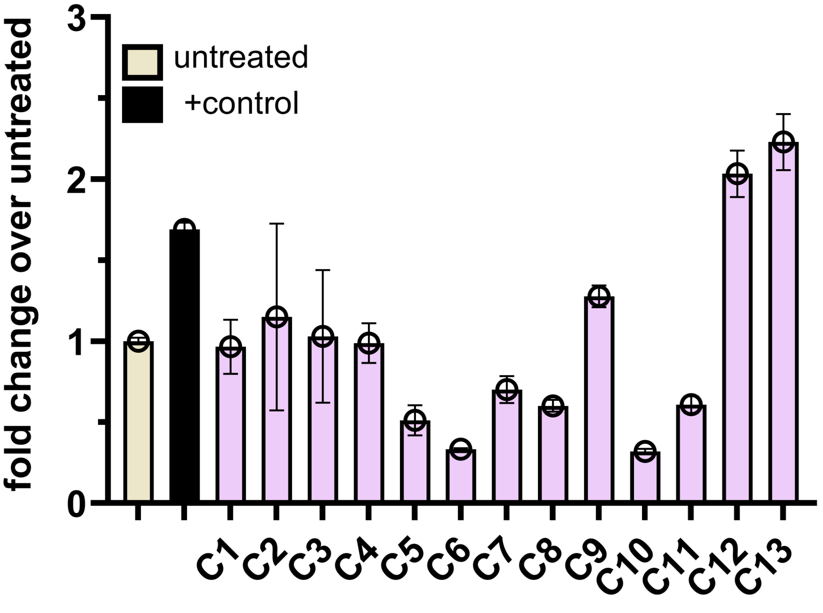


**Figure S5a:** Membrane integrity of *E. coli* after CPP treatment measured by propidium iodide (PI) uptake. PI fluorescence of *E. coli* treated with cationic CPPs(C1–C13). Data are represented as mean ± SD (n=3).

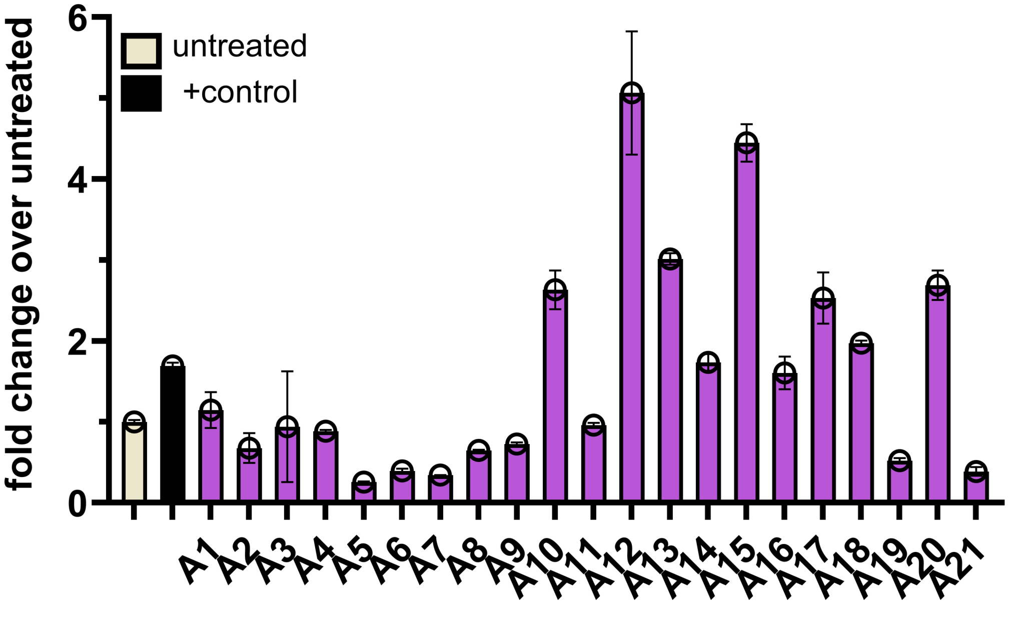


**Figure S5b:** Membrane integrity of *E. coli* after CPP treatment measured by propidium iodide (PI) uptake. PI fluorescence of *E. coli* treated with amphipathic CPPs (A1–A21). Data are represented as mean ± SD (n=3).

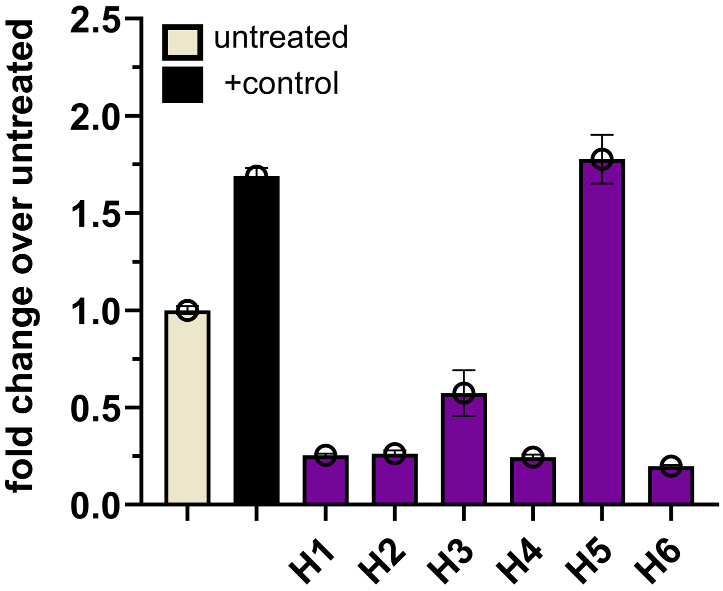


**Figure S5c:** Membrane integrity of *E. coli* after CPP treatment measured by propidium iodide (PI) uptake. PI fluorescence of *E. coli* treated with hydrophobic (H1–H6). Data are represented as mean ± SD (n=3).

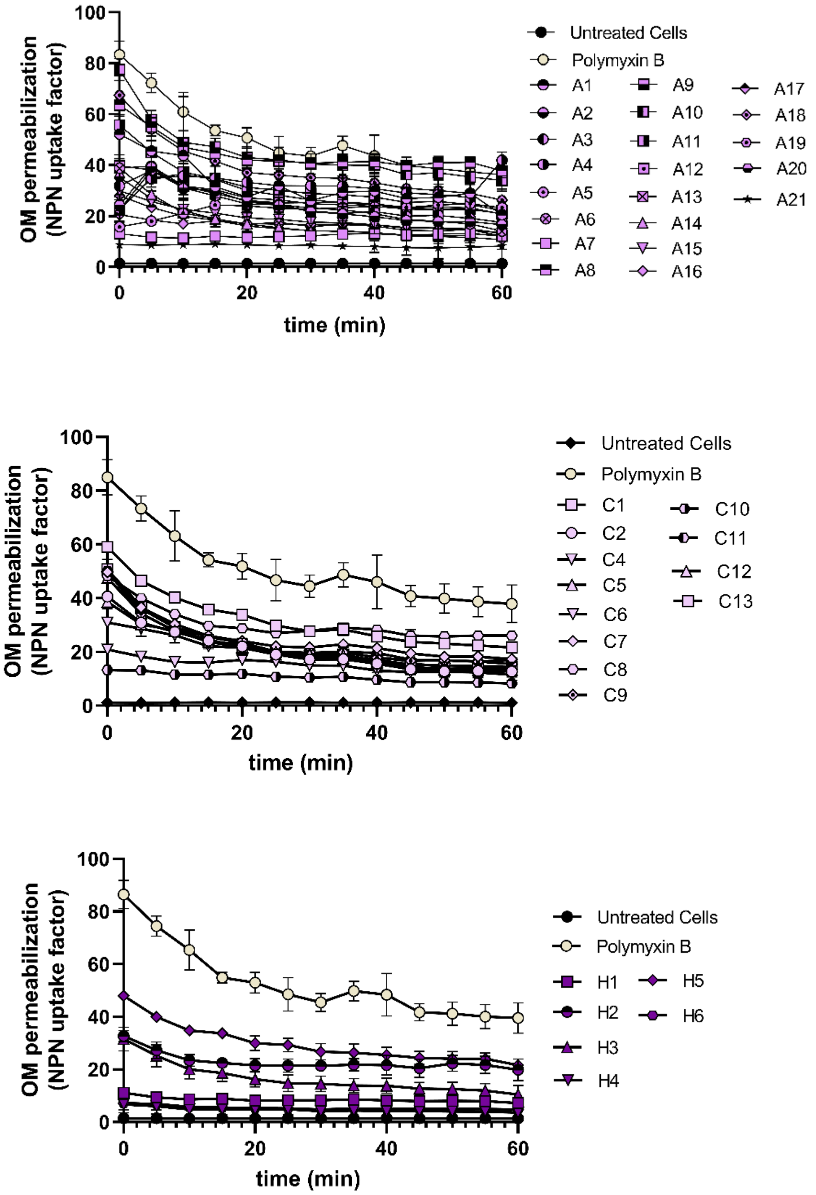


**Figure S6**: OM permeabilization of *E. coli* by CPPs measured by NPN uptake. The OM permeabilization (NPN uptake factor) of *E. coli* was measured over 60 min after treatment with amphipathic (A1–A21, top), cationic (C1–C13, middle) and hydrophobic (H1–H6), bottom) peptides at 50 μM. Polymyxin B was the positive control and untreated cells negative control.

#### Materials

The following materials were obtained from commercial sources: Cell culture media DMEM (11995065), Fetal Bovine Serum (A52568-01), Penicillin-Streptomycin (#P4333), Puromycin Dihydrochloride (#P4512) from Sigma-Aldrich. TAMRA-azide 6-isomer (TMRaz) (C8130), Fluorescein azide,6-isomer (D5130) from Lumiprobe. For the synthesis of peptides, Rink amide resin (10619) from ChemImpex and all other amino acids from Ambeed.

### Experimental Methods

#### Mammalian cell culture

HaloTag-expressing HeLa (HT HeLa) cells were cultured in Dulbecco's Modified Eagle Medium (DMEM) supplemented with 10% fetal bovine serum (FBS), 1% penicillin-streptomycin, and 1 μg/mL puromycin. Cells were maintained at 37°C with 5% $CO_2$. HaloTag HeLa cells were passaged using trypsin upon reaching 80–90% confluency and resuspended in DMEM. Cells were seeded in 96-well plates at a density of $3 \times 10^4$ cells per well, incubated for 24 h at 37°C with 5% $CO_2$.

#### Mammalian CHAMP Assay

$3 \times 10^4$ HeLa cells were seeded into 96-well plates one day before the assay, reaching 80–90% confluency. Following overnight incubation at 37°C, cells were washed three times with PBS and treated with 10 μM DBCOcl in media for 15 min. This was followed by a pulse with 50 μM of azide-tagged compounds for 1 h at 37°C. Concentration and time of incubation were adjusted based on the experiment requirements. Following the pulse step, chase step with 50 μM TMRaz in media for 15 min. Cells were washed three times with PBS between each step. After staining, cells were washed again, trypsinized , fixed with 4% formaldehyde, and analyzed using flow cytometry on the Attune NxT Acoustic Focusing Cytometer (Invitrogen).[1]

#### Digitonin Assay

Digitonin (Sigma, Part No. D141-100MG) was used for the membrane permeabilization assay in mammalian cells. For this assay, 40 μg/mL digitonin was co-incubated with peptides for 2 h following 15 min of pulse with 10 μM DBCOcl. This was followed by a chase step with 50 μM TMRaz for 15 min. Cells were then trypsinized, fixed with 4% formaldehyde, and analyzed using flow cytometry.

**MTT cell viability Assay**

HT HeLa cells were seeded into 96-well plates one day prior to the assay to achieve 80–90% confluency. After overnight incubation at 37°C, cells were washed three times with PBS and treated with 50 μM peptides for 2 h at 37 °C. Subsequently, 0.45 mg/mL MTT solution in media was added, and cells were incubated for 2 h at 37 °C. Following incubation, the plates were centrifuged, the supernatant was removed, and the resulting formazan crystals were dissolved in 100 μL of DMSO. Absorbance was measured at 570 nm using a plate reader.

**Expression of HaloTag protein and bacterial cell culture**

For expression of the HaloTag protein in WT *E. coli*, (smooth, ATCC25922), the expression plasmid Halo_pQE-30 was transformed into *Escherichia coli* and grown on an LB/agar plate with ampicillin (100□μg/mL) at 37 °C overnight. Colonies were picked and grown first in 4 mL LB broth with ampicillin at 37 °C overnight. A growth tube with 4 mL of LB media was then inoculated with the overnight culture at a ratio of 1:100 in the presence of 100 μg/mL ampicillin and cells were grown at 37 °C for 1.5 h, or until the O.D. at 600 nm reached 0.2 when cultures were induced with 1 mM Isopropyl β-d-1-thiogalactopyranoside (IPTG) at 37 °C for 2 h in a shaker incubator. Cells were then pelleted at 4000 rpm for 3 min. The pellet was washed three times with 1X PBS pH 7.4, following which cells were resuspended in 35 mL of 1X PBS for the assay.

**Bacterial CHAMP Assay**

Following IPTG-expression, 4 mL of 50 μM **DBCOcl** would be added to the *E. coli* cells and were incubated for 30 min at 37 °C. 100 μL of the cell suspension was plated in triplicate on a 96 well plates and subsequently washed 2x with PBS. Then, cells were resuspended in 50 μL of PBS and 50 μL of a 100 μM molecule stock, obtained from a 10 mM parent stock, was added to each of the wells, which was then covered and incubated for 1 hat 37 °C. In the case of time and concentration studies, similar protocol was altered to account for these changes. Control wells, on the same plate, were incubated with 100 μL of PBS for the same duration. Cells were then pelleted in a plate of holder centrifuge at 4000 RPM for 3 min, and the supernatant was discarded. After washing 2x with PBS, the cells were immediately resuspended in 50 μL of PBS and to that was added 50 μL of 100 μM TMRaz and incubated for 1 h at 37 °C. Cells were pelleted once again, washed 1x with PBS, and fixed with 4% formaldehyde for 30 min. Following formaldehyde fixation, the cells were washed 1X with PBS and resuspended in 200 μL of PBS for analysis by flow cytometry on the Attune NxT Acoustic Focusing Cytometer (Invitrogen).[2]

**Bacterial assay with PMBN**

DBCO-labeled smooth *E. coli* was prepared as stated earlier and washed 2x with PBS. Cells were resuspended in 50 μL of a 100 μM molecule stock and immediately 50 μL PBS supplemented with 20 μg/mL PMBN and was added to reach a final concentration of 10 μg/mL PMBN. Cells were incubated for 1 h at 37 °C in the shaker incubator. Cells were then pelleted in a plate holder centrifuge at 4000 RPM for 3 min, and the supernatant was discarded. After washing 2x with PBS, the cells were immediately resuspended in 50 μL of PBS and to that was added 50 μL of 100 μM TMRaz and incubated for 1 h at 37 °C. Cells were pelleted once again, washed 1x with PBS, and fixed with 4% formaldehyde for 30 min. Following formaldehyde fixation, the cells were washed 1X with PBS and resuspended in 200 μL of PBS for analysis by flow cytometry on the Attune NxT Acoustic Focusing Cytometer (Invitrogen).

**Membrane Integrity Assay via SYTO9**

To measure if the cell membrane is comprised due to cell death or damage by the peptides, DBCO-labeled smooth *E. coli* was prepared as stated earlier and washed 2x with PBS. 100 μL of the cell suspension was plated in triplicate on a 96 well plates and subsequently washed 2x with PBS. Then, cells were resuspended in 50 µL of PBS and 50 μL of a 20 μM molecule stock, obtained from a 10 mM parent stock, was added to each of the wells, which was then covered and incubated for 1 hat 37 °C. Control wells, on the same plate, were incubated with 100 μL of PBS for the same duration. Cells were then pelleted in a plate holder centrifuge at 4000 RPM for 3 min and the supernatant was discarded. After washing 2x with PBS, the cells were immediately resuspended in 50 μL of PBS and to that was added 50 μL of 100 μM SYTO9 and incubated for 1 hat 37 °C. Cells were pelleted once again, washed 1x with PBS, and resuspended in 200 μL of PBS for analysis by flow cytometry on the Attune NxT Acoustic Focusing Cytometer (Invitrogen)

**Propidium Iodide Assay**

To measure if the cell membrane is comprised due to cell death or damage by the peptides, DBCO-labeled smooth *E. coli* was prepared as stated earlier and washed 2x with PBS. 100 μL of the cell suspension was plated in triplicate on a 96 well plates and subsequently washed 2x with PBS. Then, cells were resuspended in 50 µL of PBS and 50 μL of a 20 μM molecule stock, obtained from a 10 mM parent stock, was added to each of the wells, which was then covered and incubated for 1 hat 37 °C. Control wells, on the same plate, were incubated with 100 μL of PBS for the same duration. Cells were then pelleted in a plate holder centrifuge at 4000 RPM for 3 min and the supernatant was discarded. After washing 2x with PBS, the cells were immediately resuspended in 50 μL of PBS and to that was added 50 μL of 100 μM PI and incubated for 1 hat 37 °C.

Cells were pelleted once again, washed 1x with PBS, and resuspended in 200 μL of PBS for analysis by flow cytometry on the Attune NxT Acoustic Focusing Cytometer (Invitrogen).

##### NPN Uptake Assay

For outer membrane permeability, we tested using an NPN uptake assay. 50 μL of NPN (Millipore-Sigma), at a final concentration of 10 μM in PBS, was added to the DBCO-labeled *E. coli* cells at stationary phase. 50 μL of given molecules were added to achieve a final concentration of 50 μM for a total volume of 100 μL per well. For positive control, 20 μL of Polymyxin B, to achieve a final concentration of 16 μg/mL was added to the cells. All samples were completed in triplicate. Fluorescence intensity, at 37 °C with shaking, was measured using an Agilent Biotek Synergy H1 well-plate reader using an excitation wavelength of 355 nm and an emission wavelength of 405 nm. Measurements were taken every 5 min for 60 min total, and OM permeabilization was calculated using the NPN uptake factor formula below:

(Fluorescence of sample with NPN – Fluorescence of sample without NPN)/ (Fluorescence of PBS with NPN – Fluorescence of PBS without NPN). Raw NPN fluorescence values were normalized using OD600nm.

#### Synthesis and Characterization of Test Molecules

##### Synthesis of peptides via solid phase peptide synthesis (SPPS)

Cell Penetrating Peptides were synthesized via solid phase peptide chemistry. To a 15 mL peptide synthesis vessel, an appropriate amount of resin was added. Rink amide resin (0.48 mmol/g loading capacity) was used to ensure C-terminal amidation upon cleavage. It was followed by the addition of 20% piperidine in *N, N*-Dimethylformamide (DMF, 15 mL), and then washed (3X) with *N, N* Dimethylformamide. After the last wash, 4 equiv. of amino acid was added along with 4 equiv. of ethyl cyanohydroxyiminoacetate (Oxyma) and 4 equiv. of *N, N'*-Diisopropylcarbodiimide (DIC). The remainder of the amino acids were coupled in the same manner. After coupling with the last amino acid, the azide coupling was done outside the automated peptide synthesizer. For azide coupling on the N-terminus, 4 equiv. of Oxyma, 4 equiv. of DIC and 4 equiv. of 2-azidoacetic acid were added and let it shake for 2 h at room temperature. Peptides were cleaved from the resin using a TFA/TIPS/$H_2O$ mixture (95:2.5:2.5, v/v/v) shaking at room temperature for 2 h. The solution was filtered and precipitated by the addition of cold diethyl ether to yield crude peptide.

**Characterization of test molecules**

Peptides were purified via reverse-phase high-performance liquid chromatography (RP-HPLC) equipped with Waters 1525 with 2489 UV/Visible Detector on a Phenomenex Luna 10 μm C8(2) 100 A (250 x 21.2 mm) column using gradient elution with $H_2O$/MeOH with 0.1% TFA at 10 mL/min. The HPLC fractions of the desired purified compounds were first concentrated under reduced pressure using a rotary evaporator. The final concentrated aqueous solutions were lyophilized to dryness using Labconco Freezone 4.5 L (-84 $^{o}$C) lyophilizer and verified to be >97% pure by RP-HPLC and MALDI-TOF mass spectrometry. Peptide identities were confirmed via high resolution electrospray ionization mass spectrometry (HRMS, ESI/MS).

#### Analytical HPLC Chromatogram and MALDI-TOF

## 2BP100

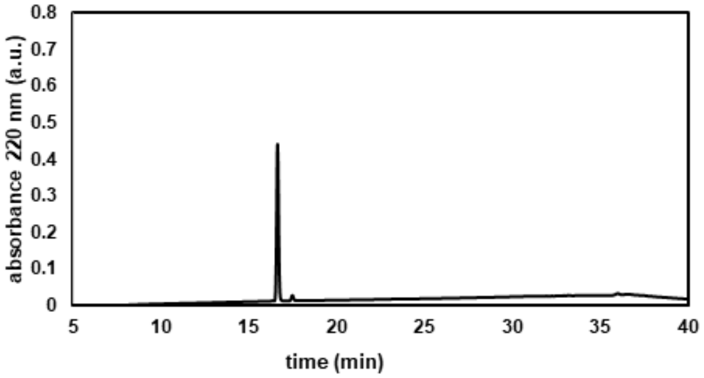


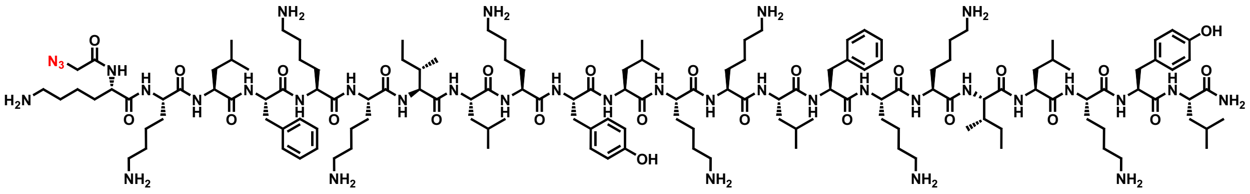

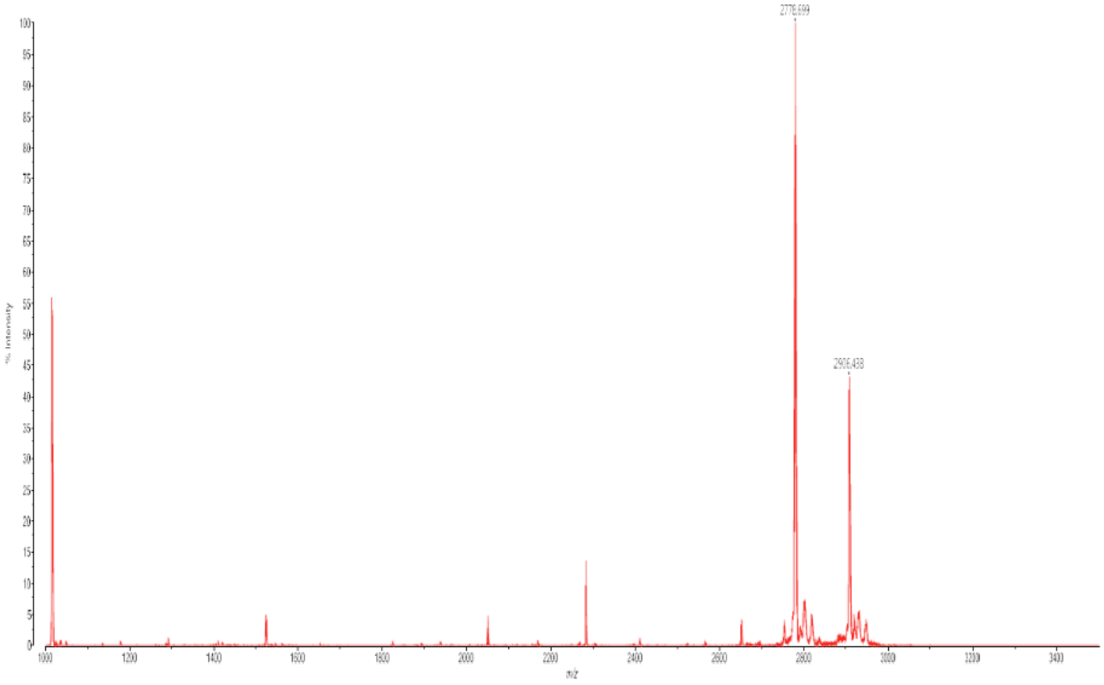

**Rev (34-50)**

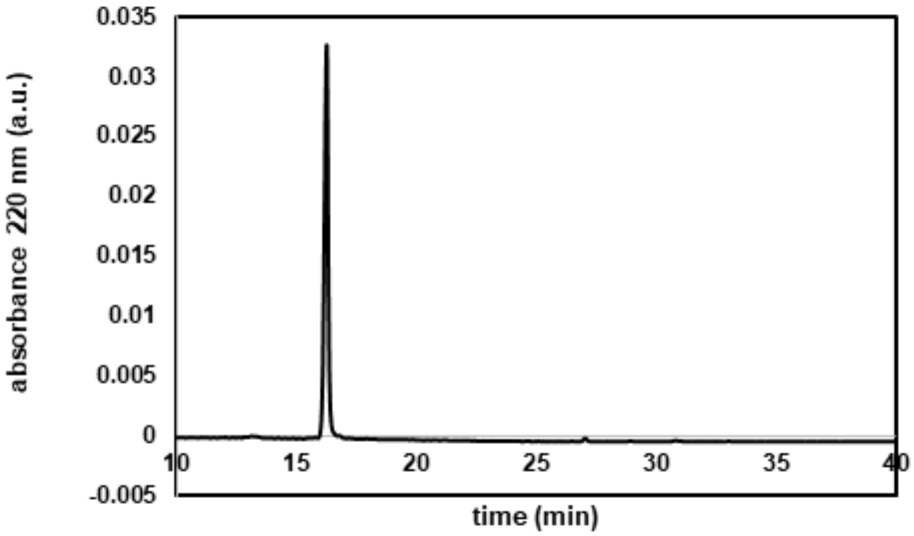

absorbance 220 nm (a.u.)
0.035
0.03
0.025
0.02
0.015
0.01
0.005
0
-0.005
10
15
20
25
30
35
40
time (min)


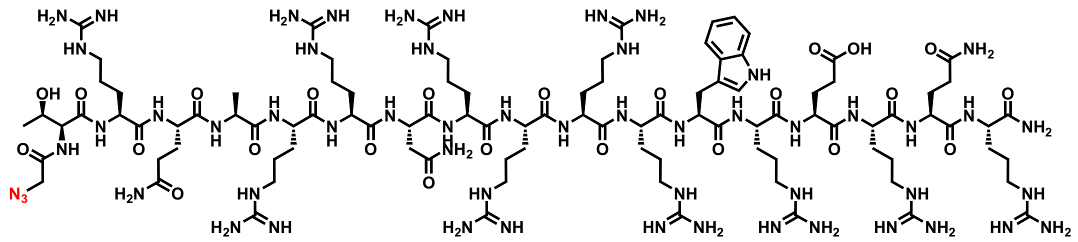

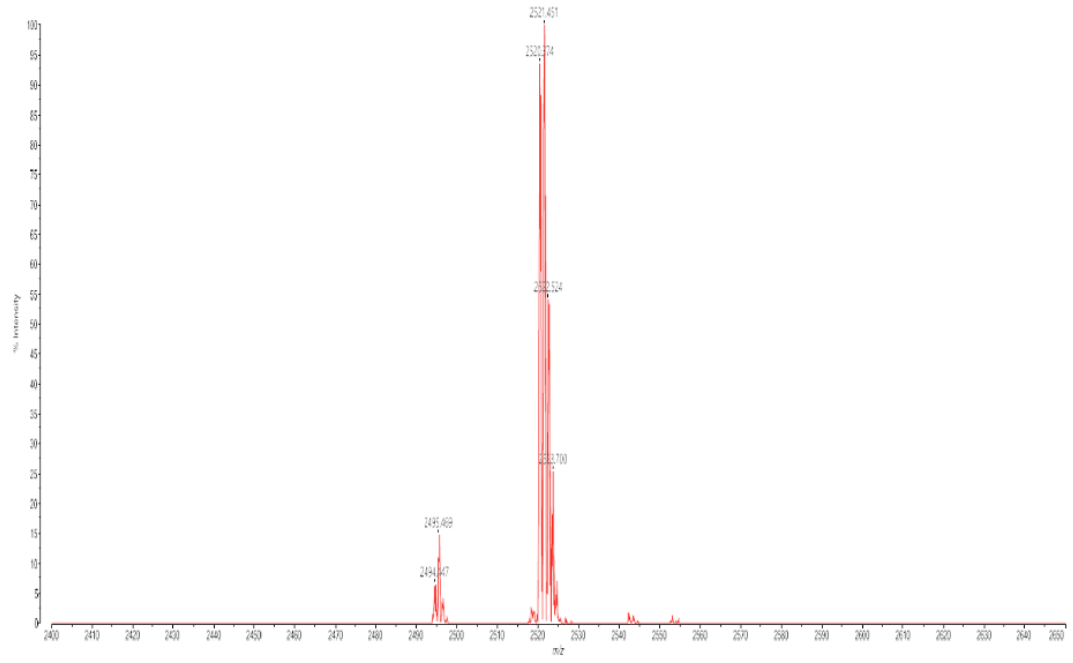

2521.451
2495.469

**r9W**

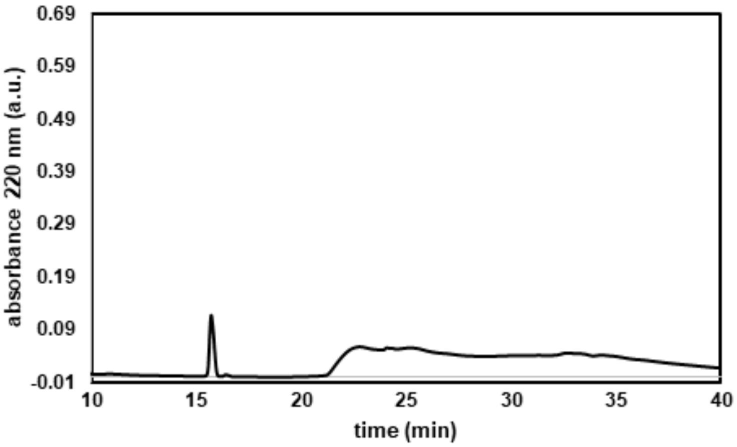

absorbance 220 nm (a.u.)
0.69
0.59
0.49
0.39
0.29
0.19
0.09
-0.01
10
15
20
25
30
35
40
time (min)


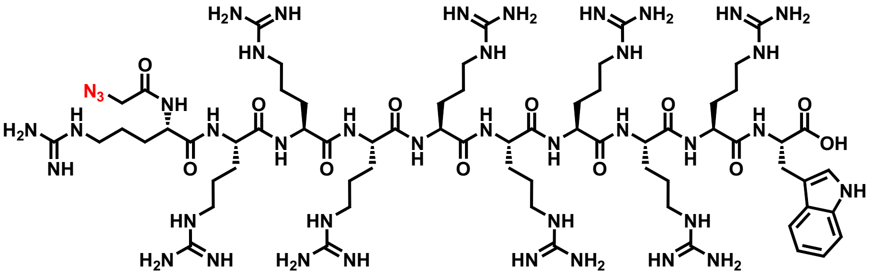

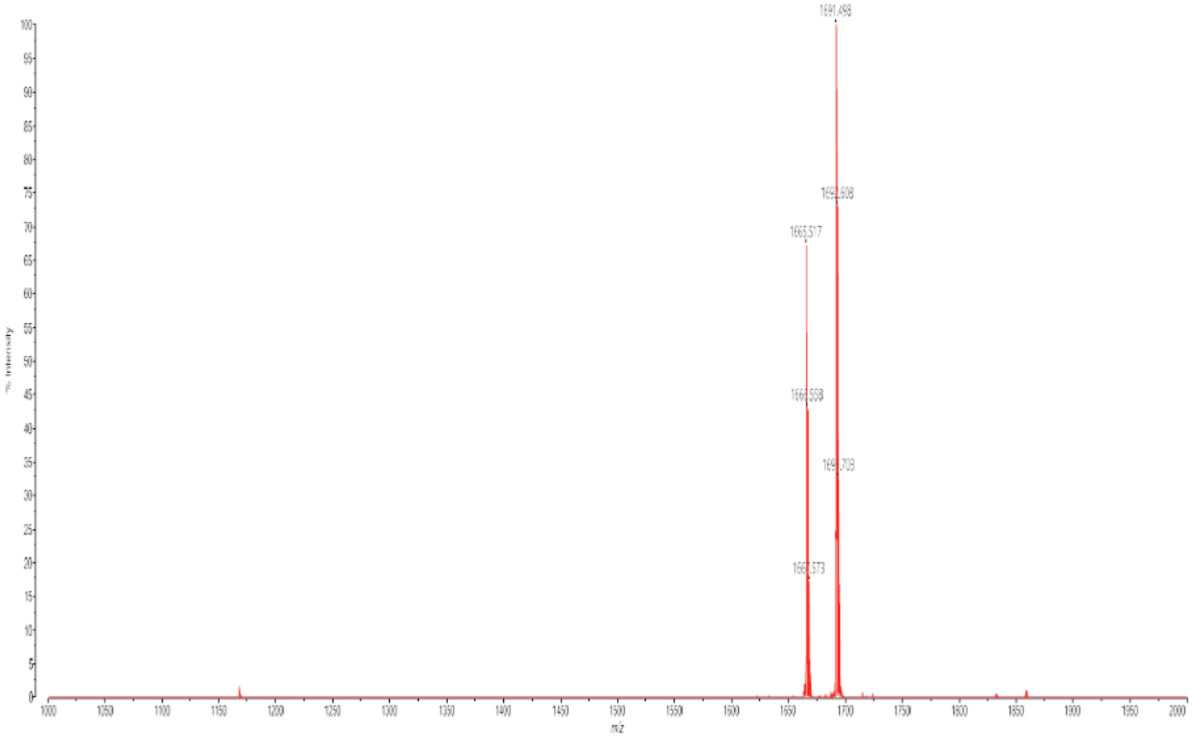

1691.498
1665.517
m/z

**(KH)9F**

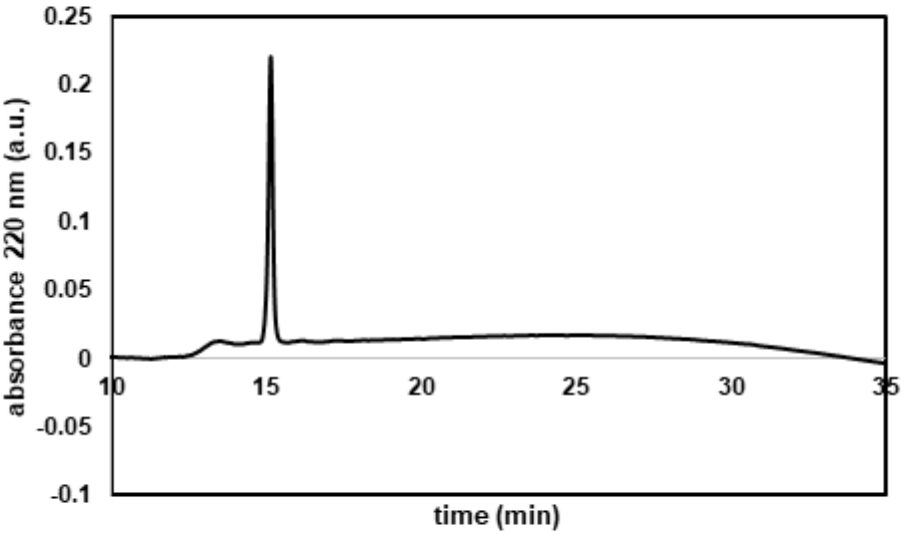
0.25
0.2
0.15
0.1
0.05
0
-0.05
-0.1
absorbance 220 nm (a.u.)
10
15
20
25
30
35
time (min)

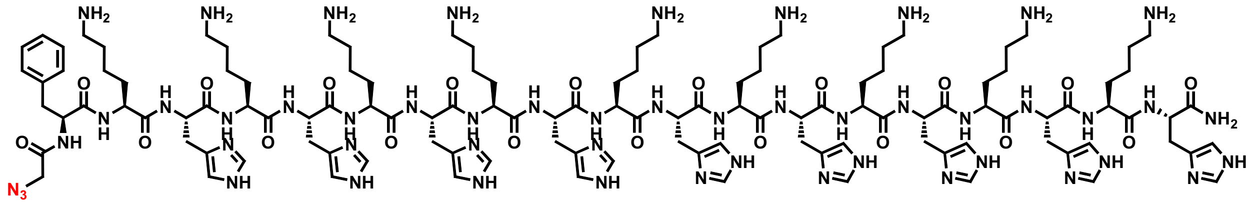

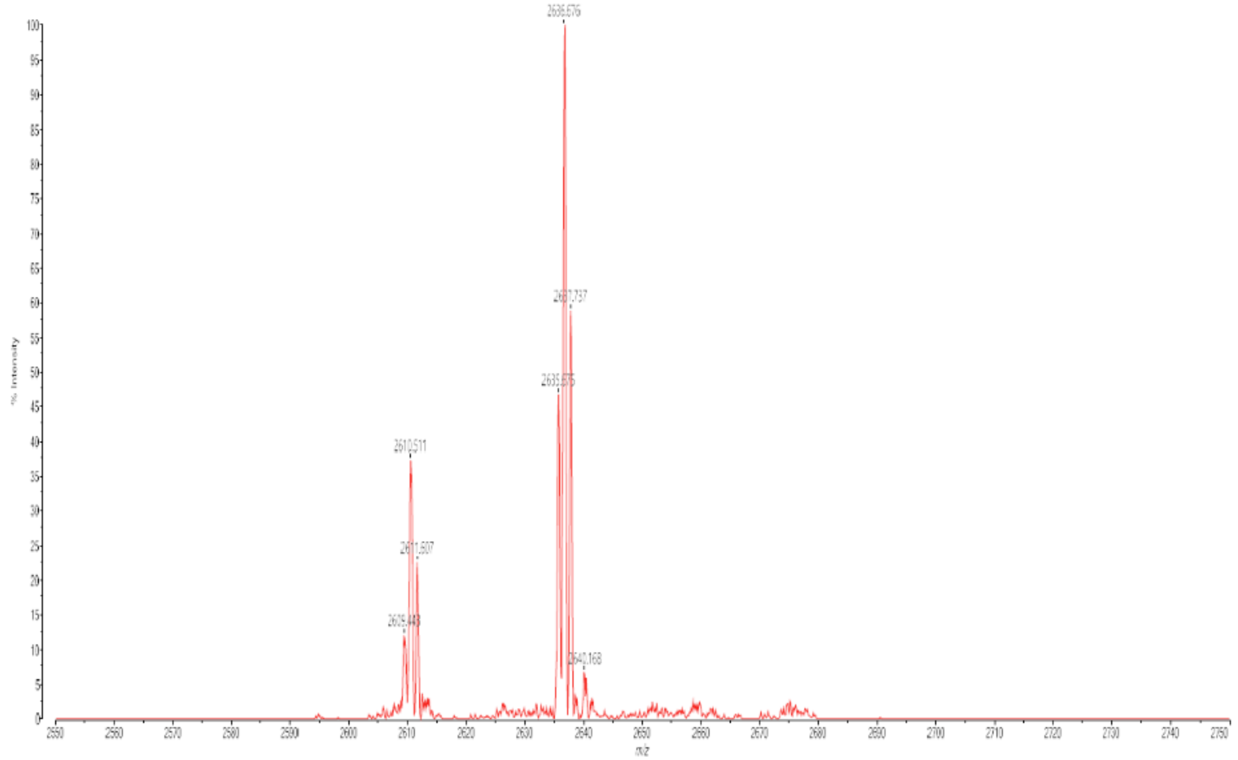
2636.676
2610.511
% Intensity
m/z

**K9F**

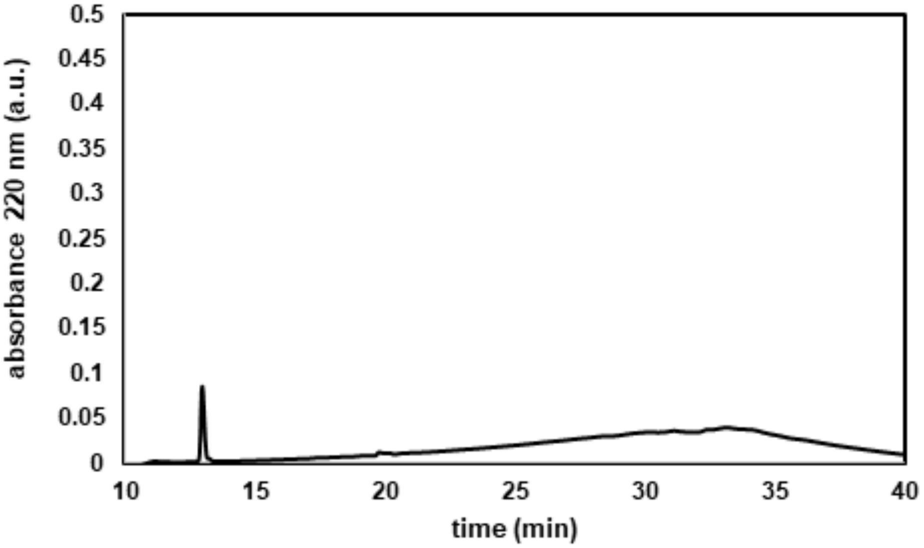

absorbance 220 nm (a.u.)
0.5
0.45
0.4
0.35
0.3
0.25
0.2
0.15
0.1
0.05
0
10
15
20
25
30
35
40
time (min)


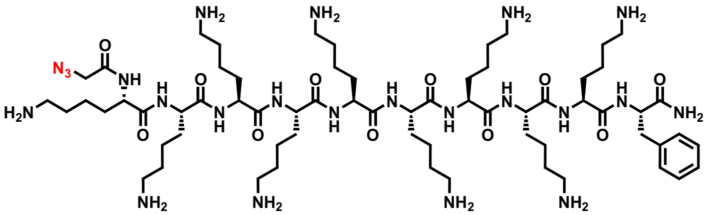

$N_3$
NH
$H_2N$
$NH_2$


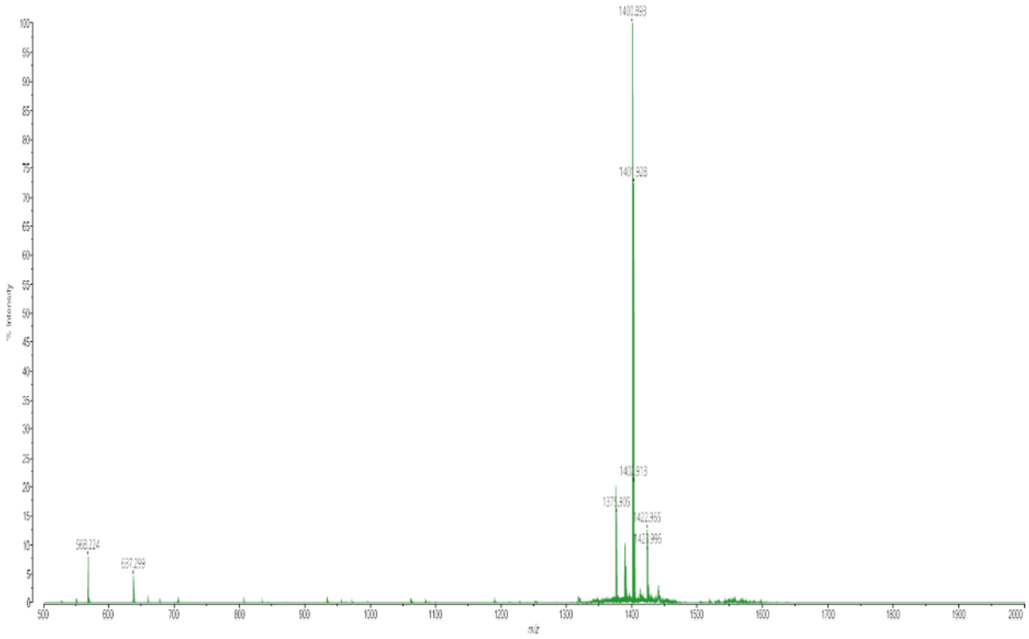

**DPV3**

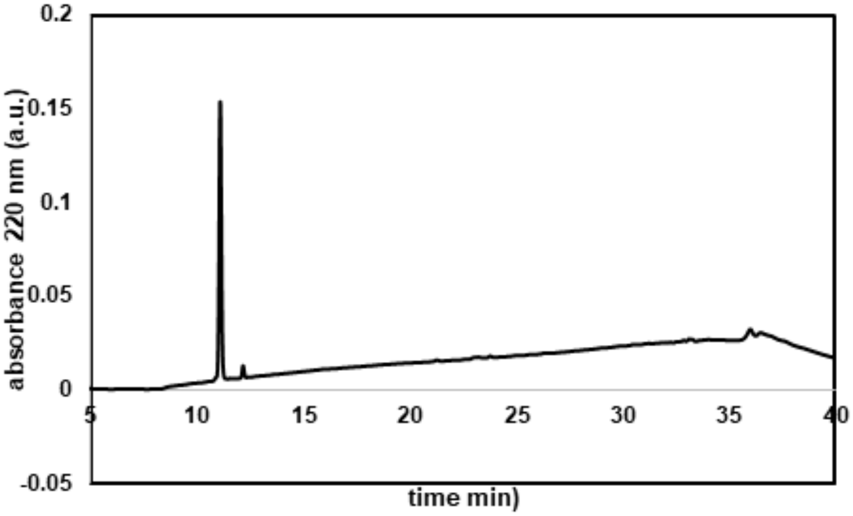

0.2
0.15
0.1
0.05
0
-0.05
absorbance 220 nm (a.u.)
5
10
15
20
25
30
35
40
time min)


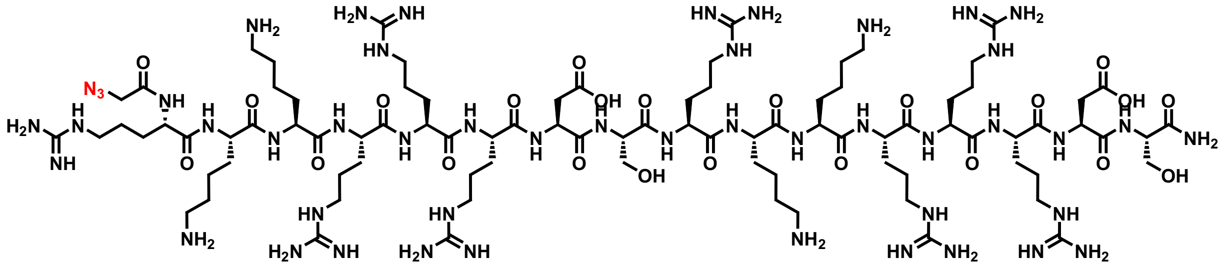

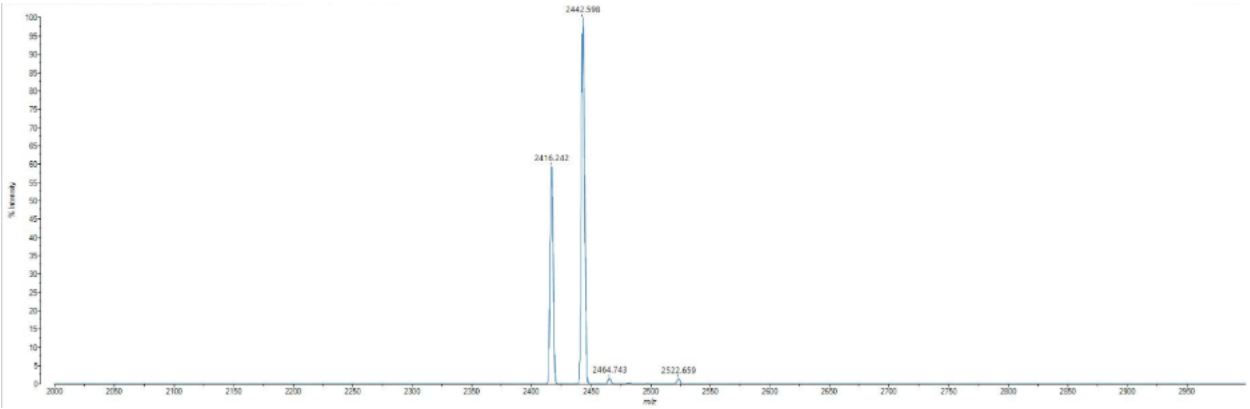

2442.598
2416.242
2464.743
2522.659

**6-Oct**

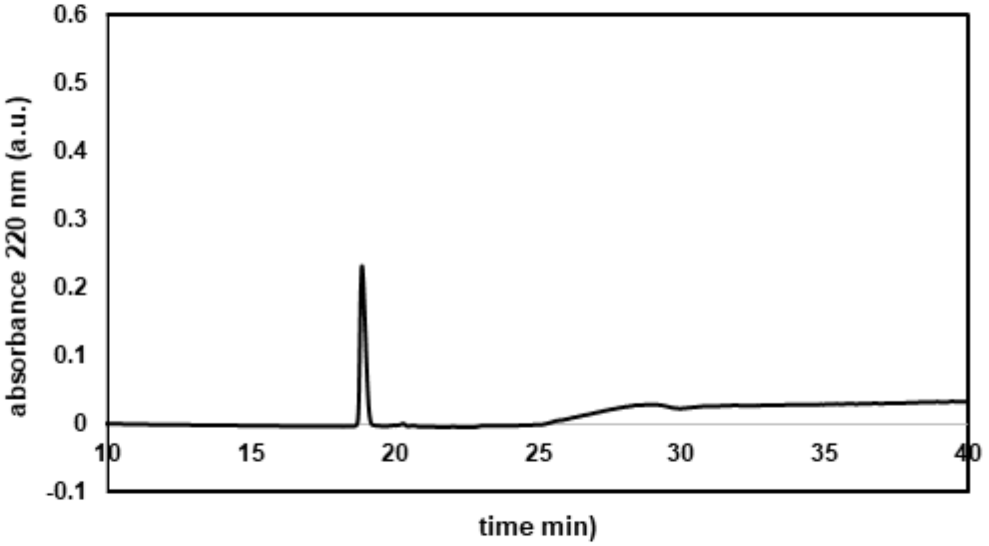


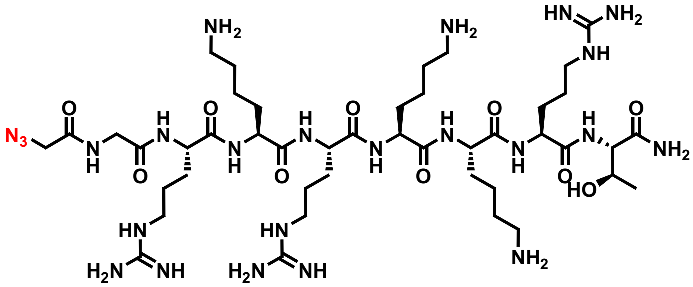

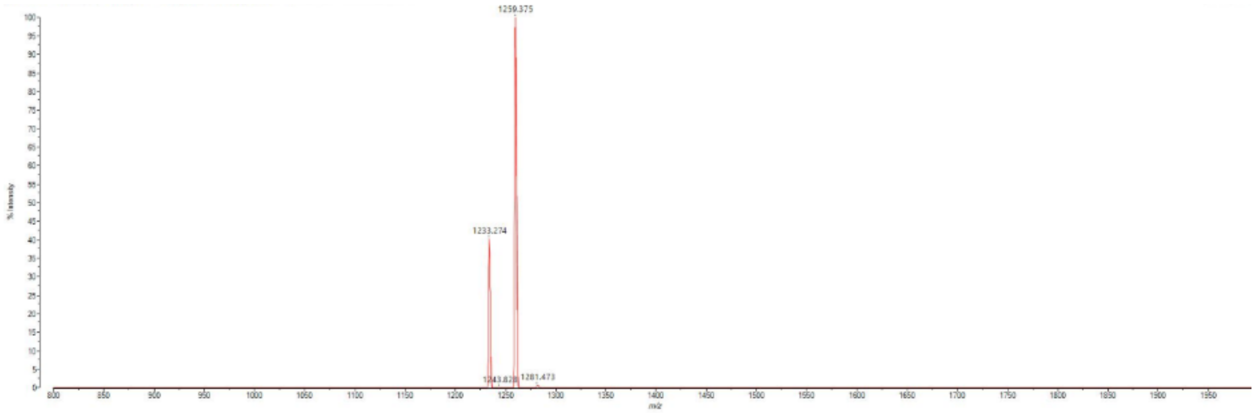

**Tat (49-57)**

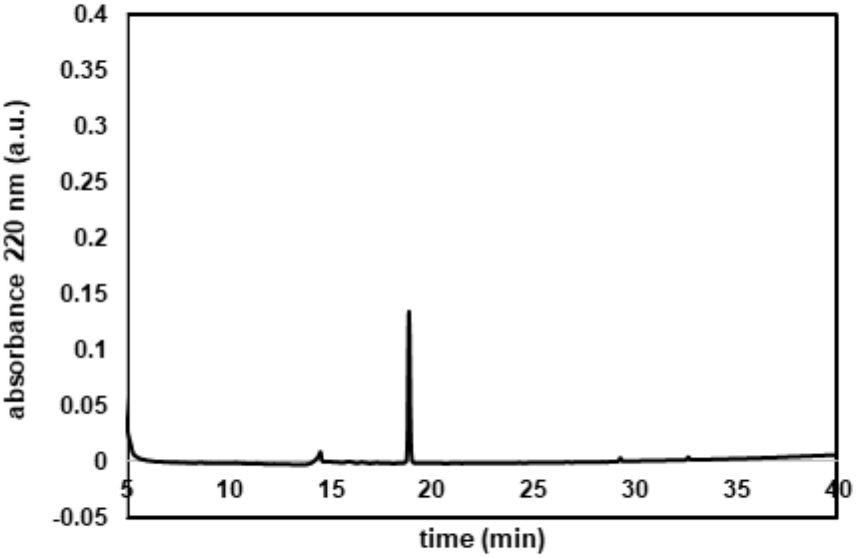
absorbance 220 nm (a.u.)
time (min)
0.4
0.35
0.3
0.25
0.2
0.15
0.1
0.05
0
-0.05
5
10
15
20
25
30
35
40

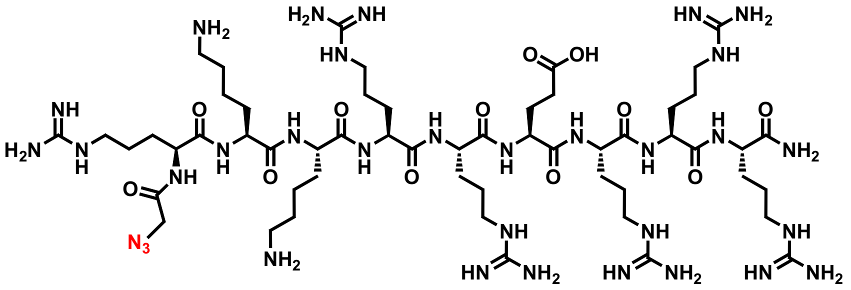
$N_3$
$NH_2$
$H_2N$
NH
HN
$NH_2$
O
OH
H
N

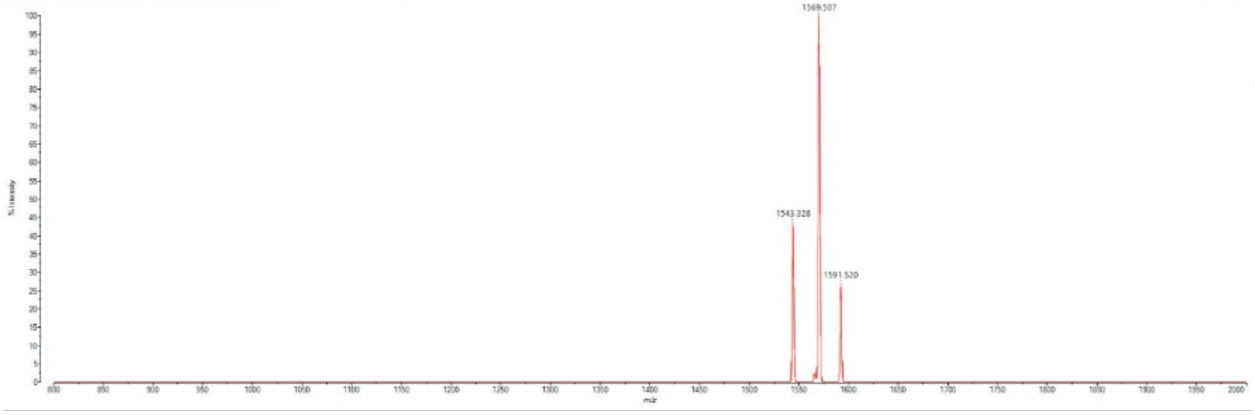
1569.507
1543.328
1591.520

**(KFF)3K**

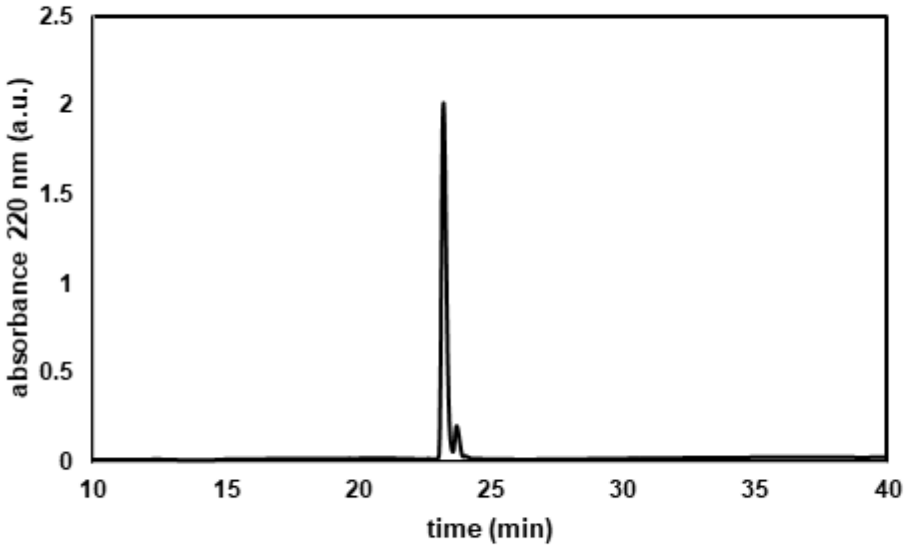

absorbance 220 nm (a.u.)
2.5
2
1.5
1
0.5
0
10
15
20
25
30
35
40
time (min)


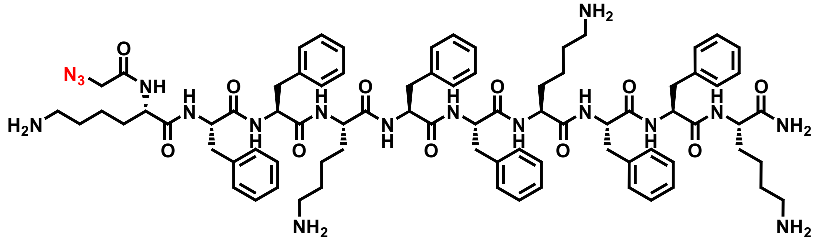

N3
NH
H2N
NH2
NH2
NH2
NH2


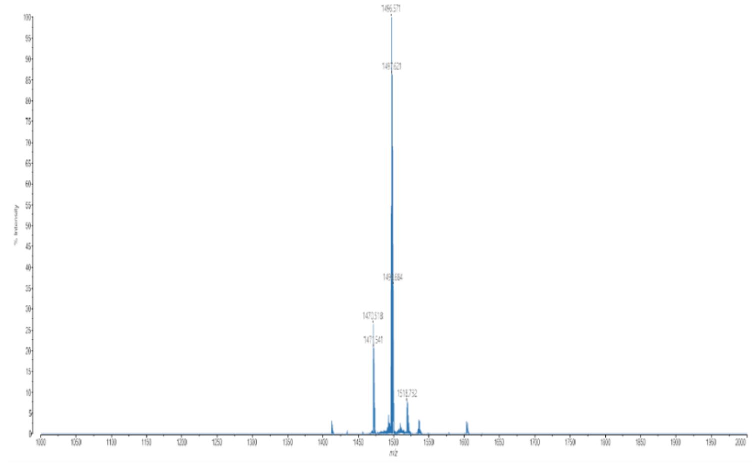

1496.571
1497.621
1470.518
1518.732
m/z

**RRRRR**

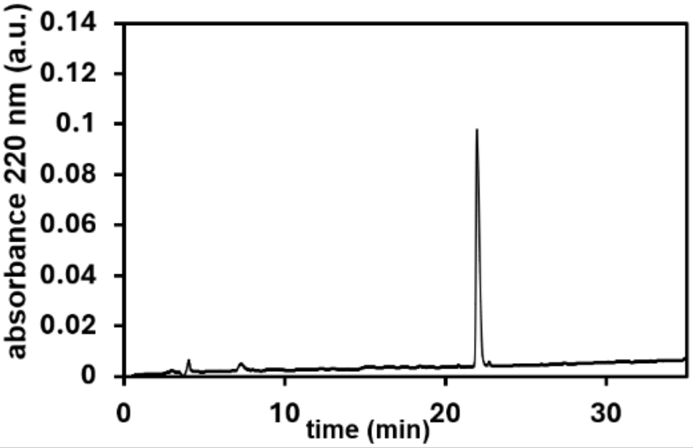


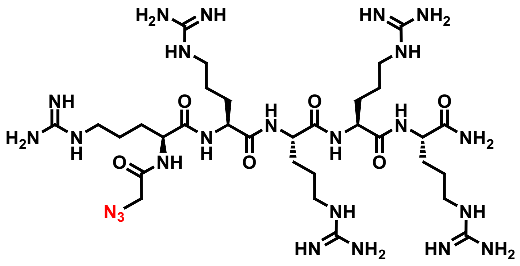


**QTOF High Resolution Mass Spectrum for R5az (m/z 880.5441for $[M+H]^+$ found 881.5522)**

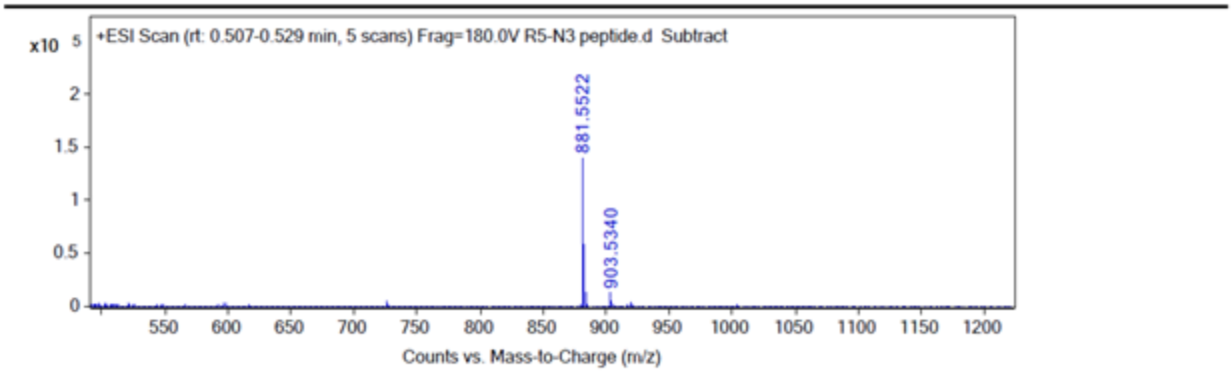

**RRRRRRR**

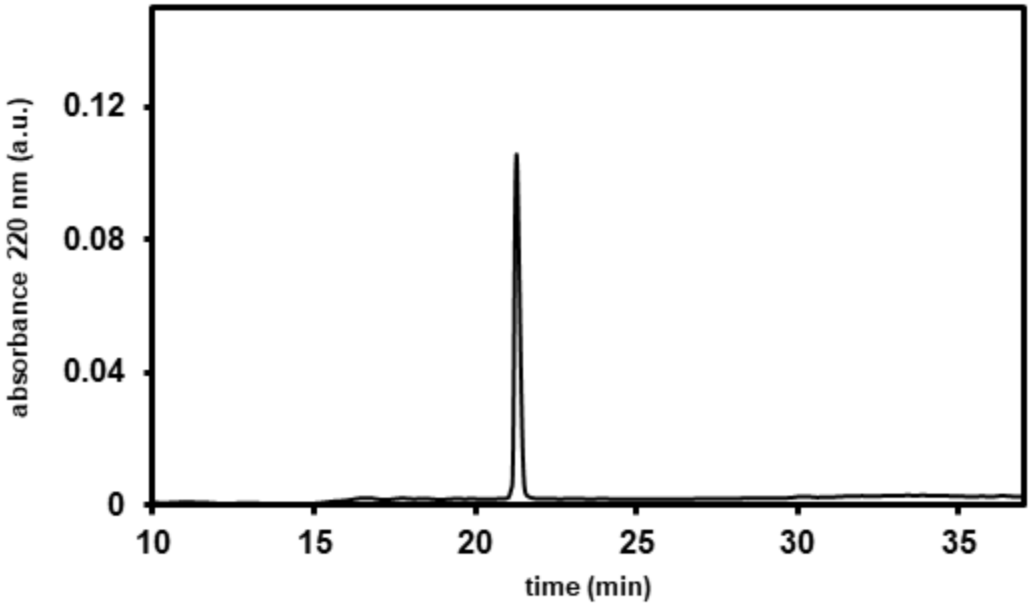


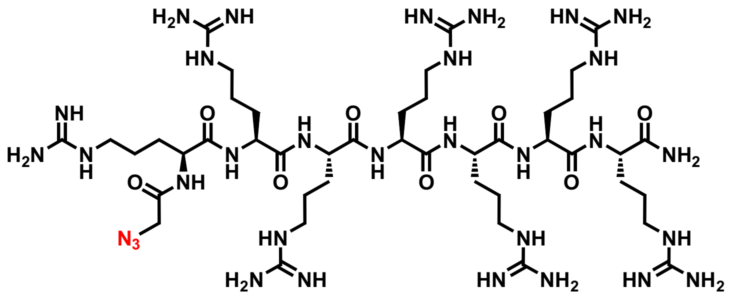

**QTOF High Resolution Mass Spectrum for R7az (m/z 1192.7463) for $[M+H]^+$ found 1193.7542)**

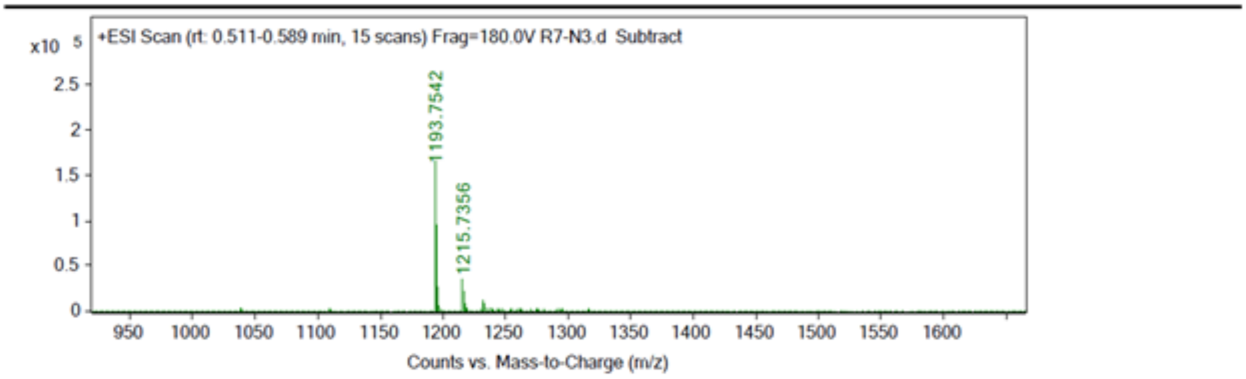

**RRRRRRRRR**

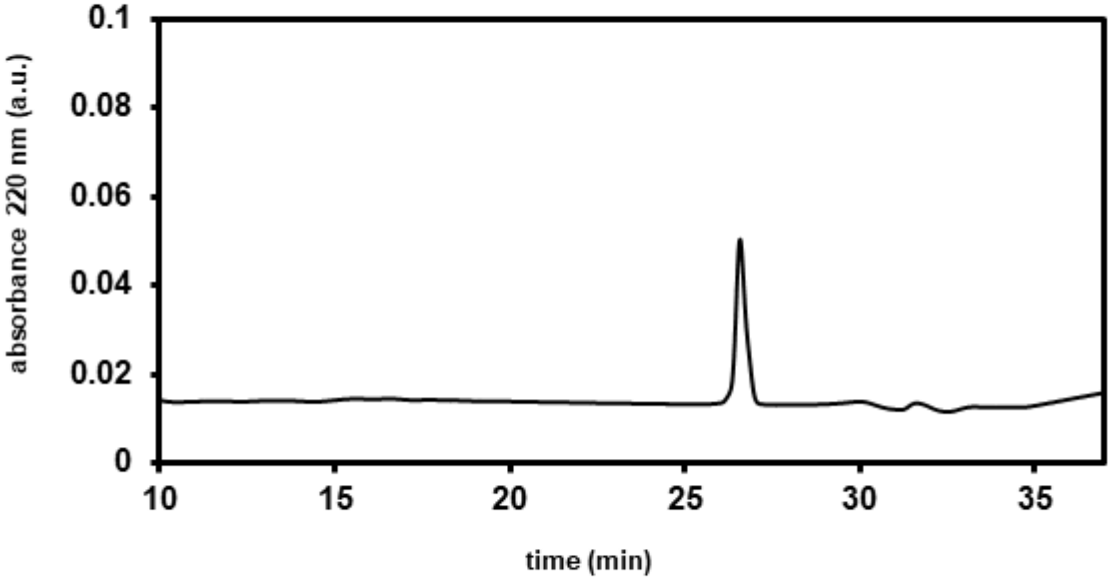


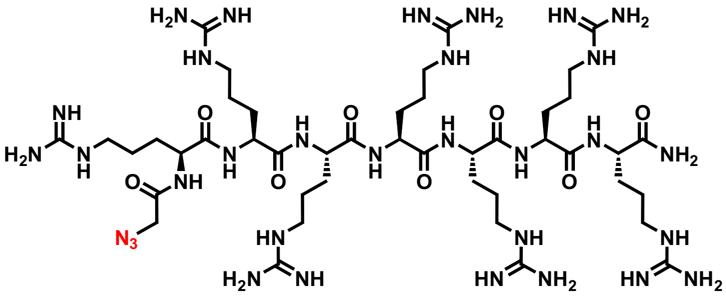

**QTOF High Resolution Mass Spectrum for R9az (m/z .1504.9485 for $[M+H]^+$ found 1505.9566)**

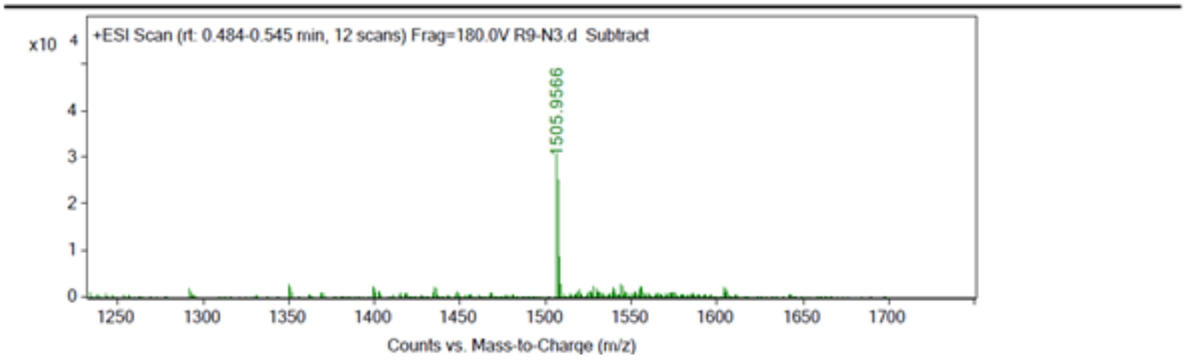

**RRRRRRRRRRR**

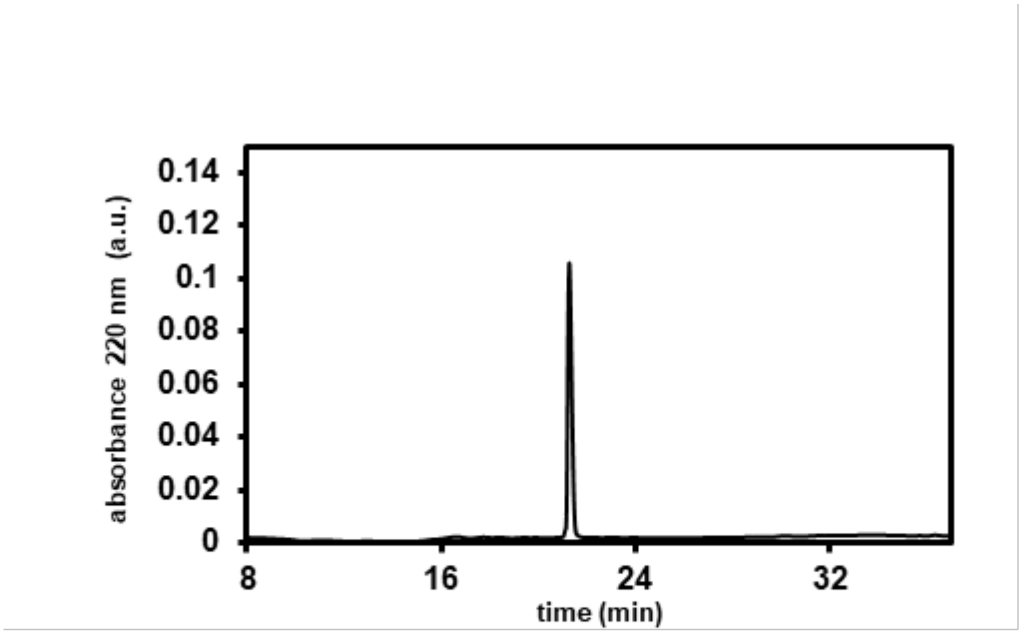


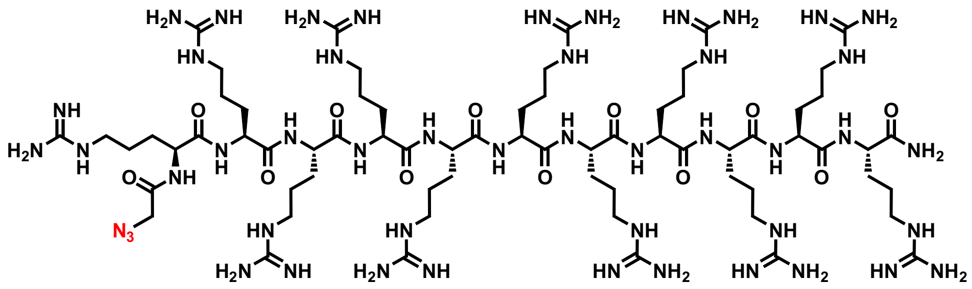

**QTOF High Resolution Mass Spectrum for R11az (m/z 1817.1501 for $[M+H]^+$ found 188 .5522)**

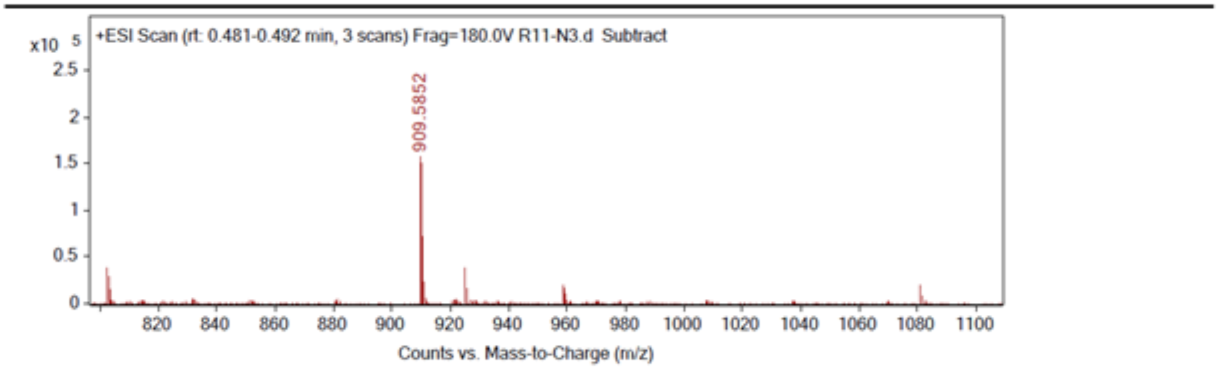

**BP100**

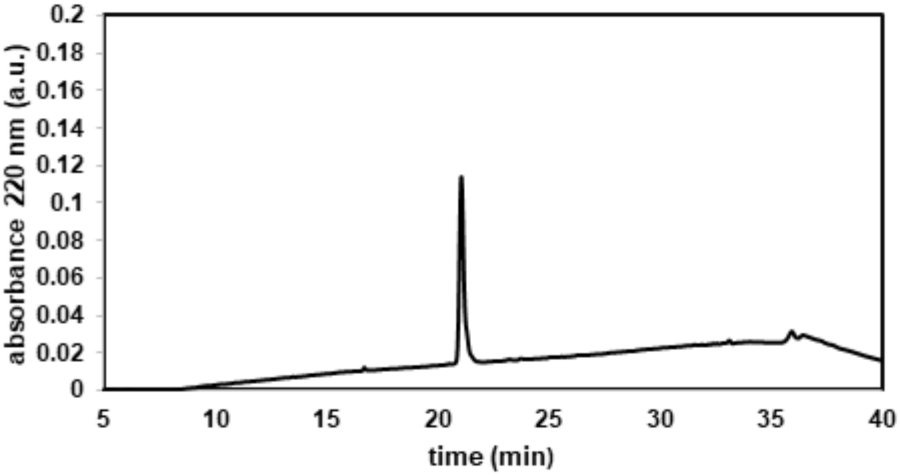

0.2
0.18
0.16
0.14
0.12
0.1
0.08
0.06
0.04
0.02
0
absorbance 220 nm (a.u.)
5
10
15
20
25
30
35
40
time (min)


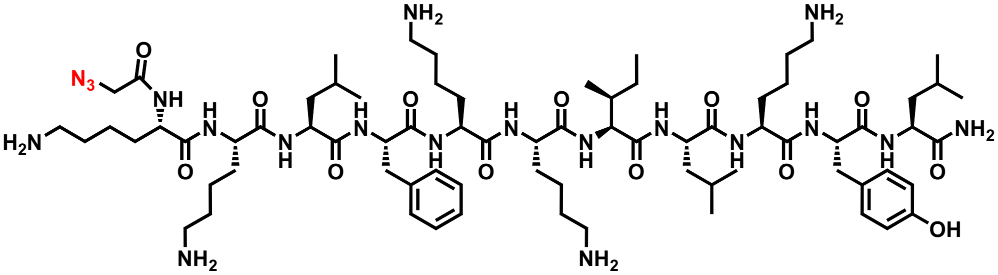

N3
O
NH
H2N
NH2
OH


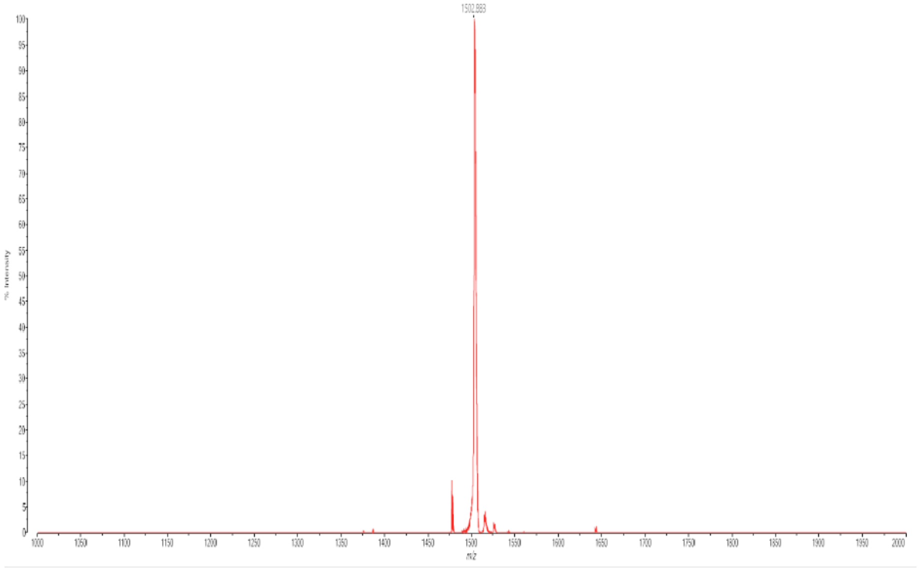

**Sc18**

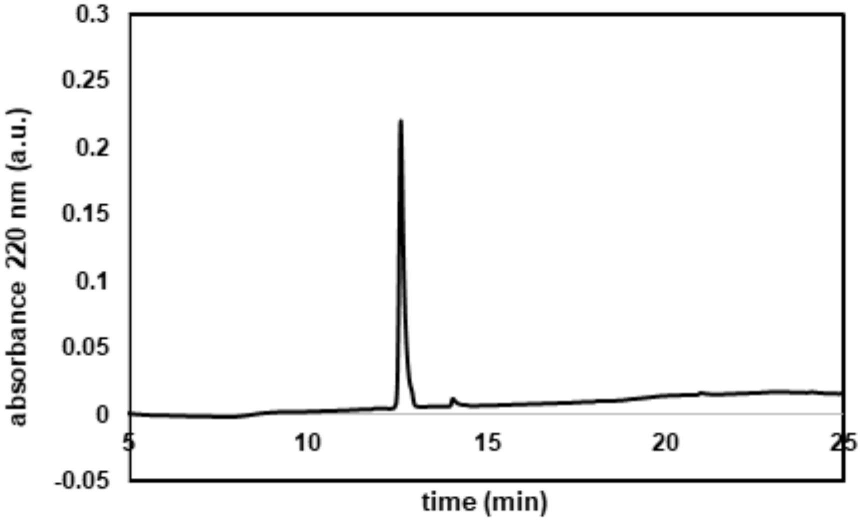

0.3
0.25
0.2
0.15
0.1
0.05
0
-0.05
absorbance 220 nm (a.u.)
5
10
15
20
25
time (min)


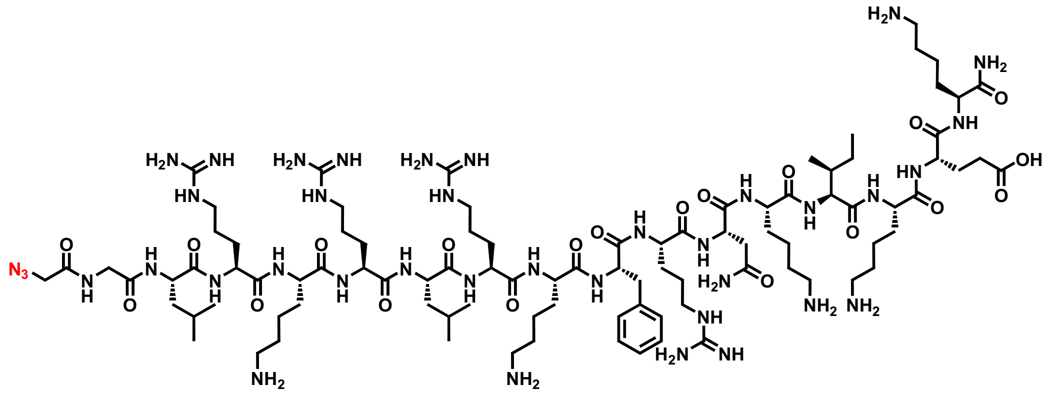

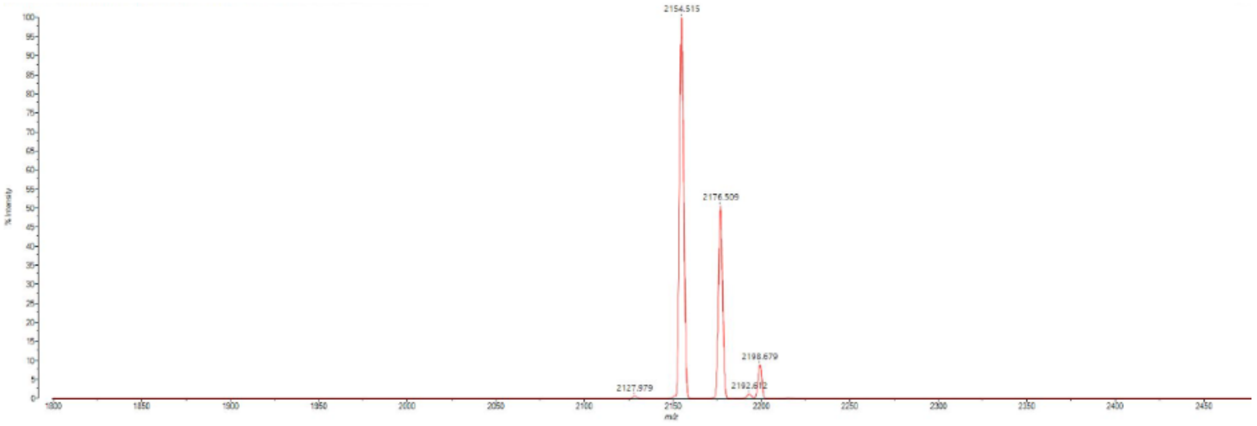

2154.515
2176.509
2198.679
2127.979

#### PenArg

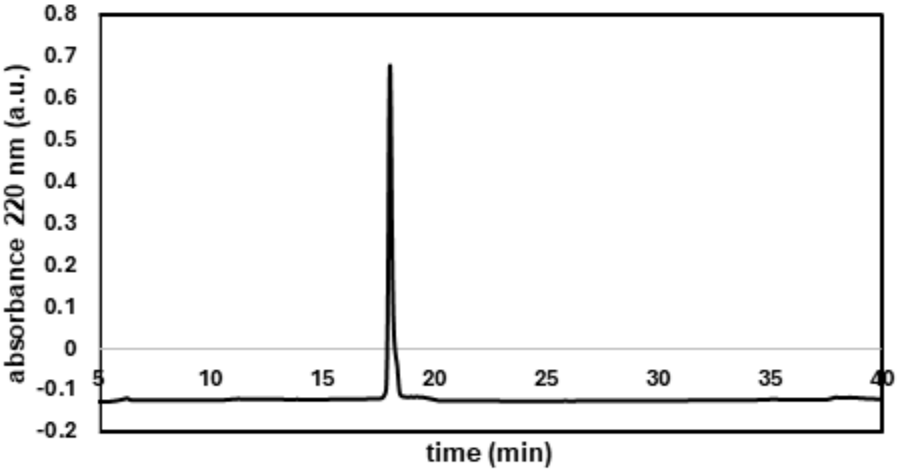
absorbance 220 nm (a.u.)
time (min)
0.8
0.7
0.6
0.5
0.4
0.3
0.2
0.1
0
-0.1
-0.2
5
10
15
20
25
30
35
40

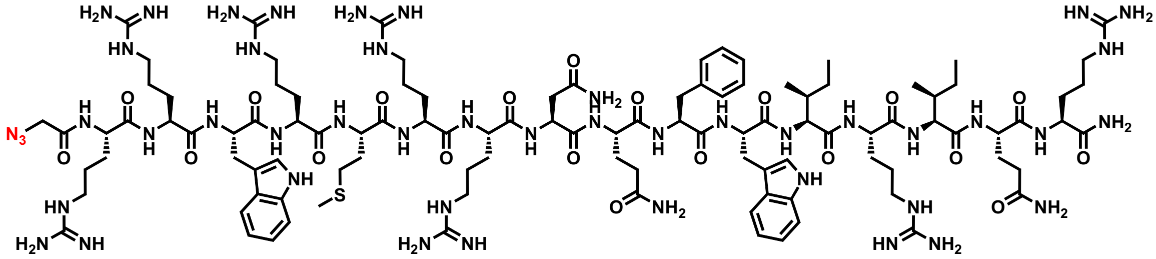

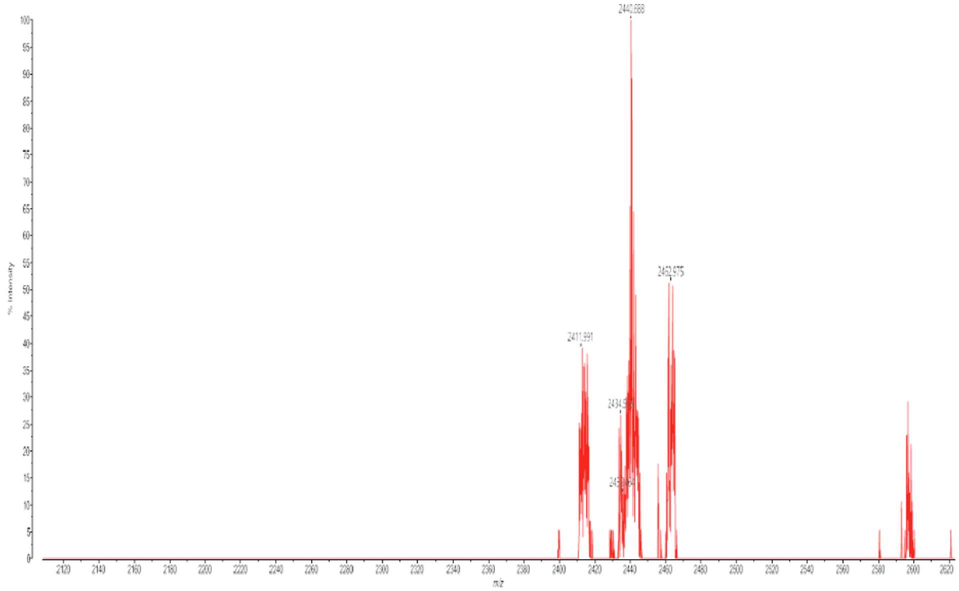

**Penetratin**

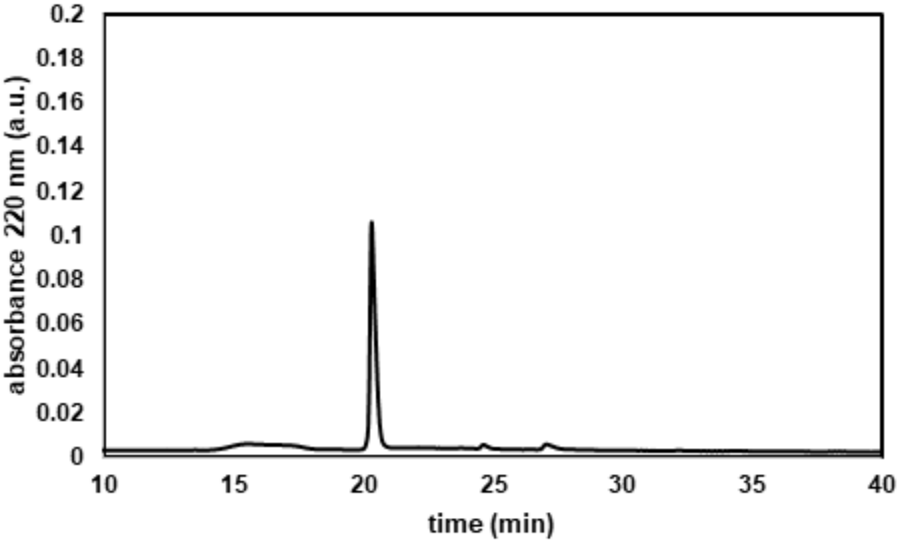

0.2
0.18
0.16
0.14
0.12
0.1
0.08
0.06
0.04
0.02
0
absorbance 220 nm (a.u.)
10
15
20
25
30
35
40
time (min)


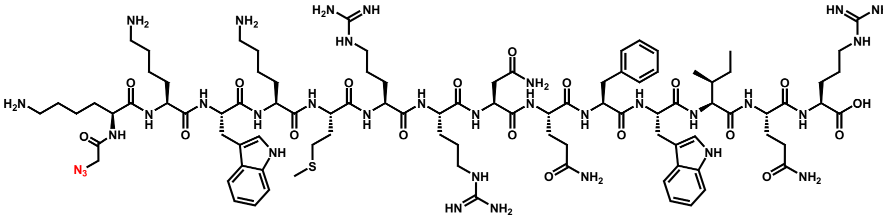

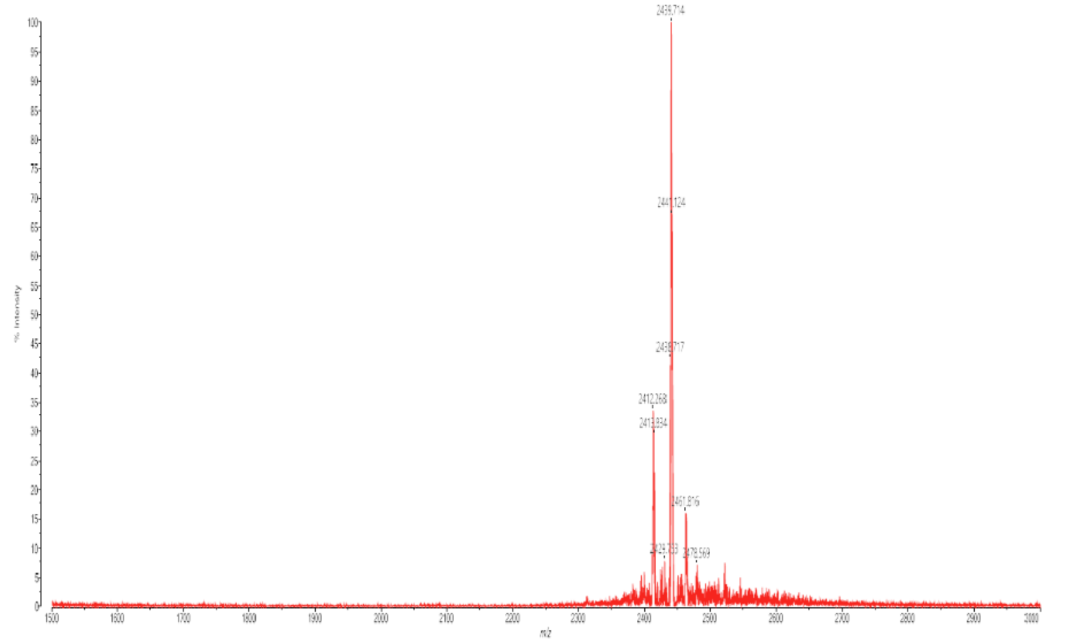

**Buforin II (5-21)**

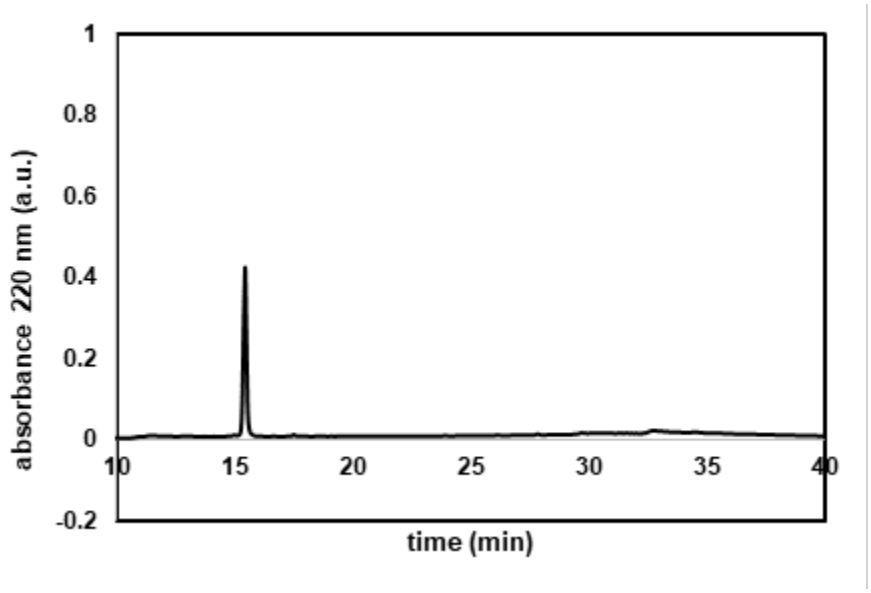


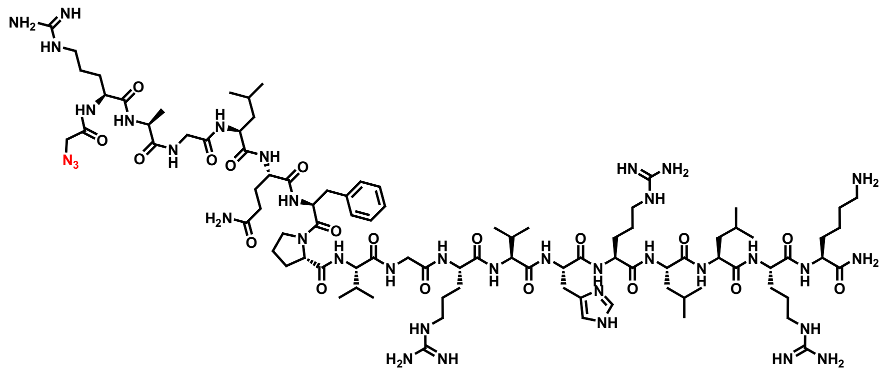

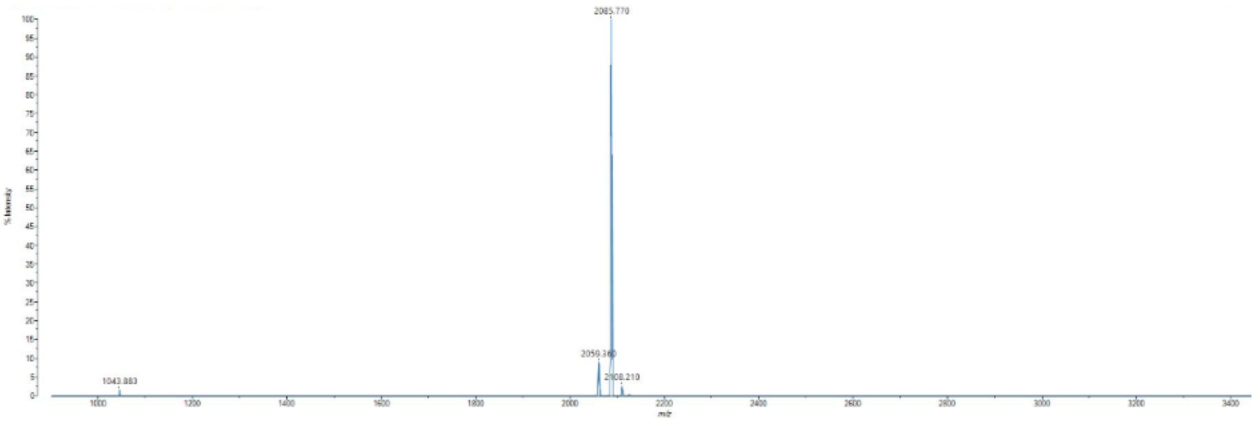

#### pAnTp (44-58)

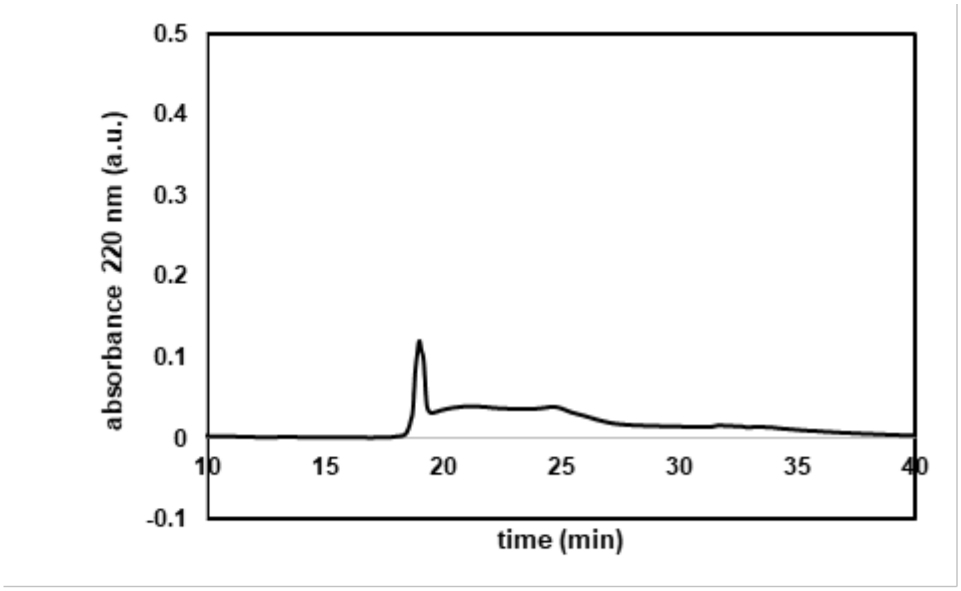

absorbance 220 nm (a.u.)
0.5
0.4
0.3
0.2
0.1
0
-0.1
10
15
20
25
30
35
40
time (min)


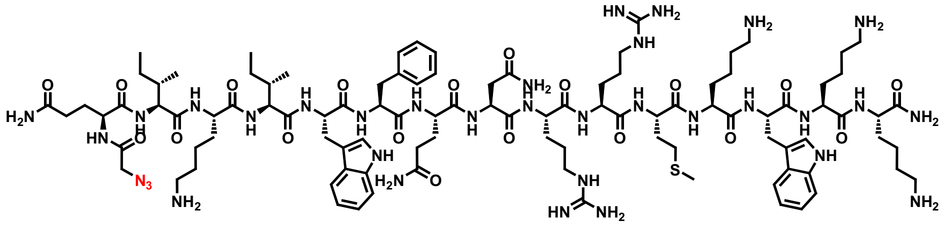

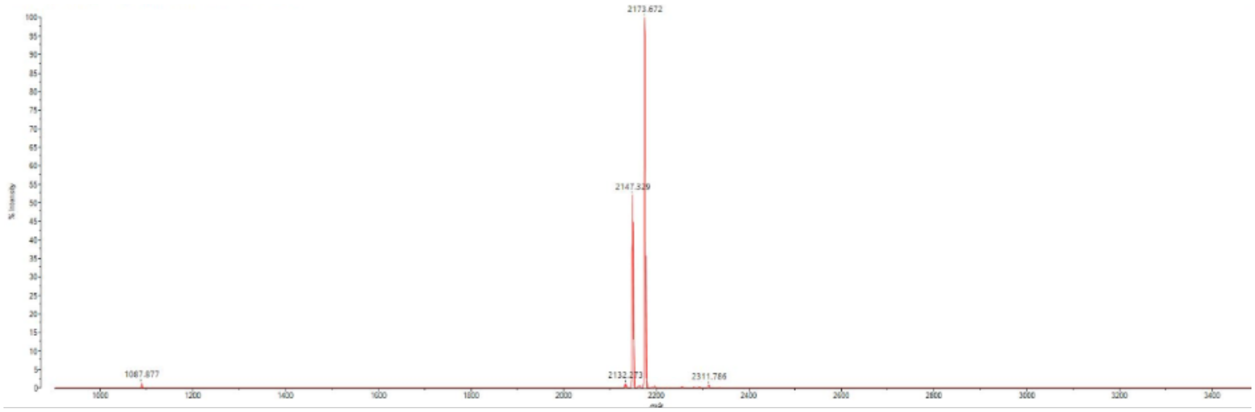

2173.672
2147.329
1087.877
2132.273
2311.786

**Cyt C 77-101**

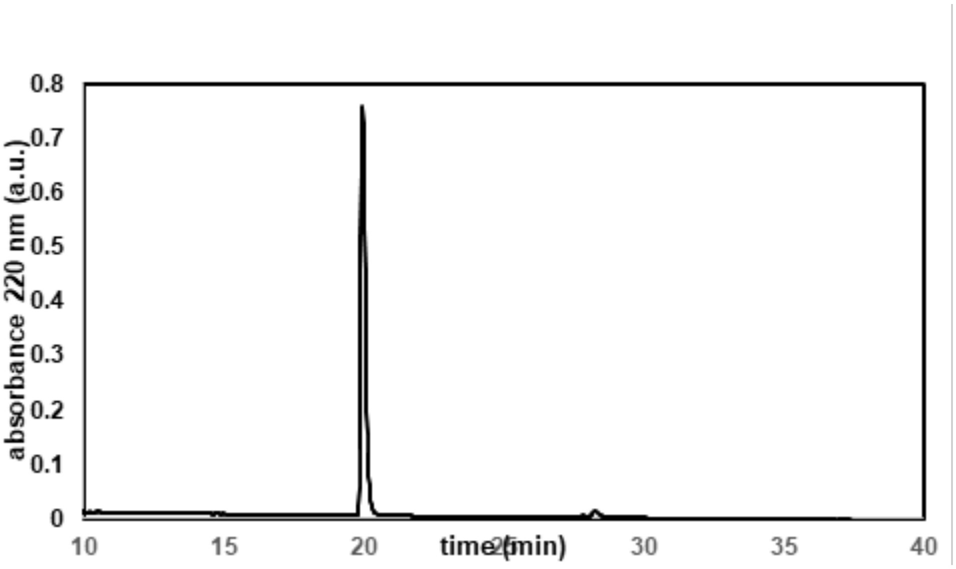
absorbance 220 nm (a.u.)
0.8
0.7
0.6
0.5
0.4
0.3
0.2
0.1
0
10
15
20
25
30
35
40
time (min)

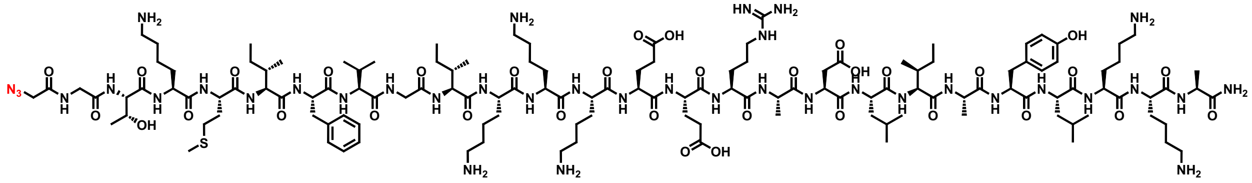
N3

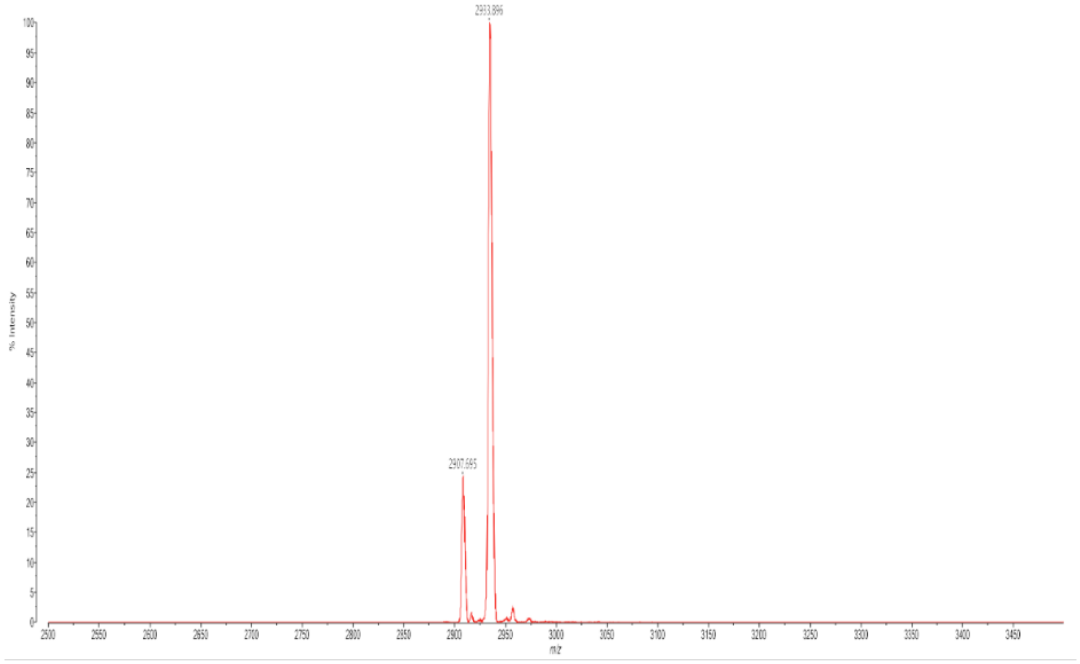
2933.896
2907.695
% Intensity
m/z

**M511**

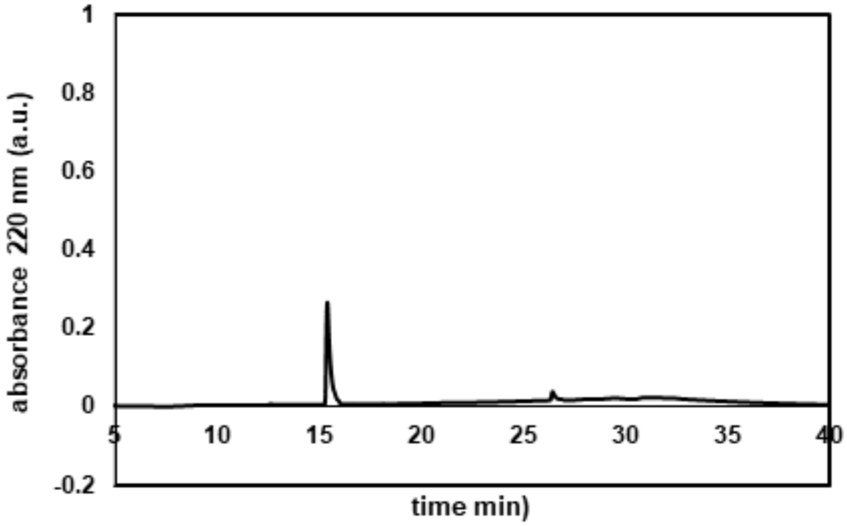

1
0.8
0.6
0.4
0.2
0
-0.2
5
10
15
20
25
30
35
40
absorbance 220 nm (a.u.)
time min)


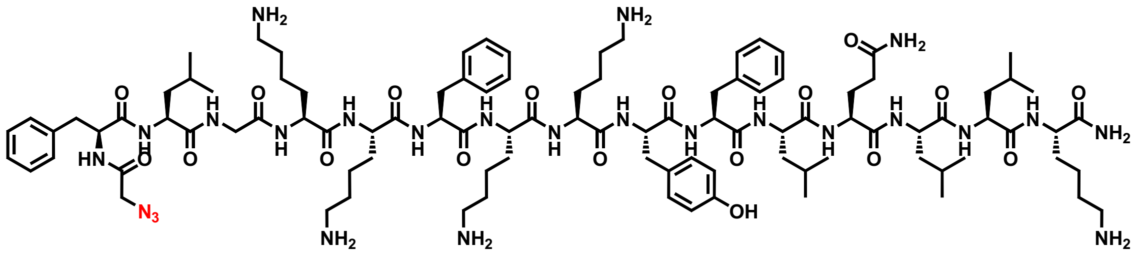

NH2
NH2
O
NH2
HN
N3
OH
NH2
NH2
NH2


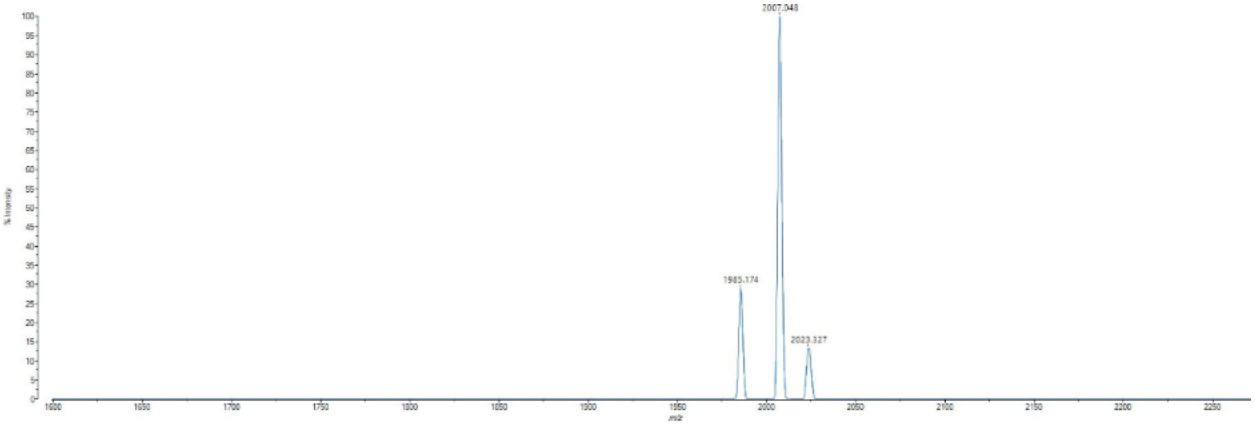

2007.048
1985.174
2023.327
2000
2050

**M591**

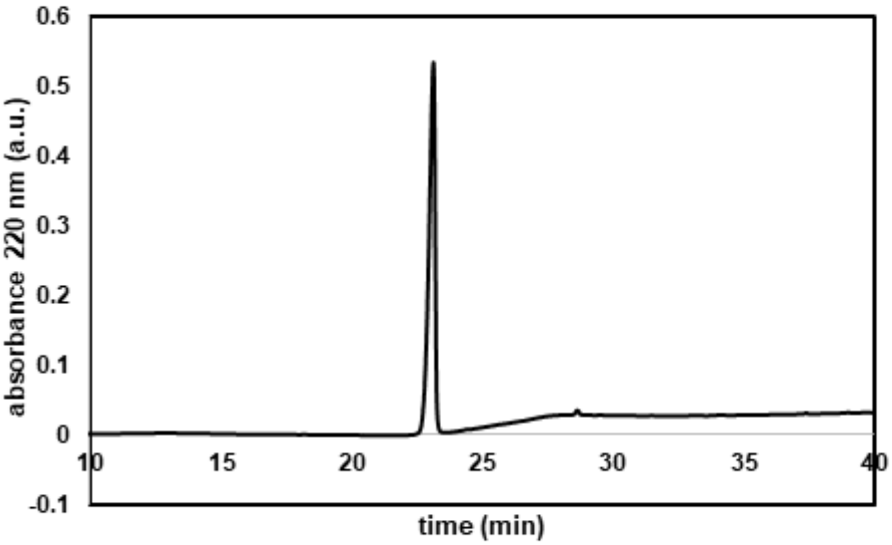

absorbance 220 nm (a.u.)
time (min)
0.6
0.5
0.4
0.3
0.2
0.1
0
-0.1
10
15
20
25
30
35
40


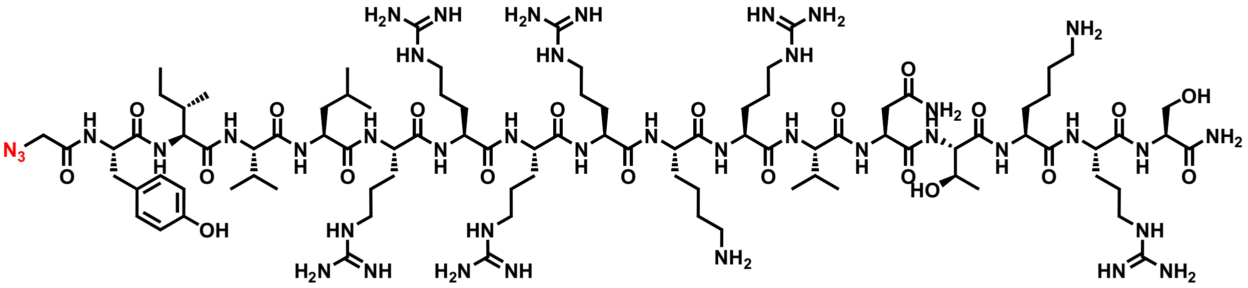

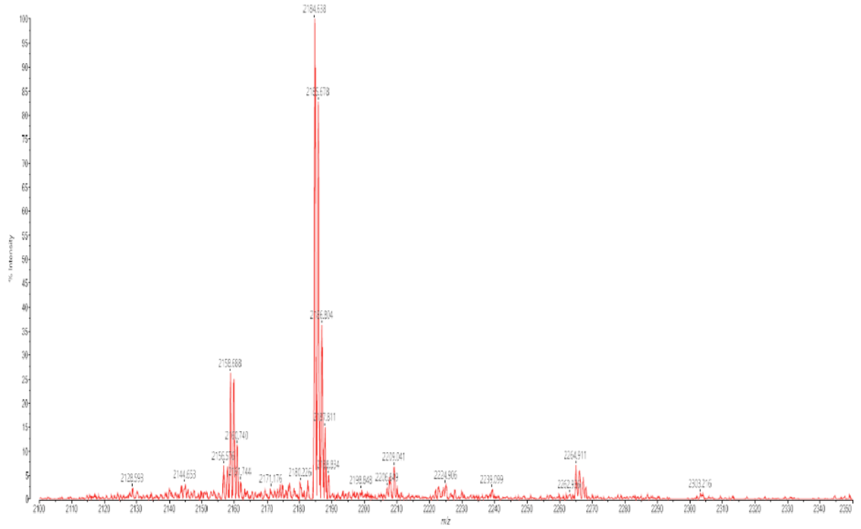

## E162

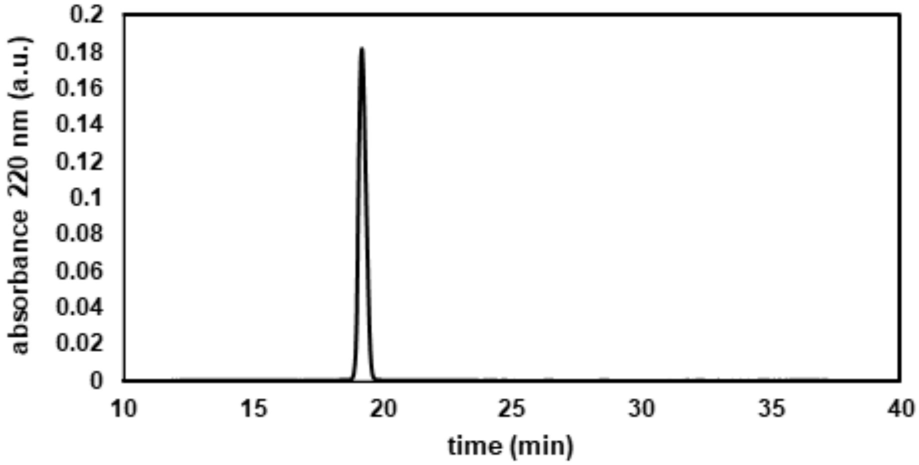
absorbance 220 nm (a.u.)
0.2
0.18
0.16
0.14
0.12
0.1
0.08
0.06
0.04
0.02
0
10
15
20
25
30
35
40
time (min)

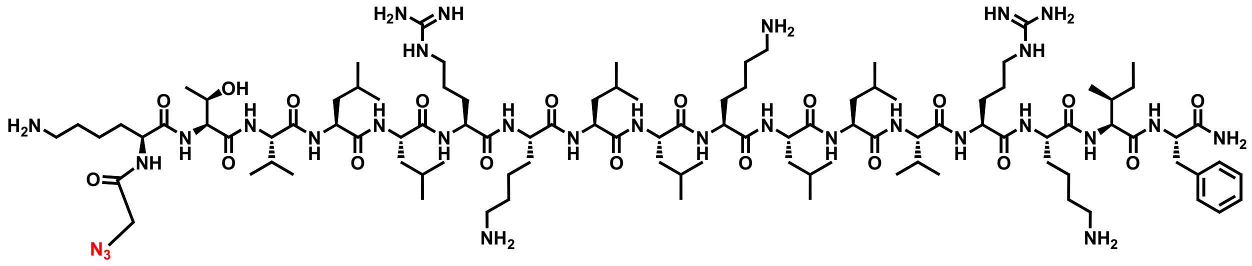

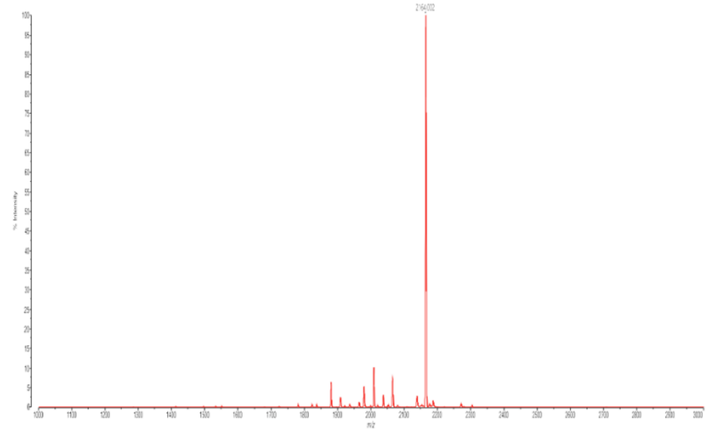

**E165**

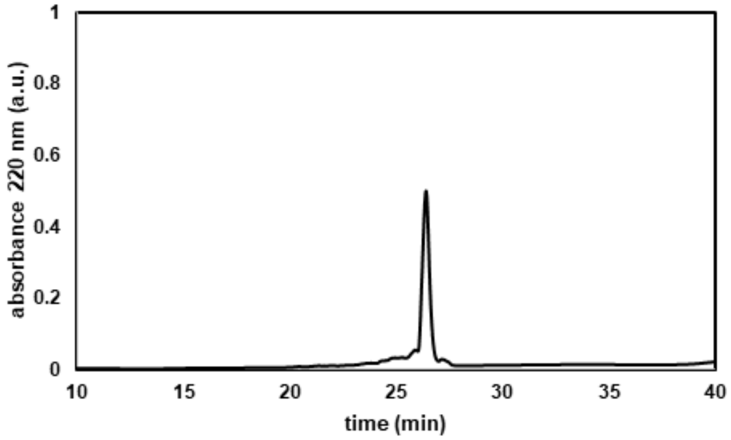


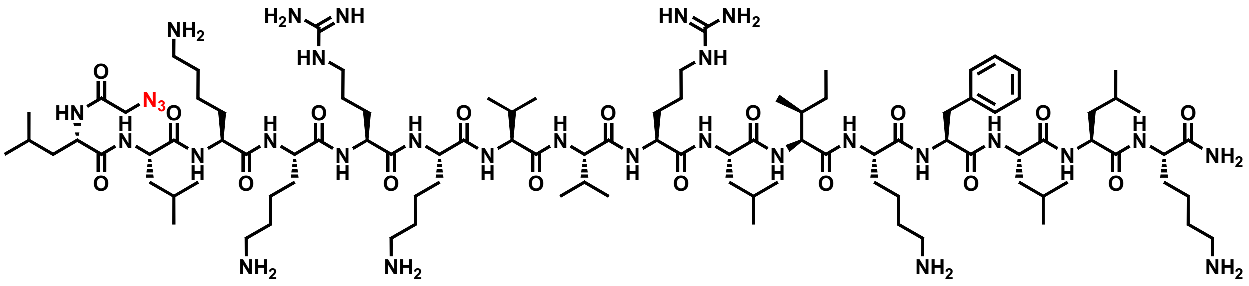

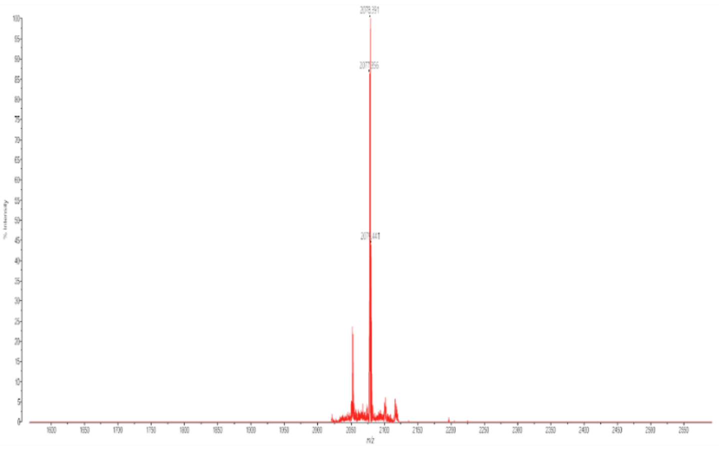

**M867**

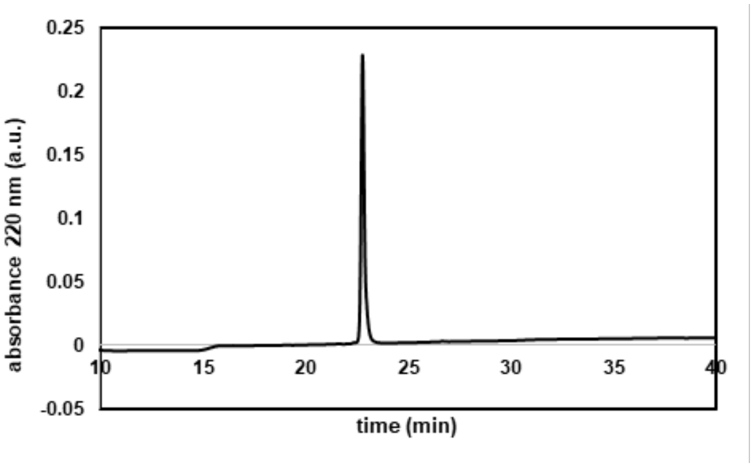
absorbance 220 nm (a.u.)
time (min)
0.25
0.2
0.15
0.1
0.05
0
-0.05
10
15
20
25
30
35
40

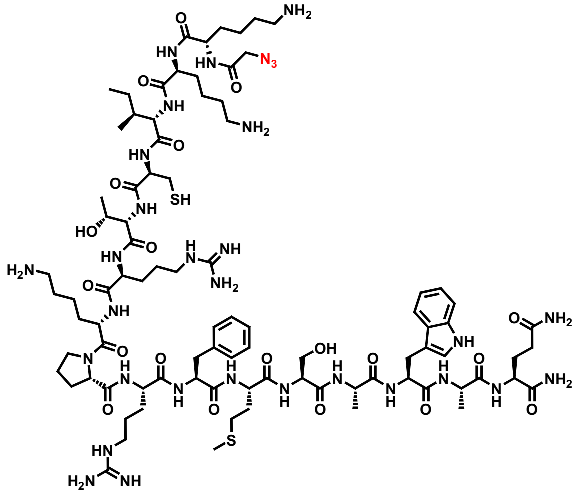

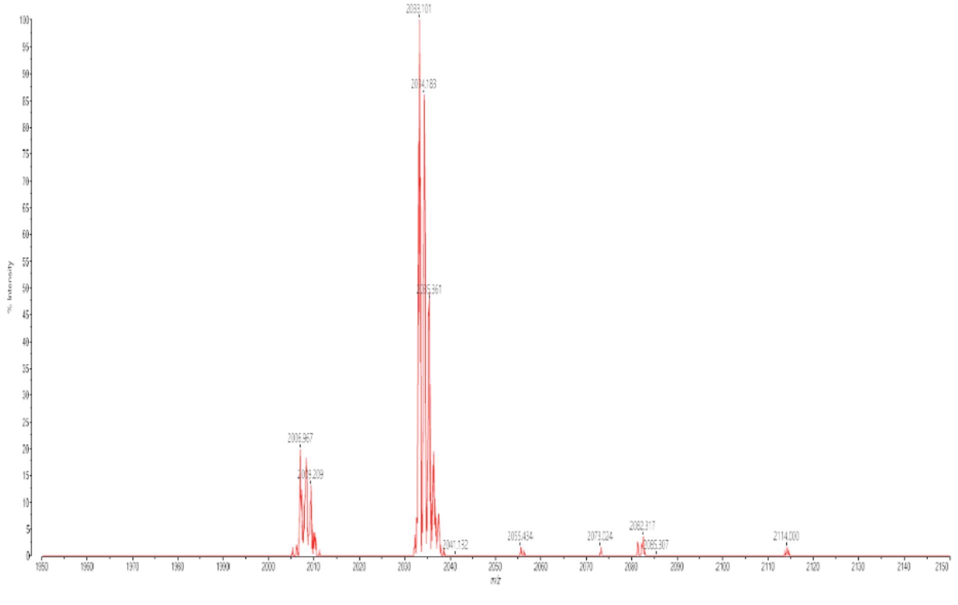

#### Crot (27-39)

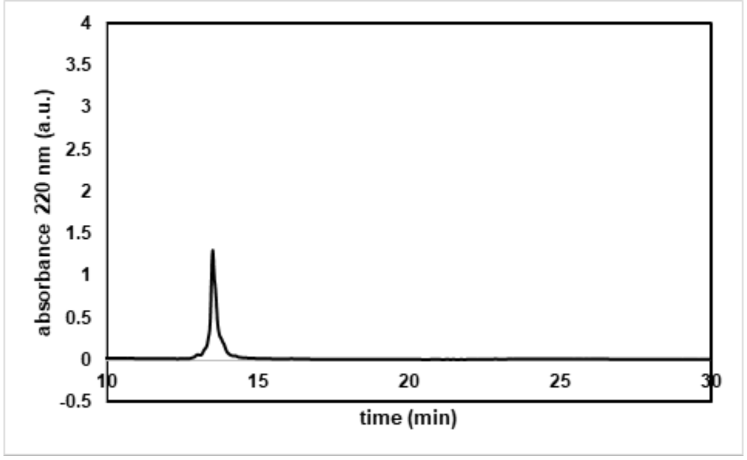
absorbance 220 nm (a.u.)
4
3.5
3
2.5
2
1.5
1
0.5
0
-0.5
10
15
20
25
30
time (min)

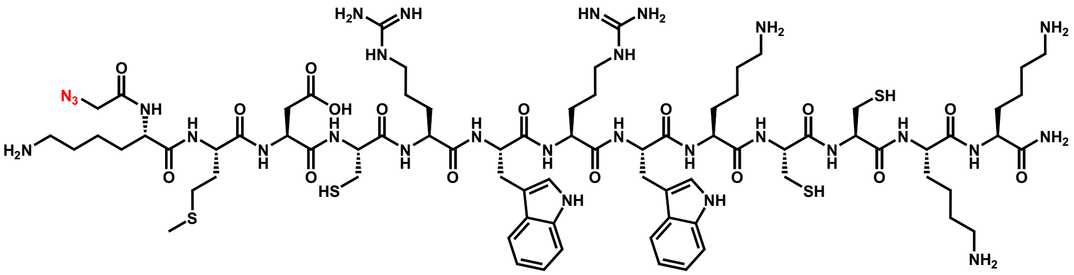

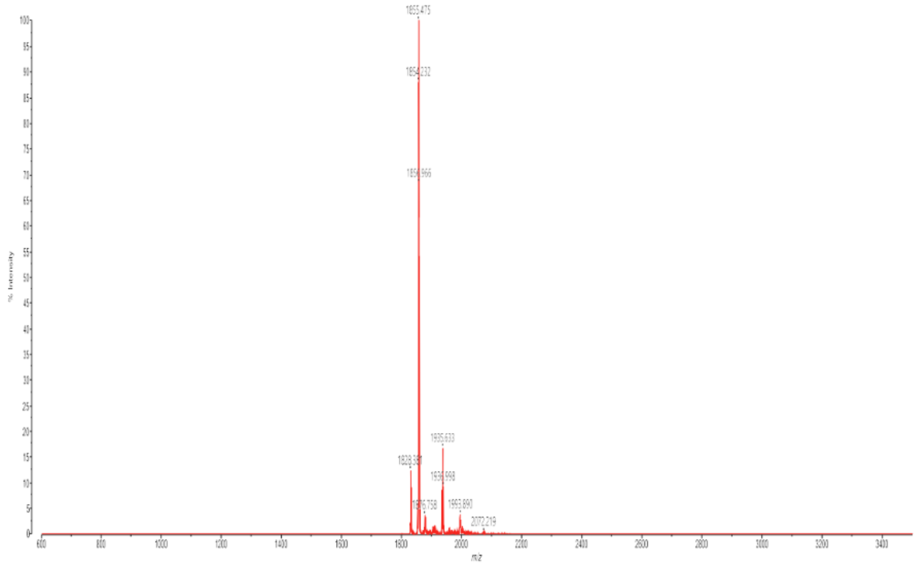
1855.475

#### Crot (27-39) derivative (2)

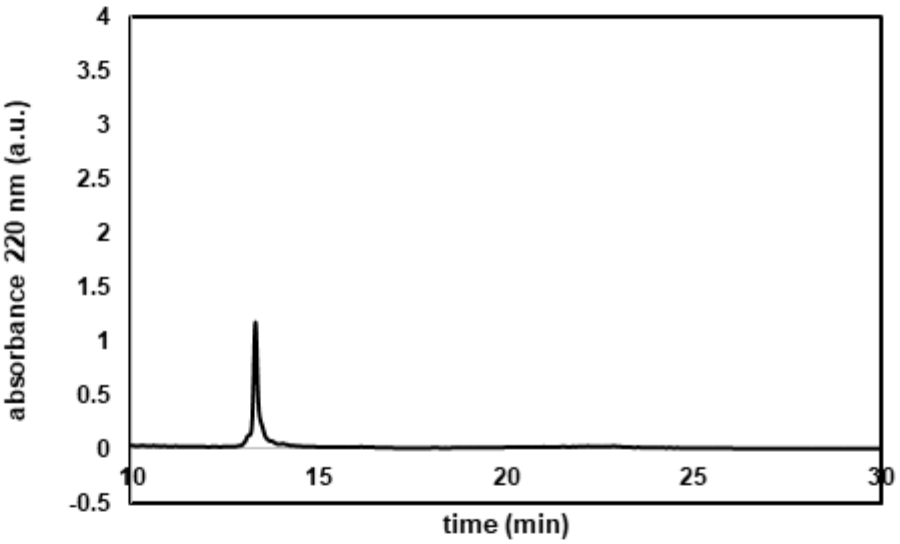


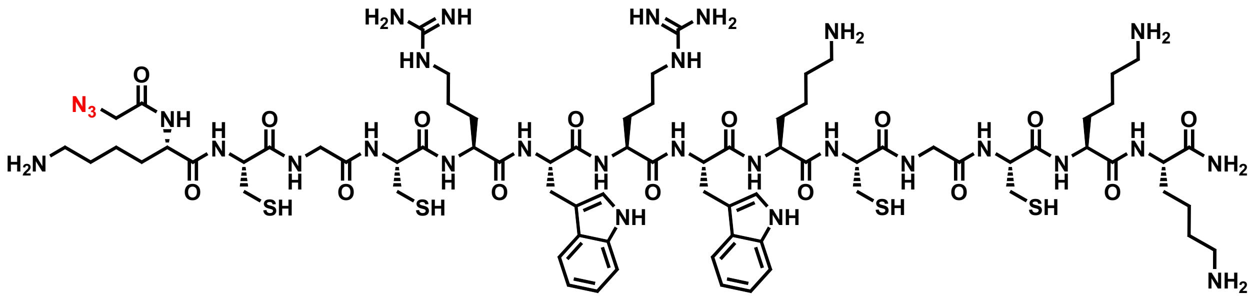


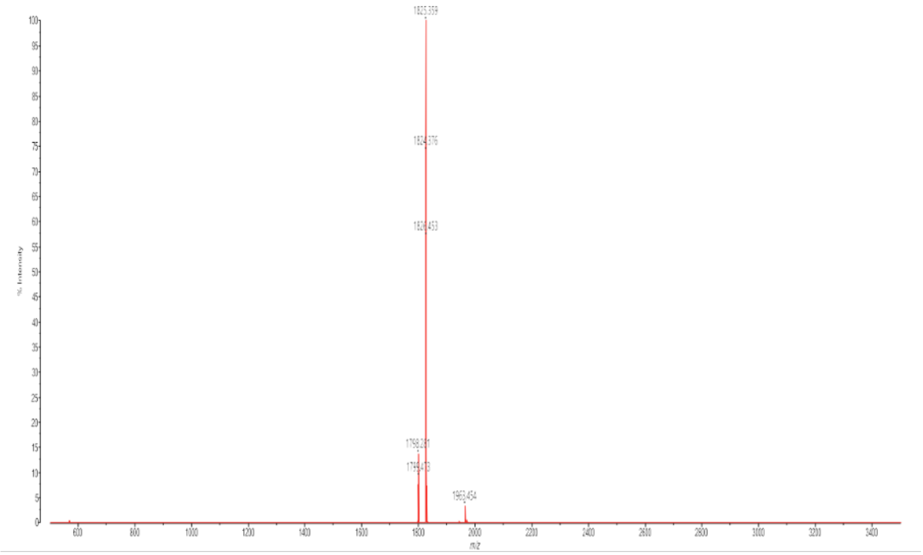

**CyLoP-1**

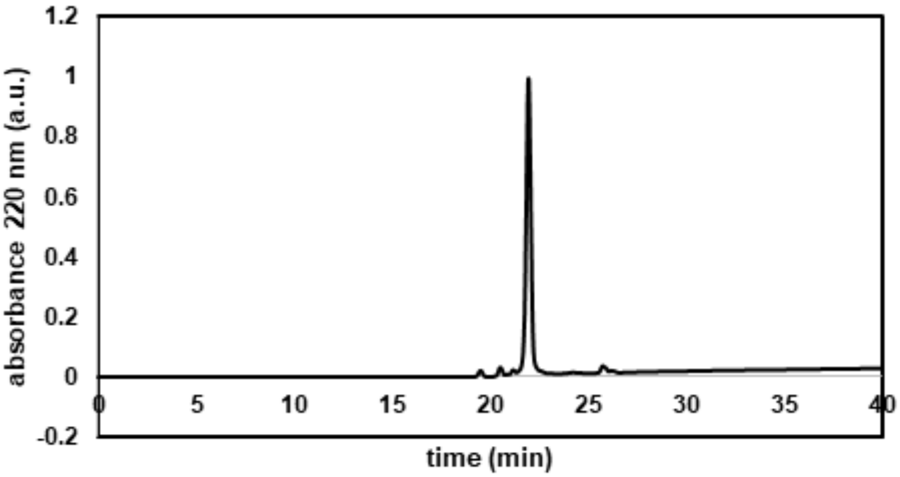
1.2
1
0.8
0.6
0.4
0.2
0
-0.2
absorbance 220 nm (a.u.)
0
5
10
15
20
25
30
35
40
time (min)

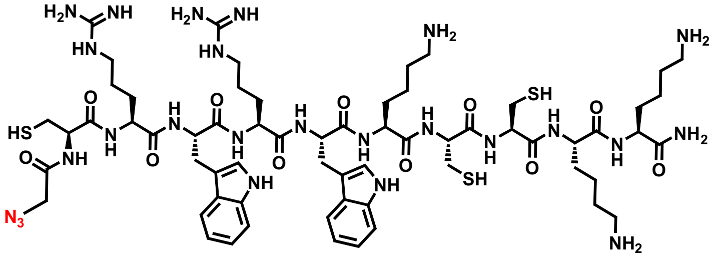

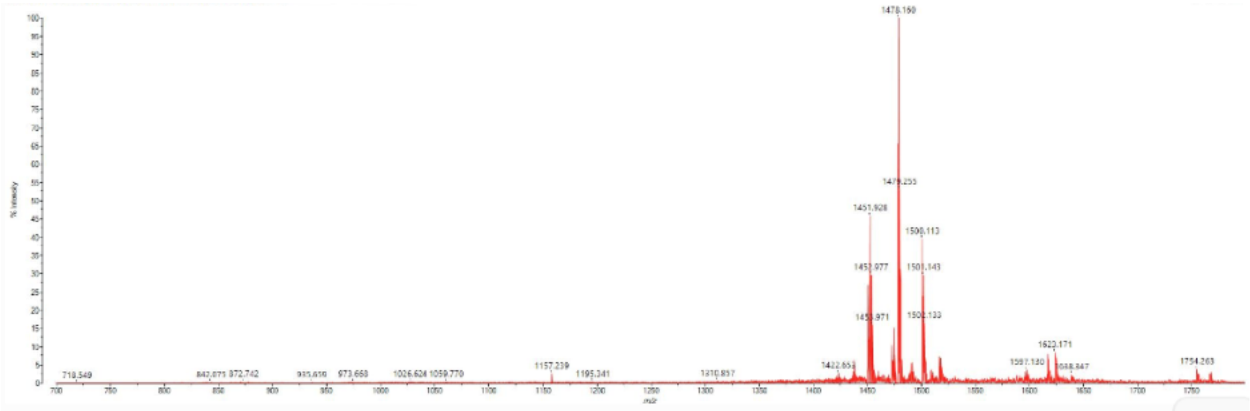

**HPV33L2-DD447**

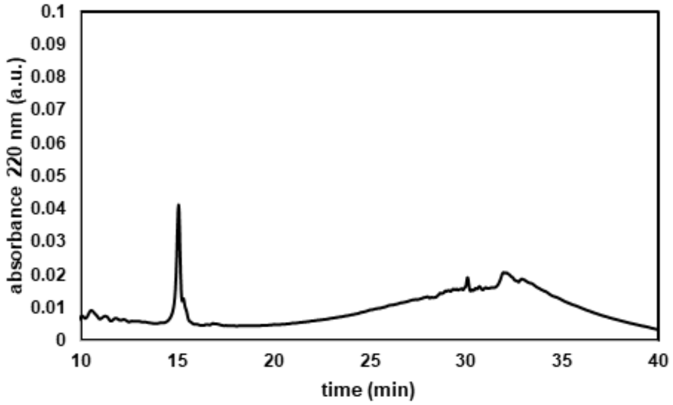


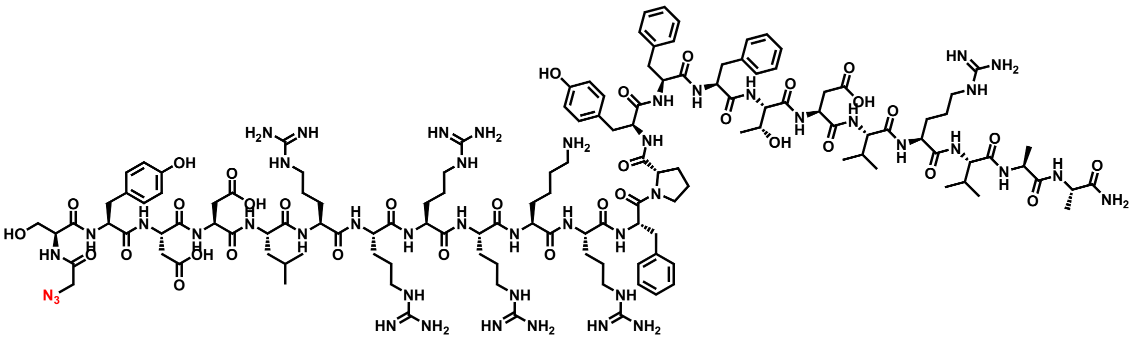

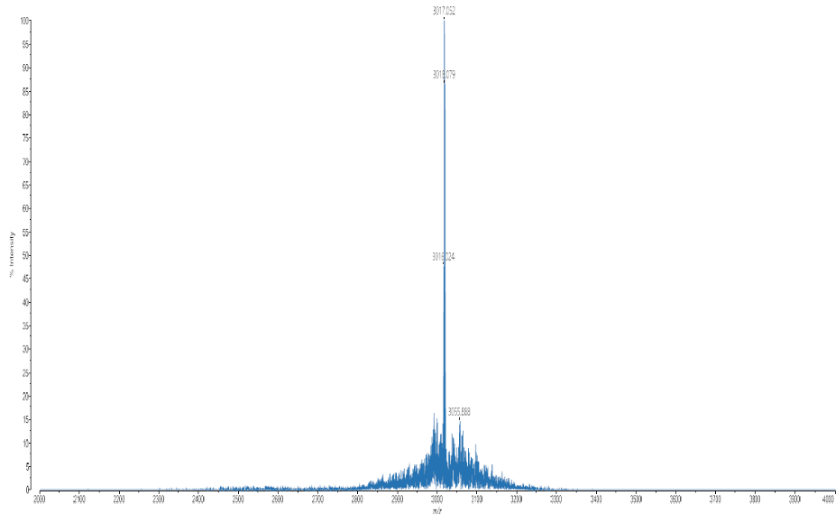

#### HPV33L2- 445/467

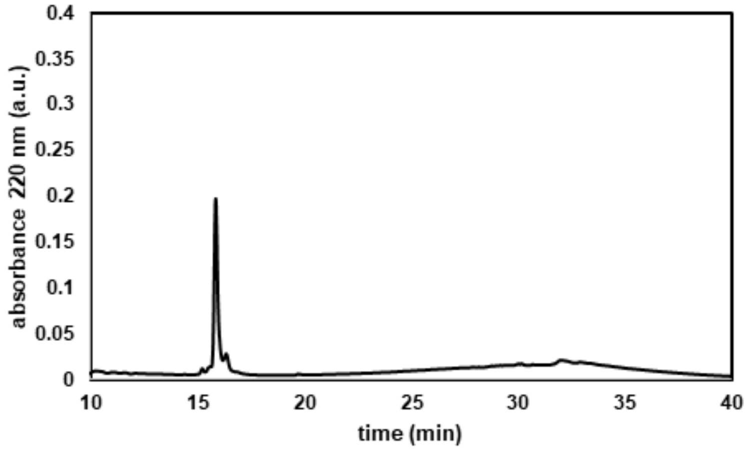


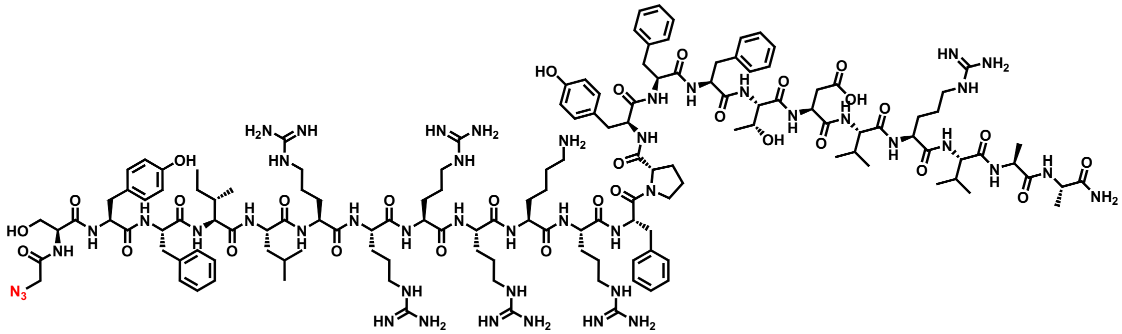

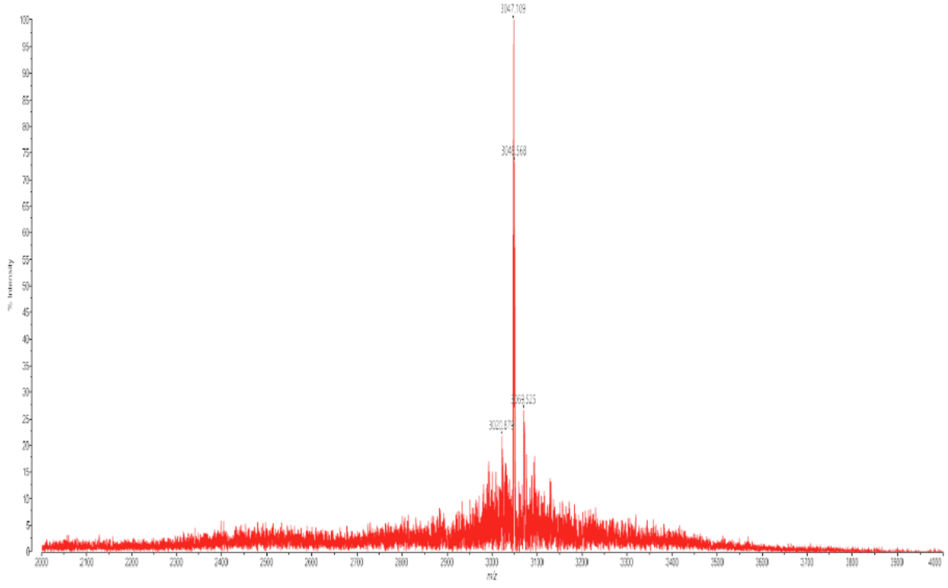

**KLA10**

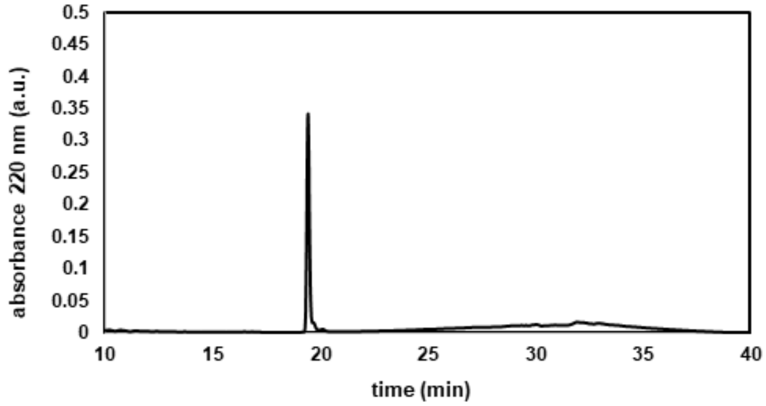

0.5
0.45
0.4
0.35
0.3
0.25
0.2
0.15
0.1
0.05
0
absorbance 220 nm (a.u.)
10
15
20
25
30
35
40
time (min)


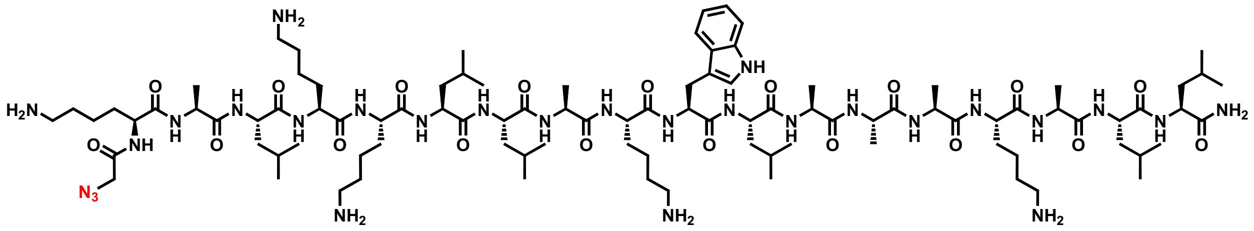

$H_2N$
$N_3$
$NH_2$
NH
$NH_2$


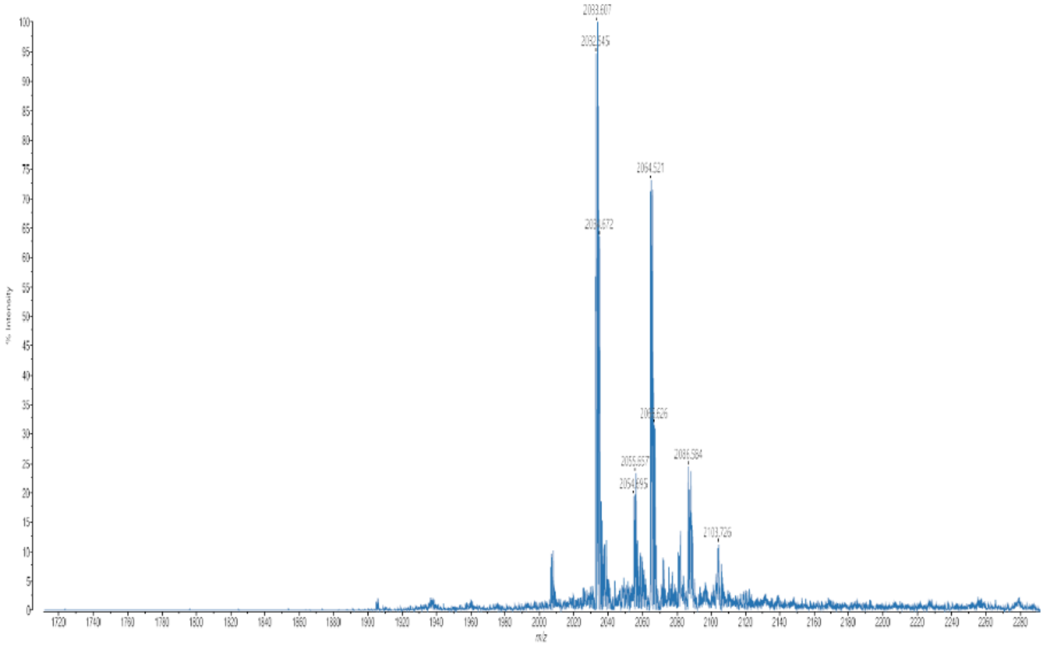

2033.607
2032.545
2064.521
2055.657
2054.895
2086.584
2103.726
% Intensity
m/z

**No. 14-12**

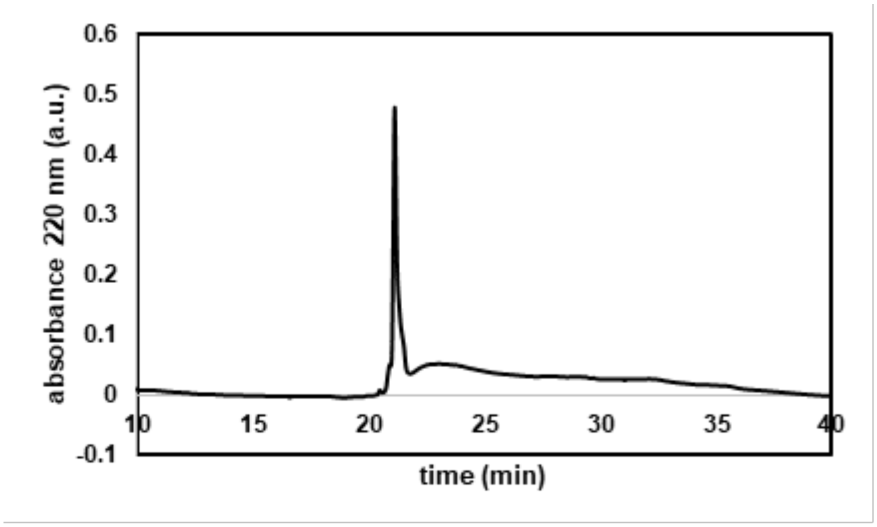
absorbance 220 nm (a.u.)
time (min)
0.6
0.5
0.4
0.3
0.2
0.1
0
-0.1
10
15
20
25
30
35
40

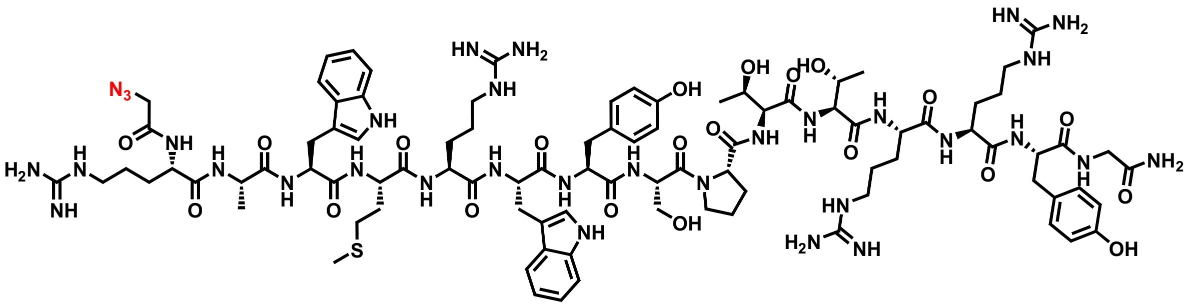

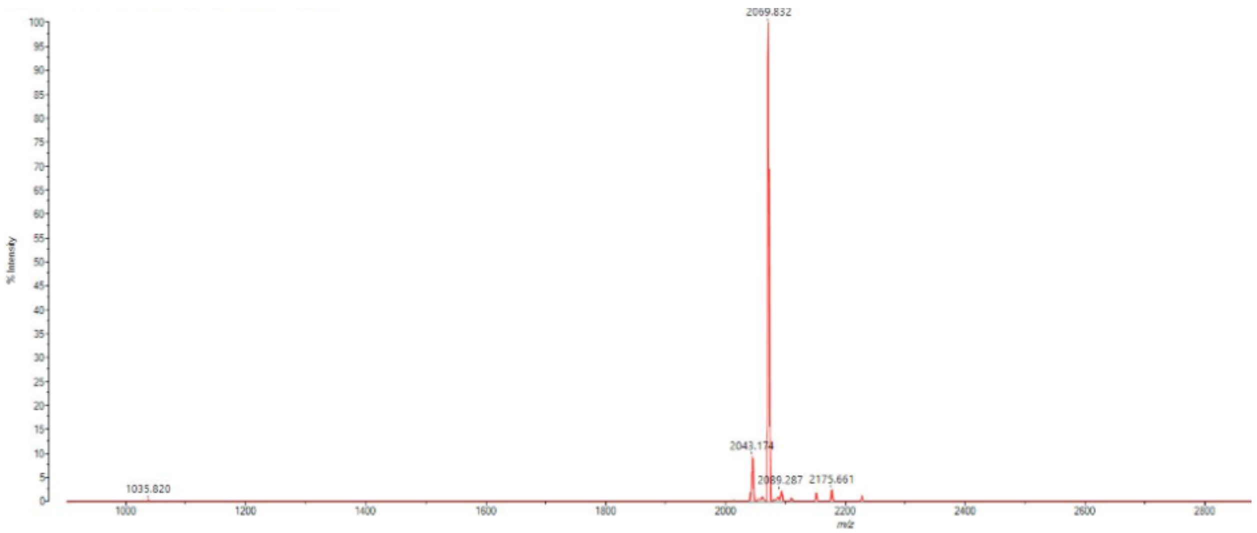
2069.832
2043.174
2089.287
2175.661
1035.820
% Intensity
m/z

**dhvar2**

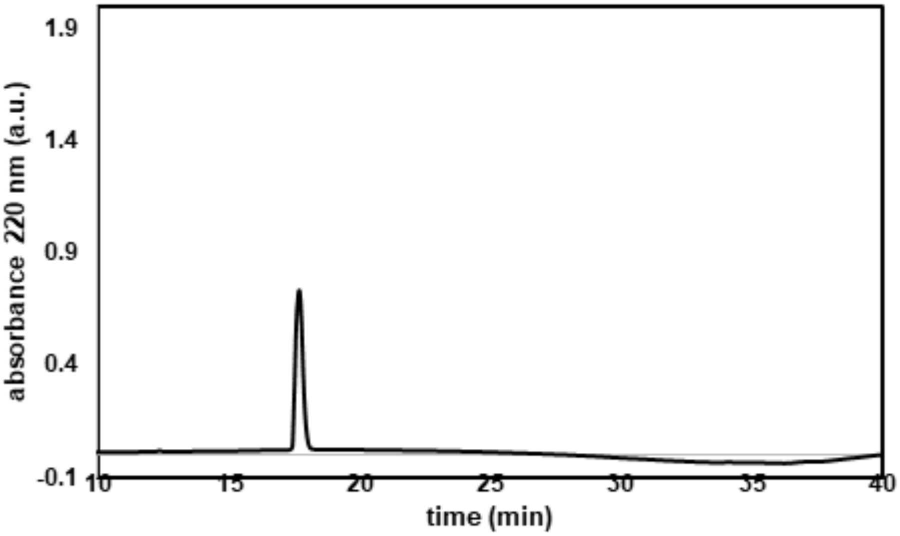

absorbance 220 nm (a.u.)
1.9
1.4
0.9
0.4
-0.1
10
15
20
25
30
35
40
time (min)


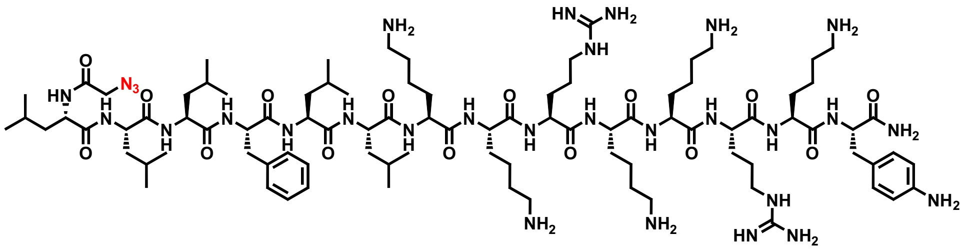

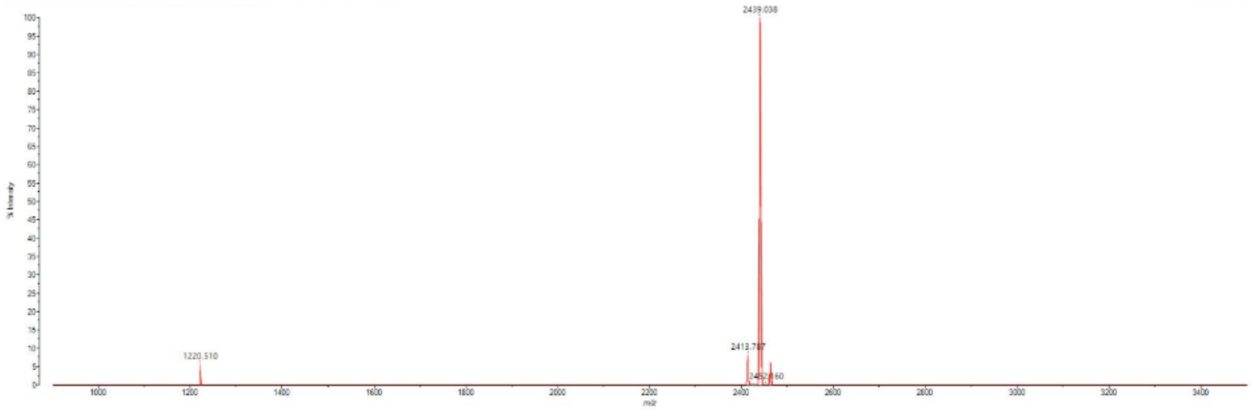

2439.038
1220.510

**ppTG1**

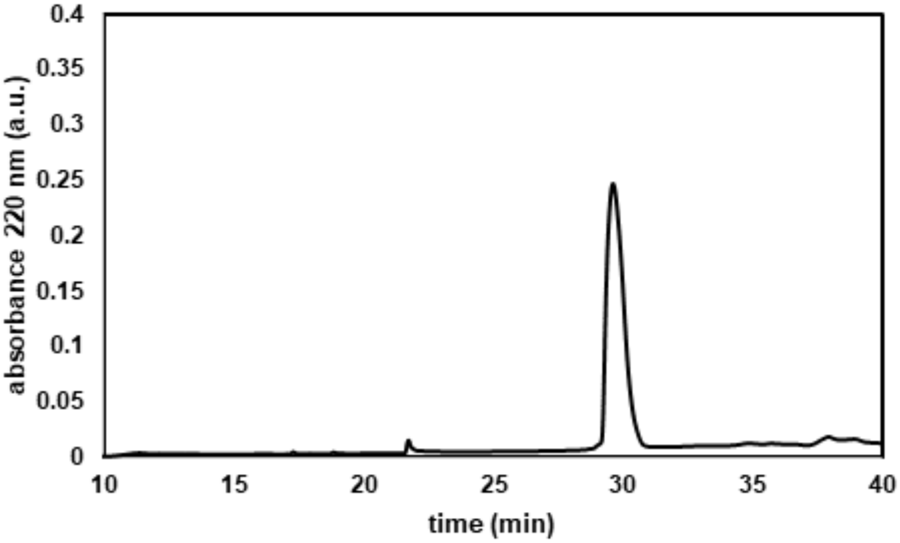

absorbance 220 nm (a.u.)
time (min)
0.4
0.35
0.3
0.25
0.2
0.15
0.1
0.05
0
10
15
20
25
30
35
40


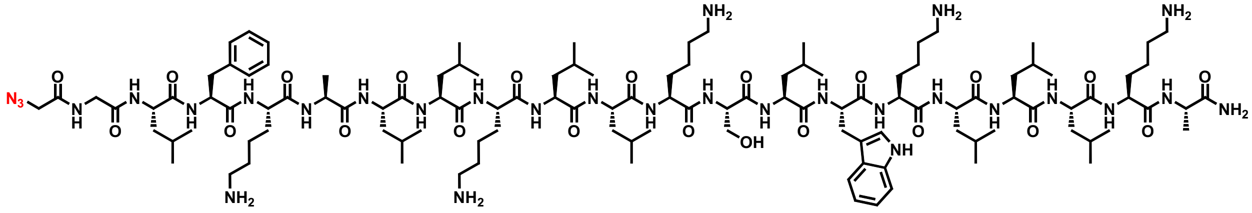

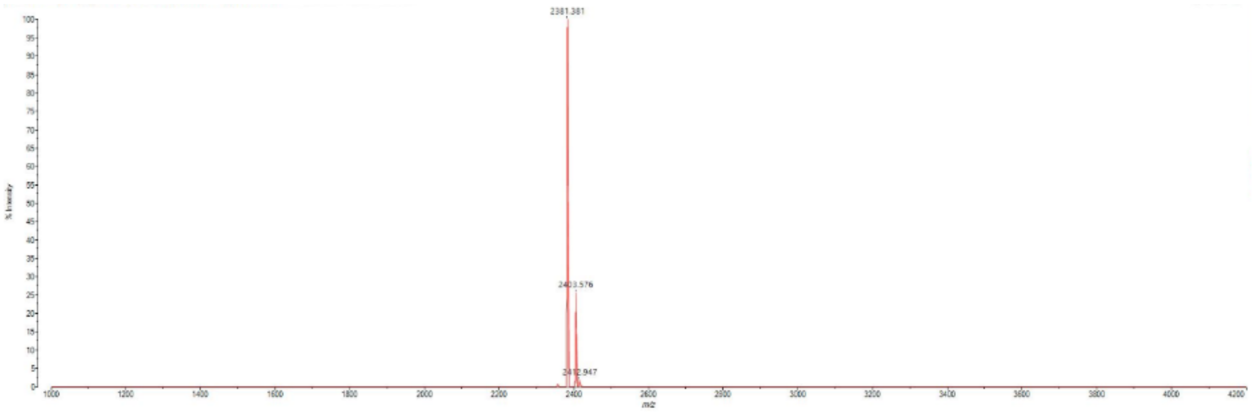

2381.381

**PolyP 3 (SAP)**

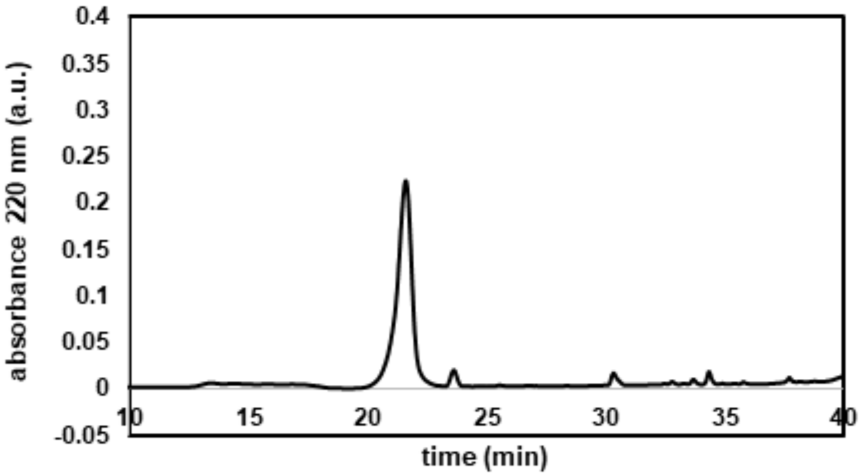

absorbance 220 nm (a.u.)
0.4
0.35
0.3
0.25
0.2
0.15
0.1
0.05
0
-0.05
10
15
20
25
30
35
40
time (min)


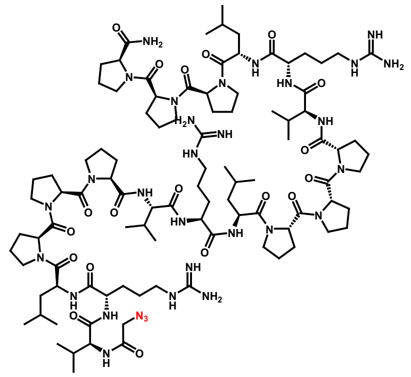

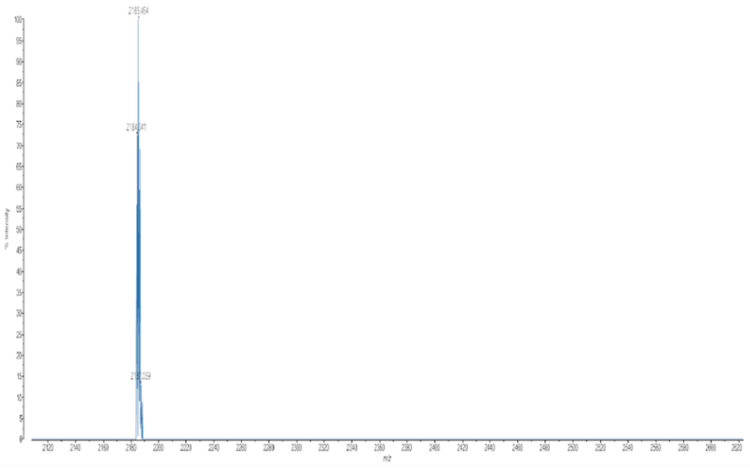

**LAH4**

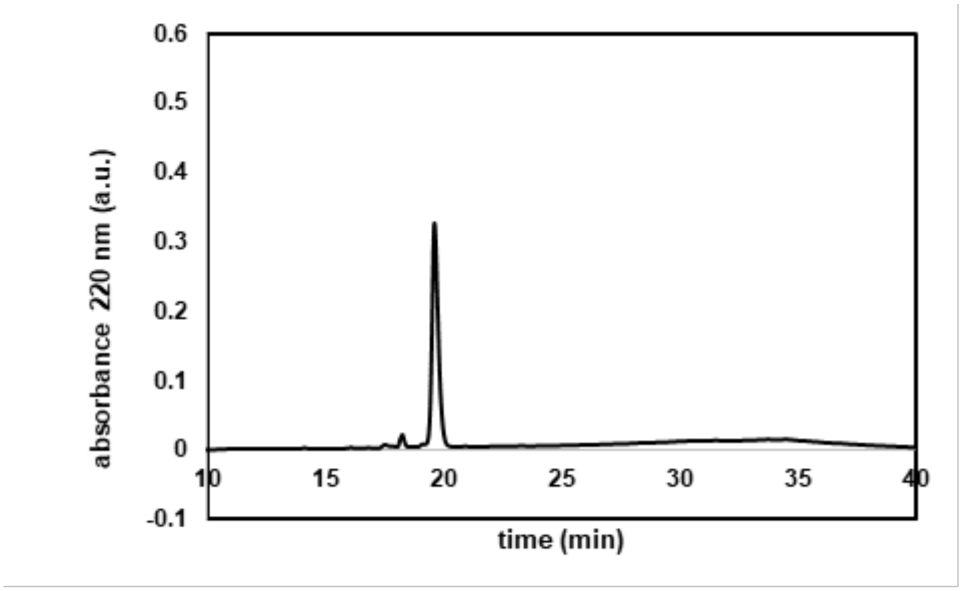

0.6
0.5
0.4
0.3
0.2
0.1
0
-0.1
absorbance 220 nm (a.u.)
10
15
20
25
30
35
40
time (min)


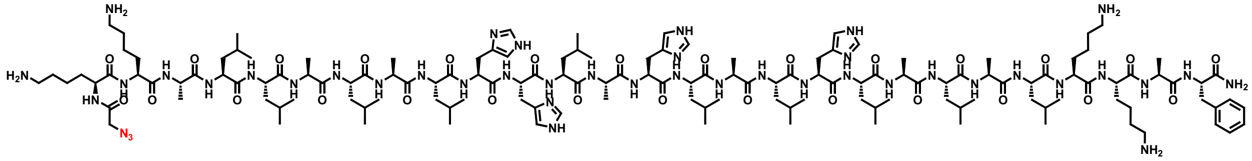

$H_2N$
$NH_2$
$N_3$
NH
$NH_2$


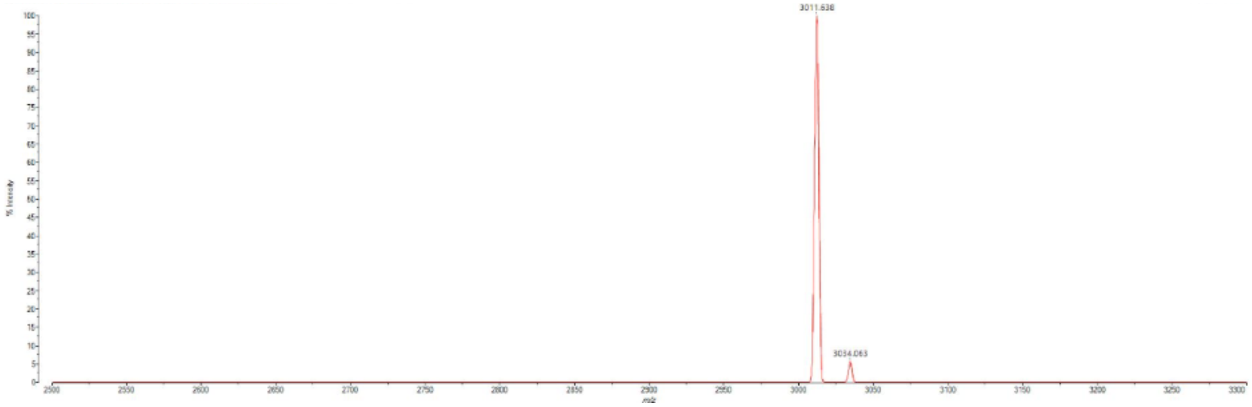

3011.638
3034.063

#### Transportan (TP)

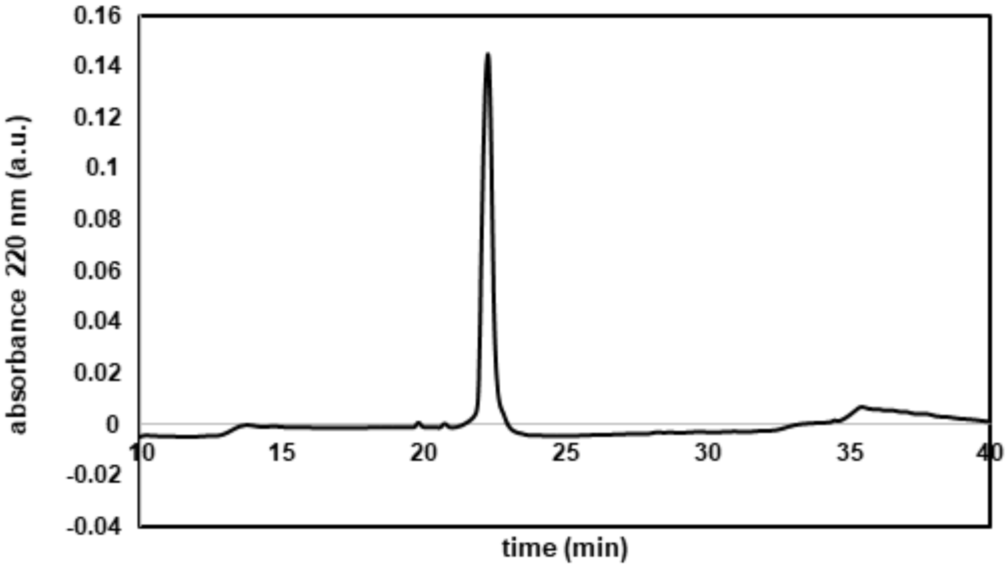


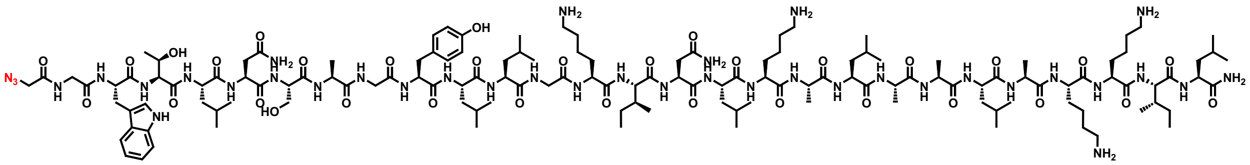

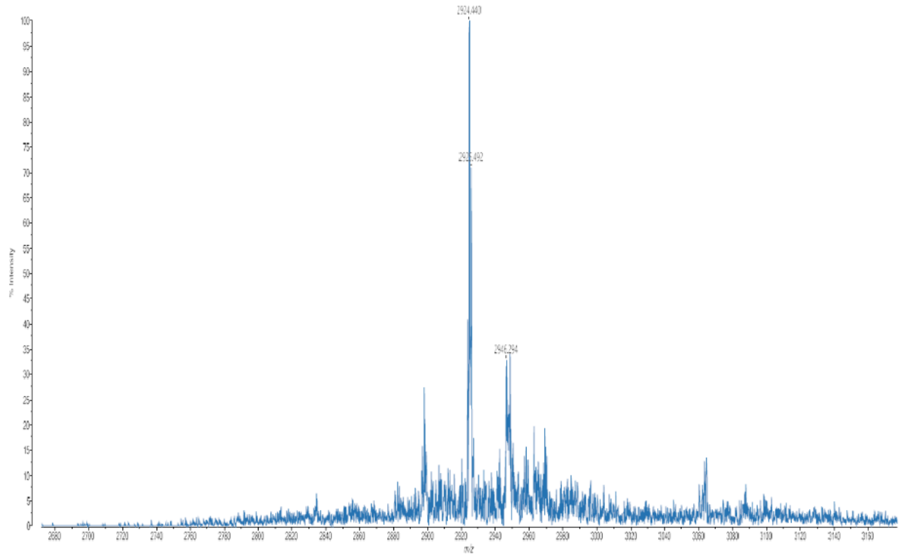

**InV3**

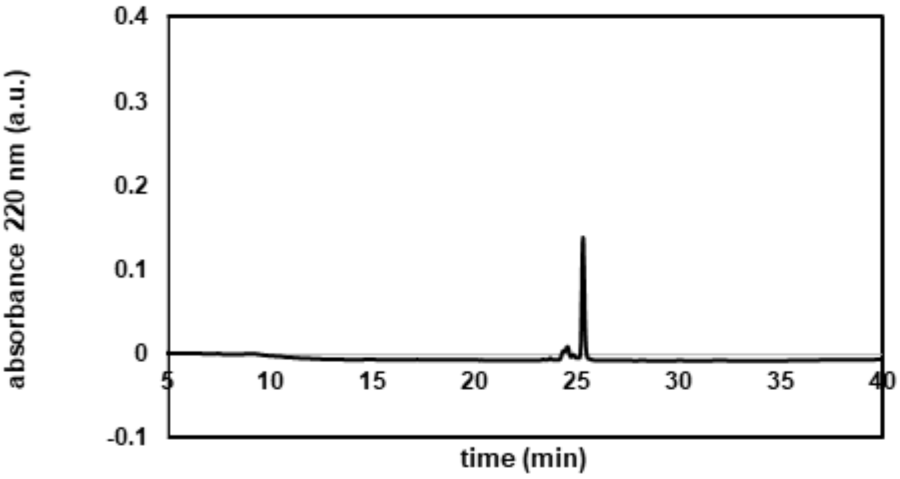

absorbance 220 nm (a.u.)
time (min)
0.4
0.3
0.2
0.1
0
-0.1
5
10
15
20
25
30
35
40


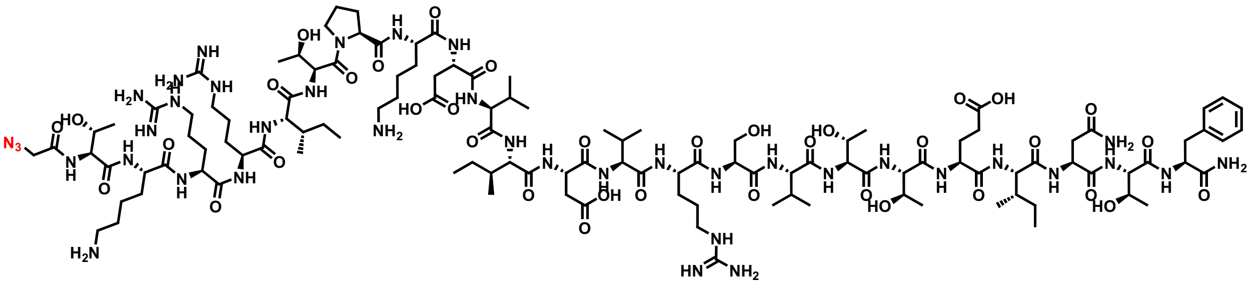

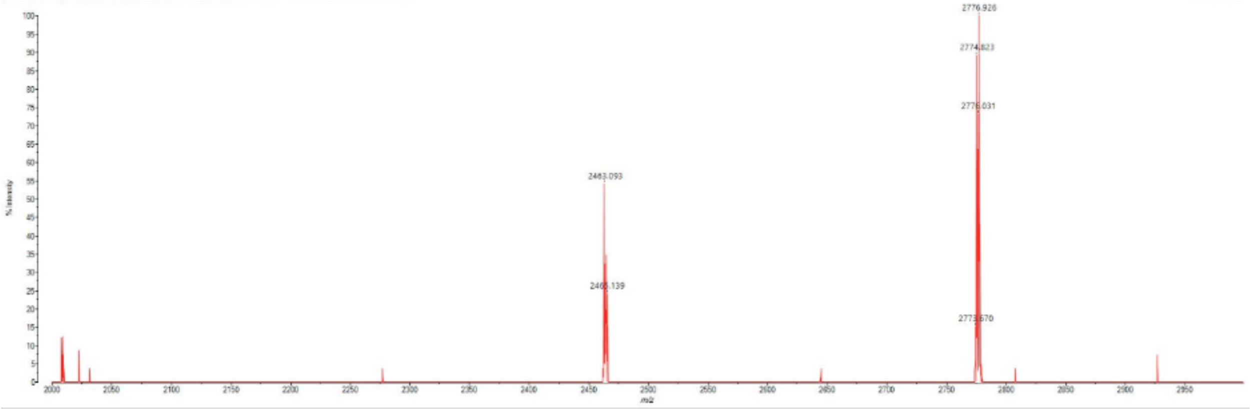

**MG2d**

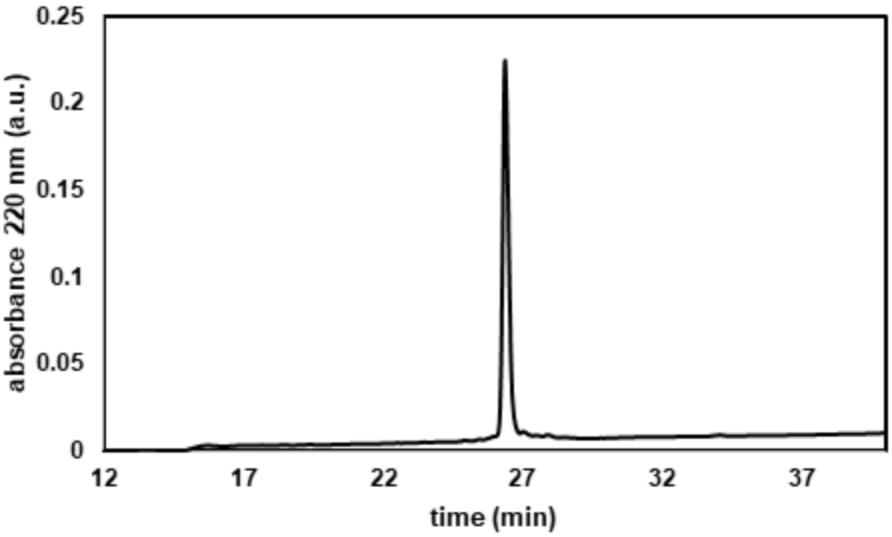


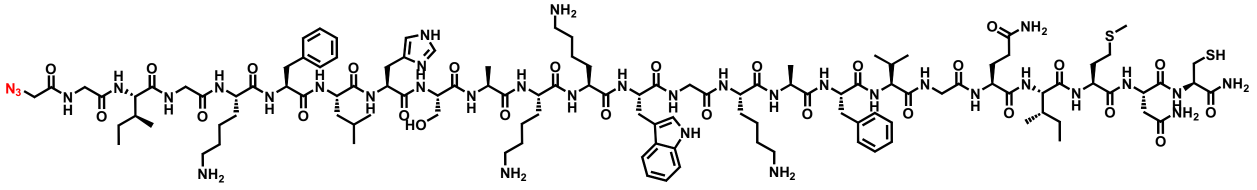

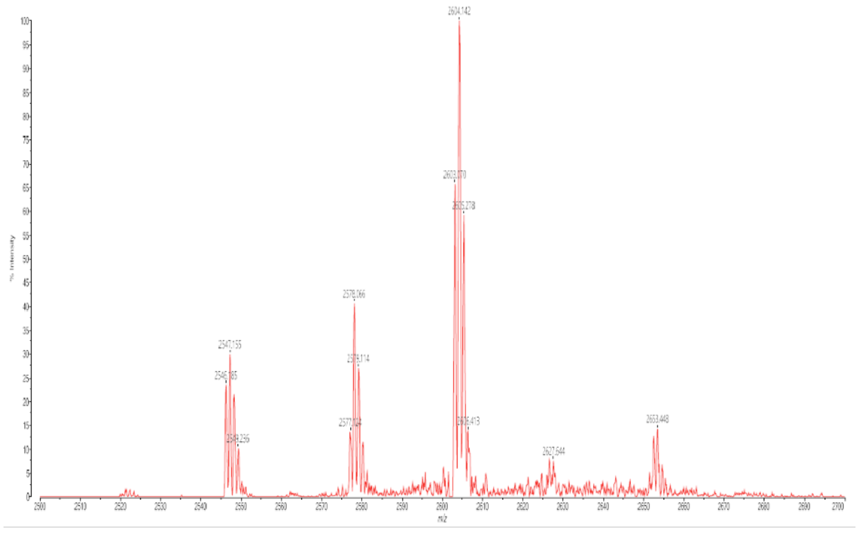

**InV3.5**

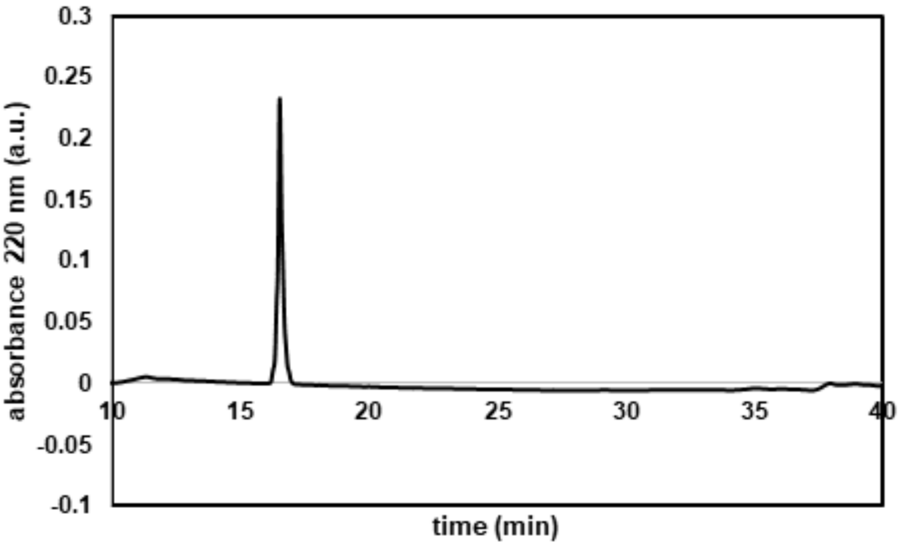

0.3
0.25
0.2
0.15
0.1
0.05
0
-0.05
-0.1
absorbance 220 nm (a.u.)
10
15
20
25
30
35
40
time (min)


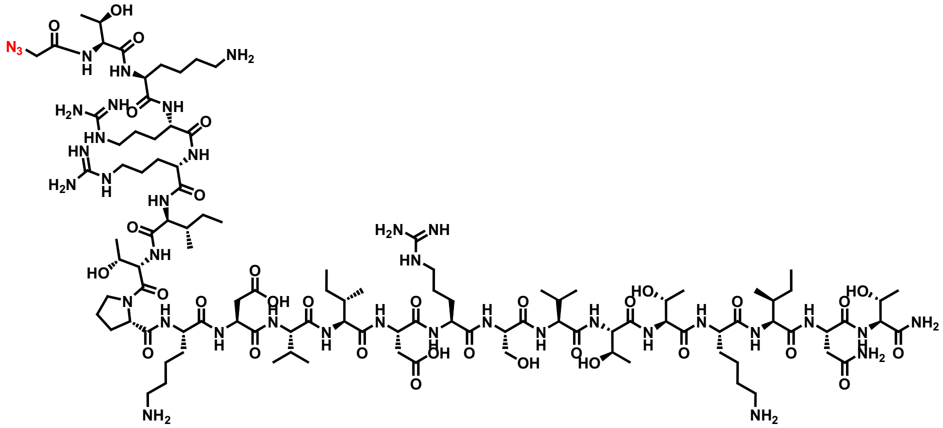

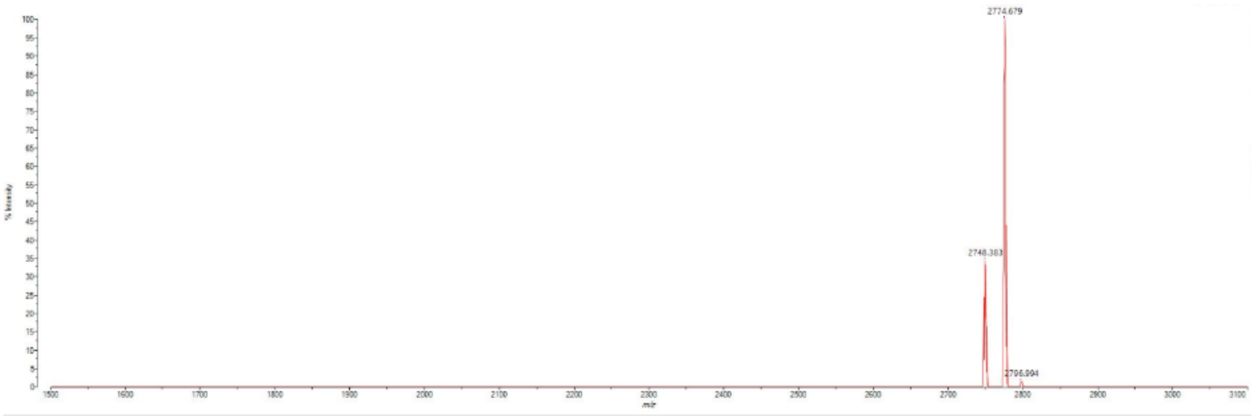

2774.679
2748.383
2796.994

**C1: 2BP100**

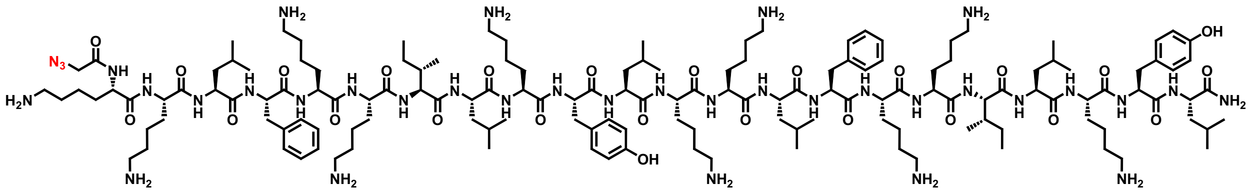

**C2: Rev (34-50)**

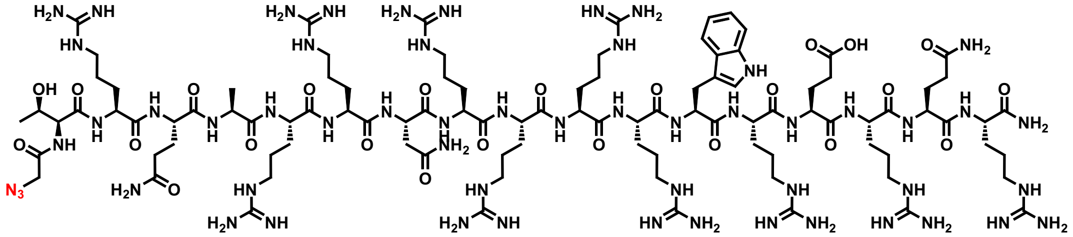

**C3: KH (9)**

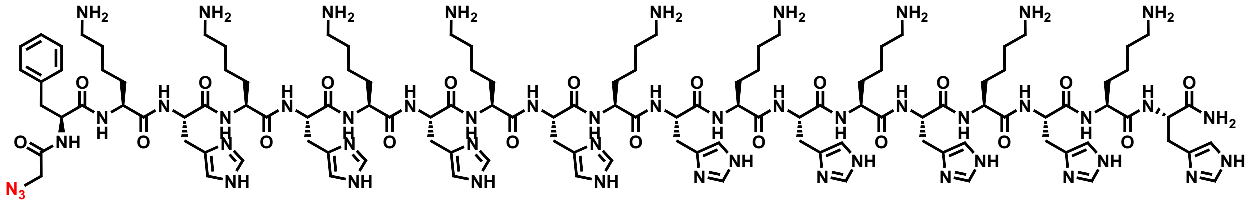

**C4: K9F**

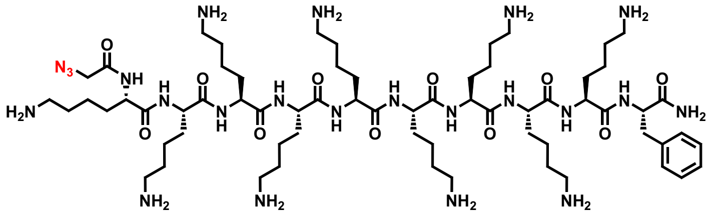

**C5: DPV3**

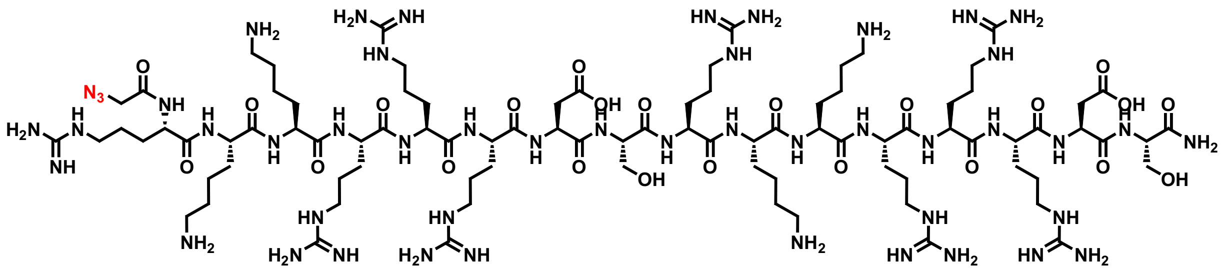

**C6: 6-Oct**

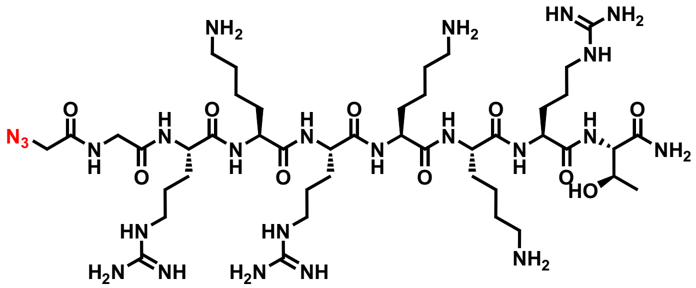

**C7: Tat (49-57)**

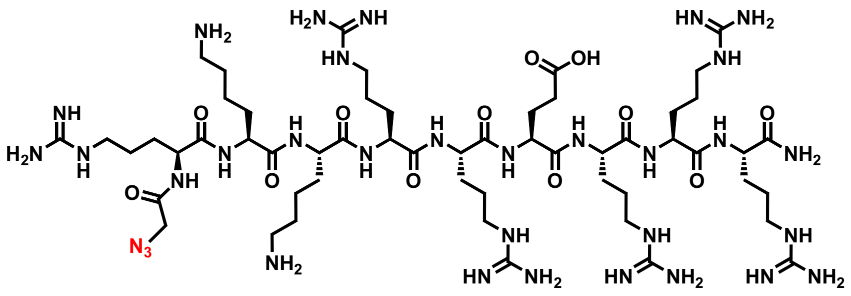

**C8: (KFF)3K**

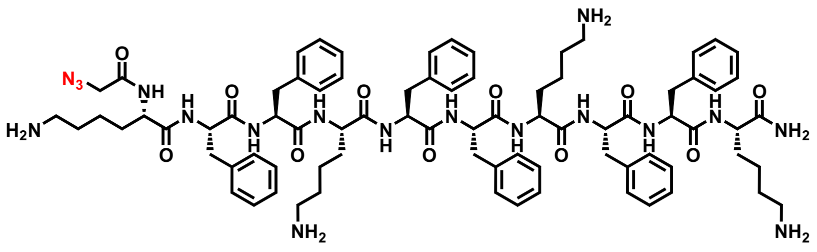

**C9: Pen2W2F**

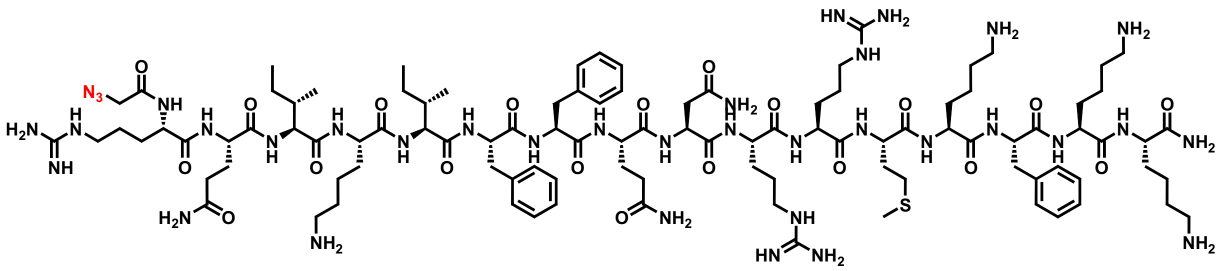

**C10: R5**

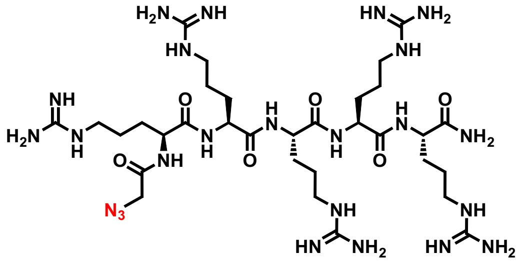

**C11: R7**

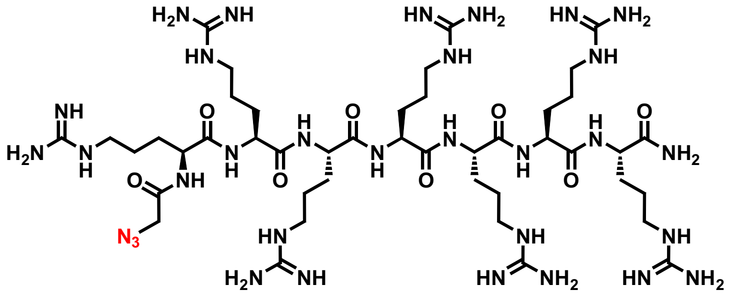

**C12: R9**

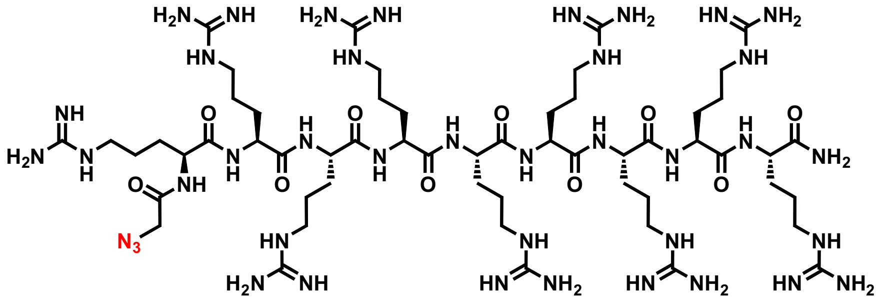

**C13: R11**

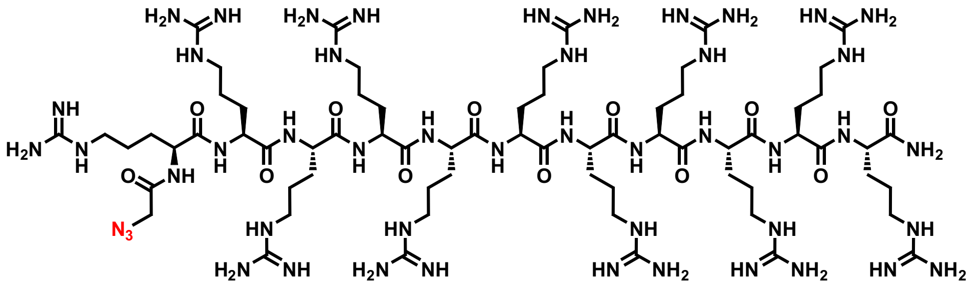

**A1: BP100**

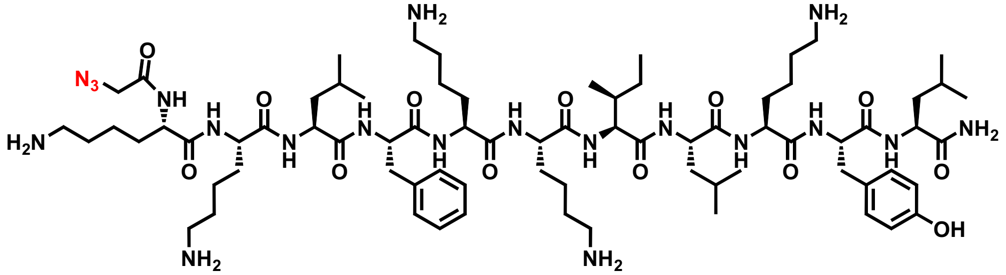

**A2: Sc 18**

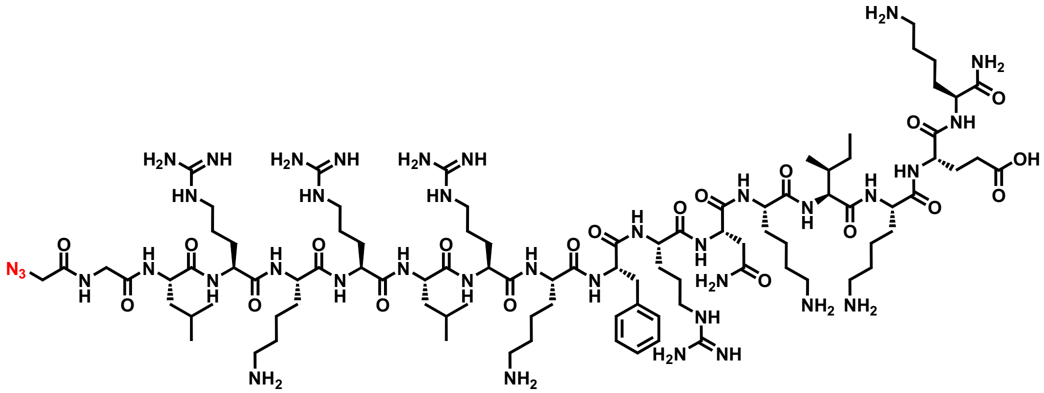

**A3: PenArg**

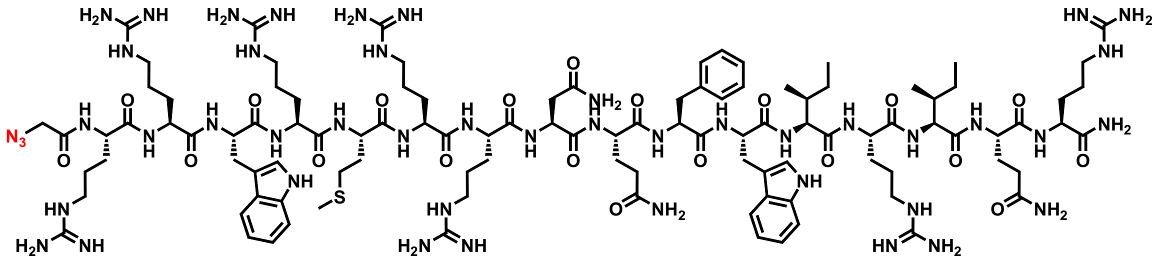

#### A4: Penetratin

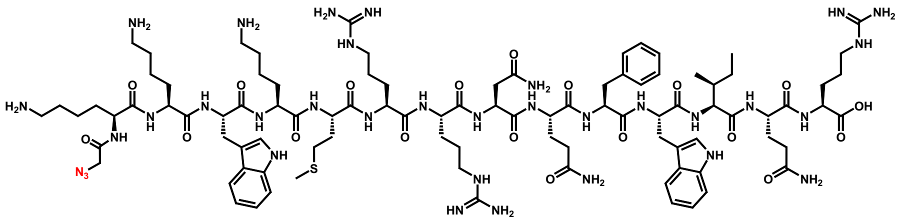

#### A5: Buforin II (5-21)

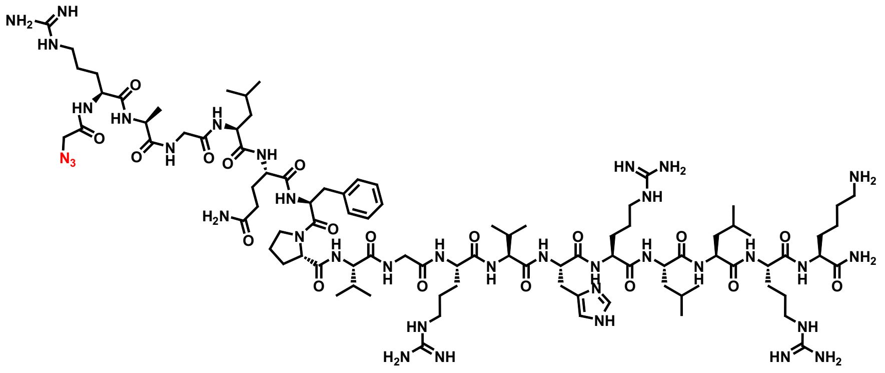

#### A6: pAntp (44-58)

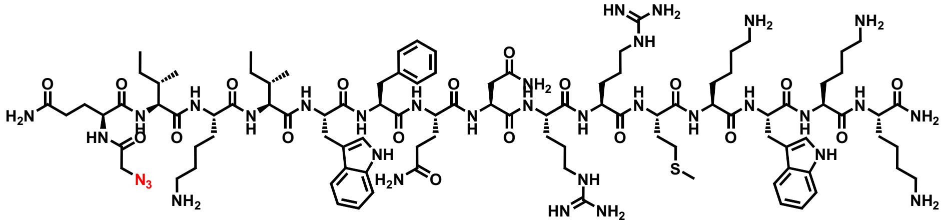

#### A7: Cyt C 77-101

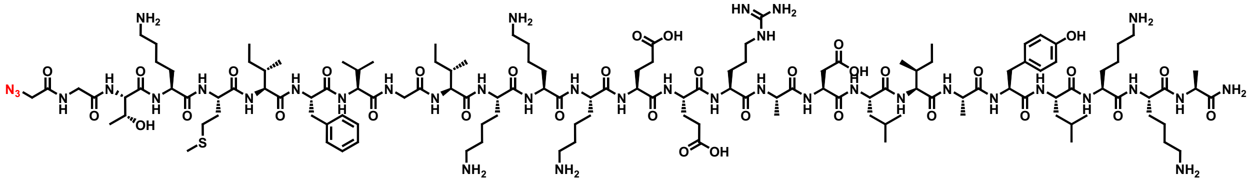

**A8: M511**

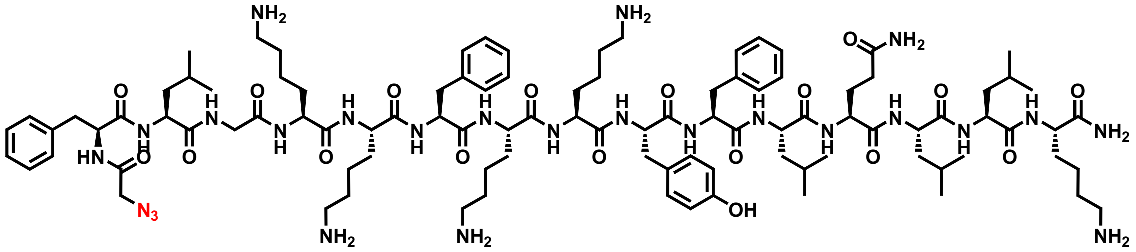

**A9: M591**

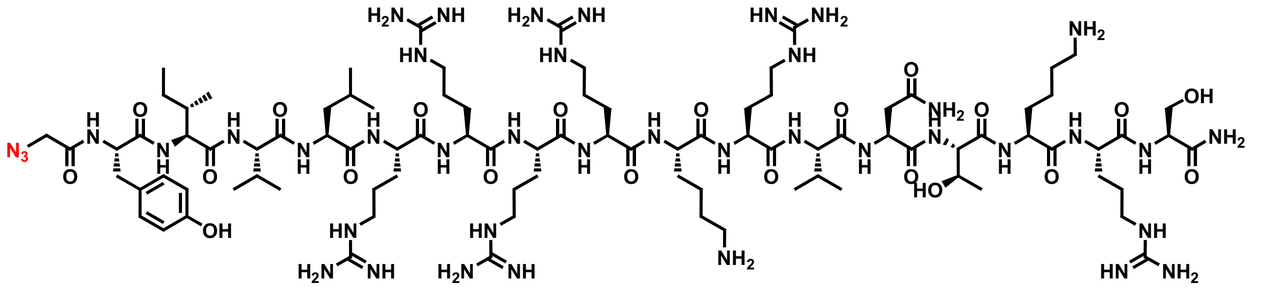

**A10: E162**

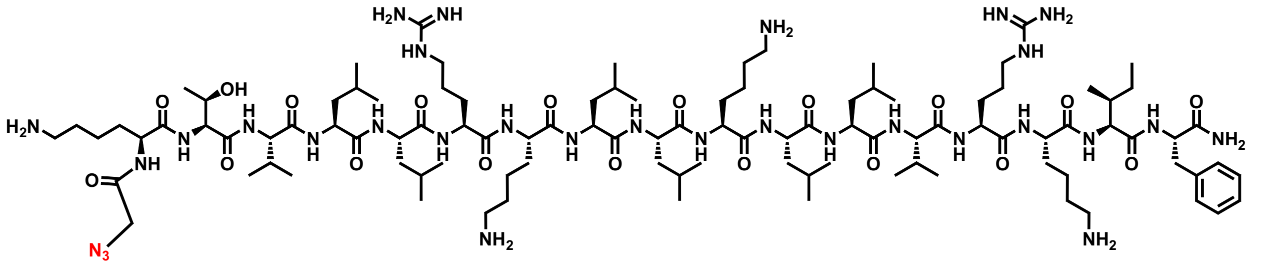

**A11: E165**

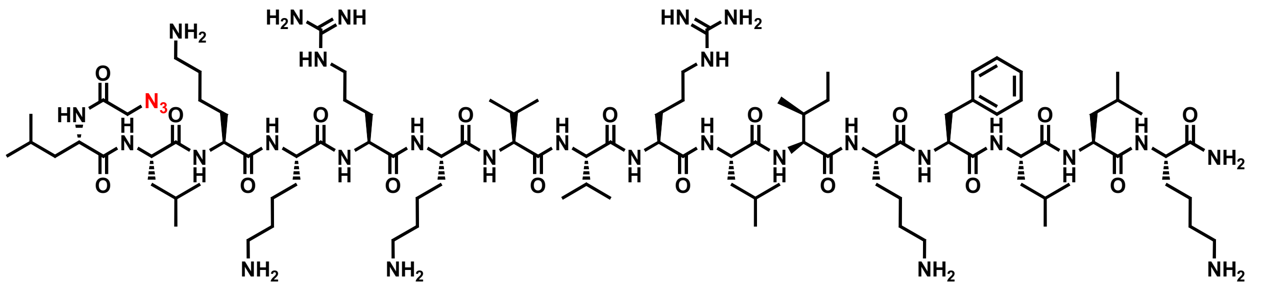

**A12: M867**

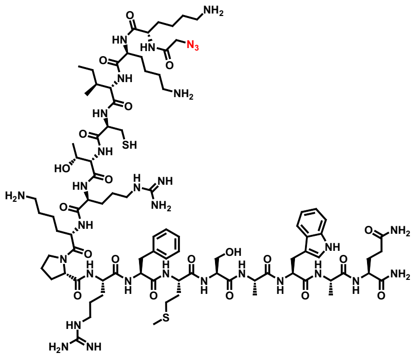

**A13: Crot (27-39)**

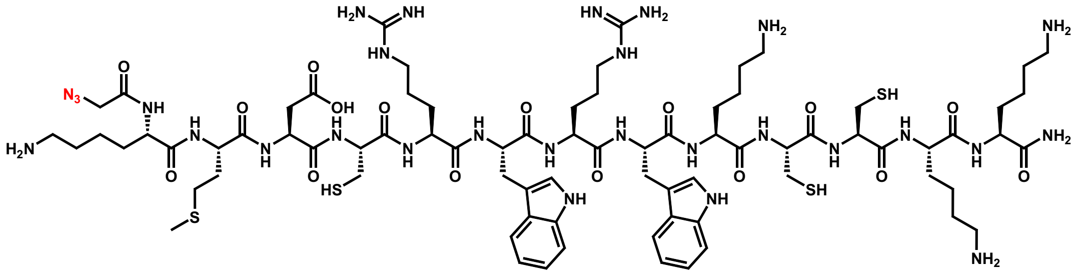

**A14: Crot (27-39) derivative 2**

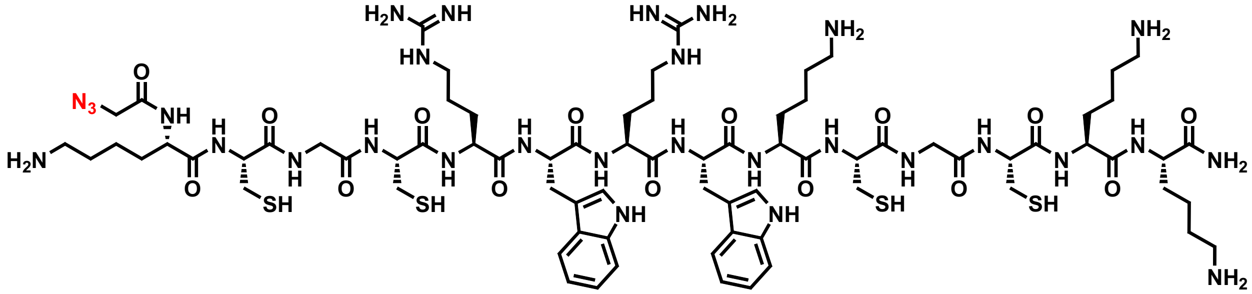

**A15: CyLoP-1**

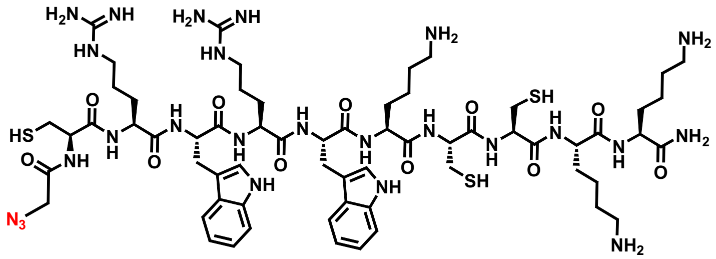

#### A16: HPV33L2-DD447

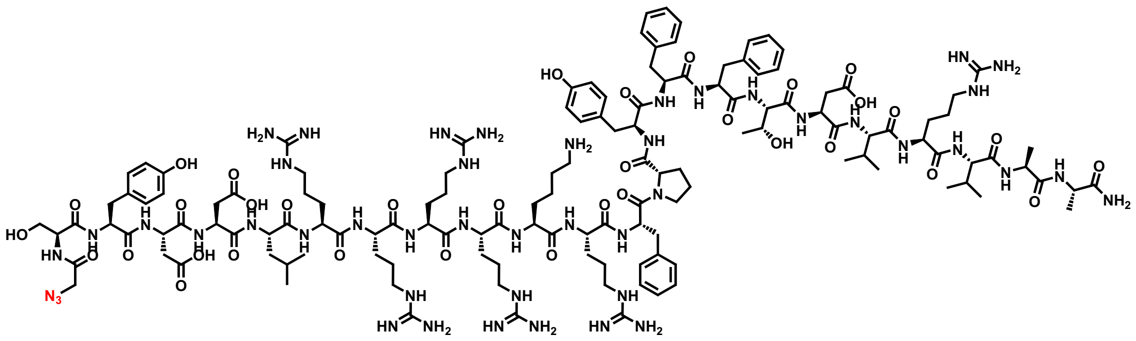

#### A17: HPV33L2-445/467

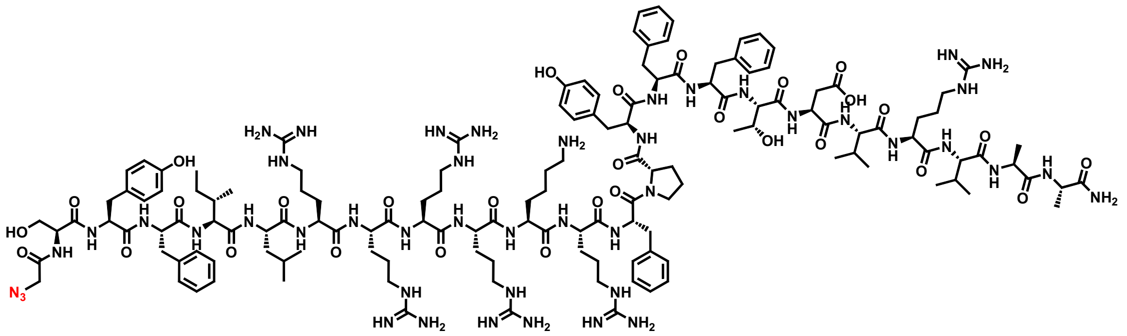

#### A18: KLA10

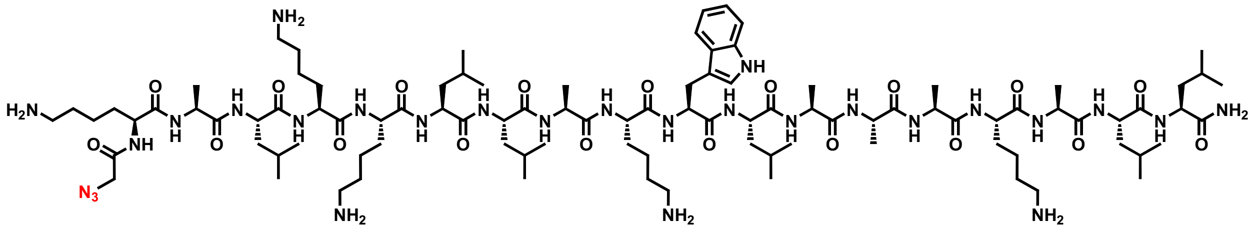

## A19: No. 14-12

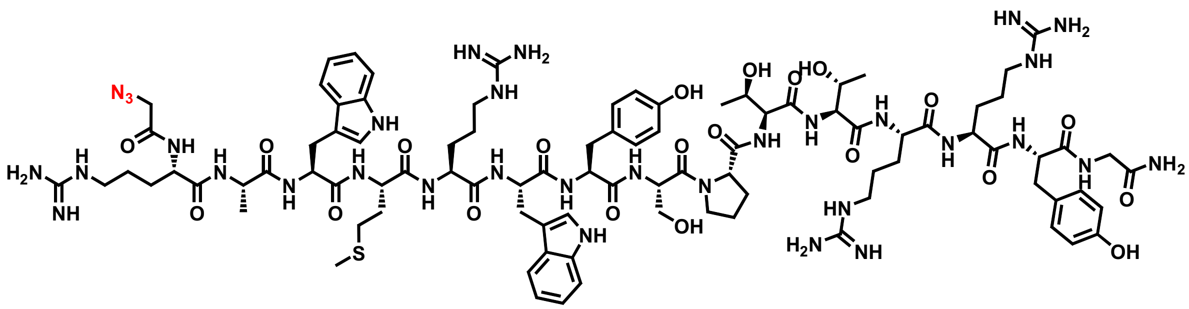

#### A20: dhvar2

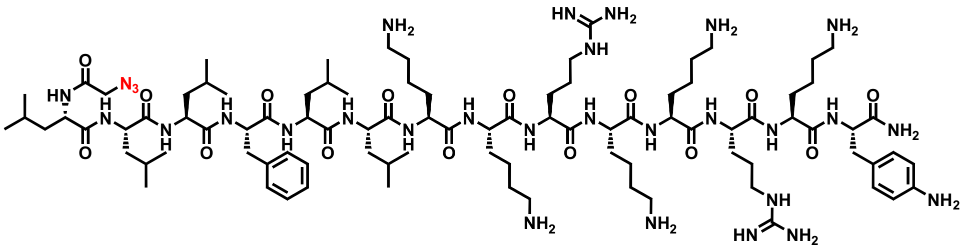


#### A21: ppTG1

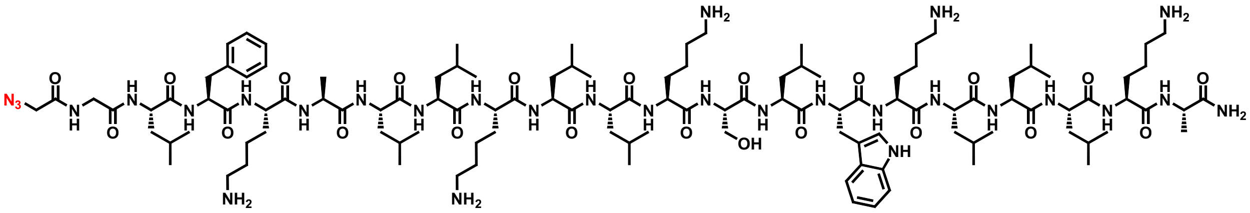


#### H1: PolyP3(SAP)

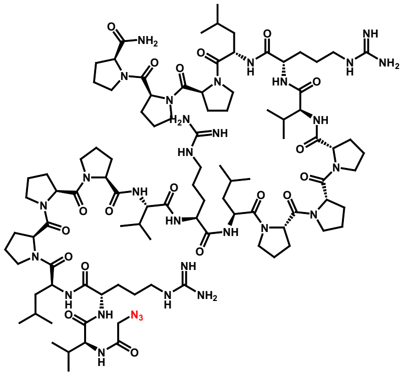


#### H2: LAH4

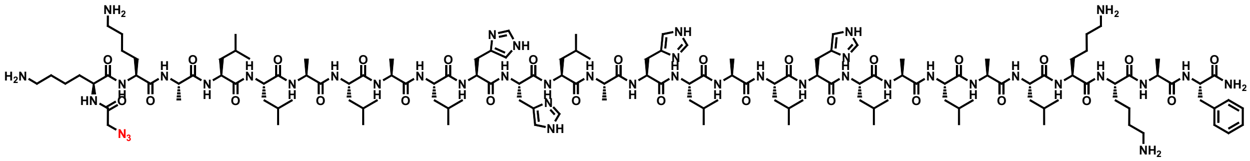

#### H3: Transportan

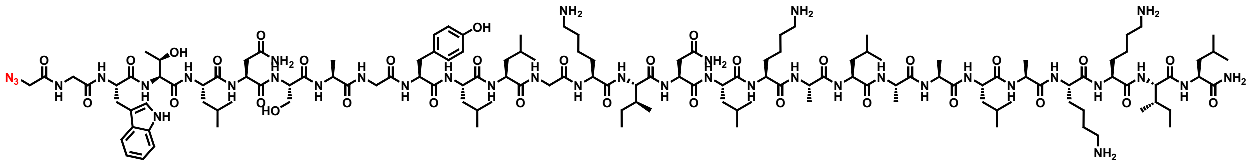

#### H4: InV3

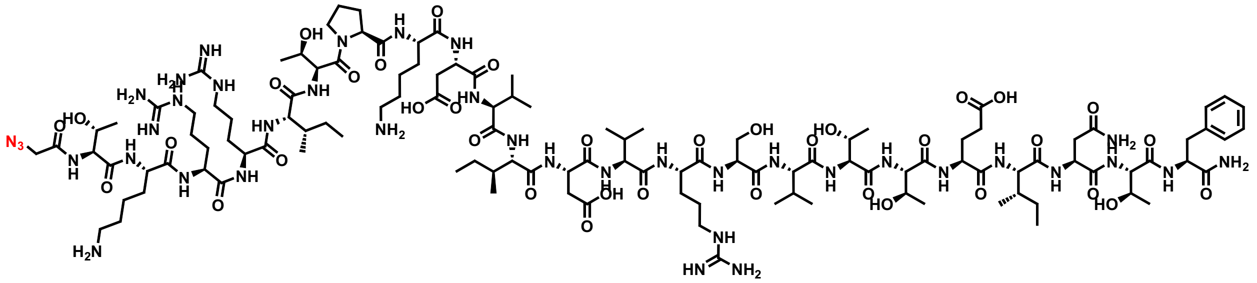

## H5: MG2d

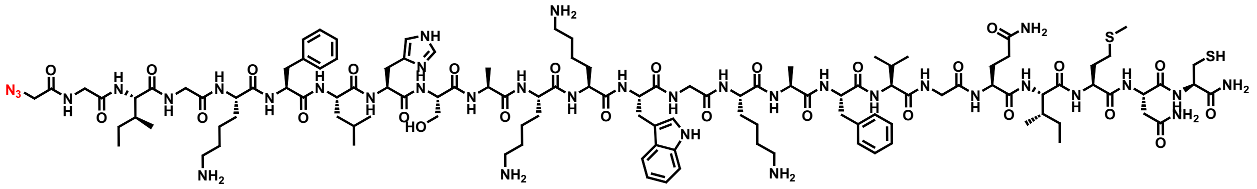

#### H6: InV3.5

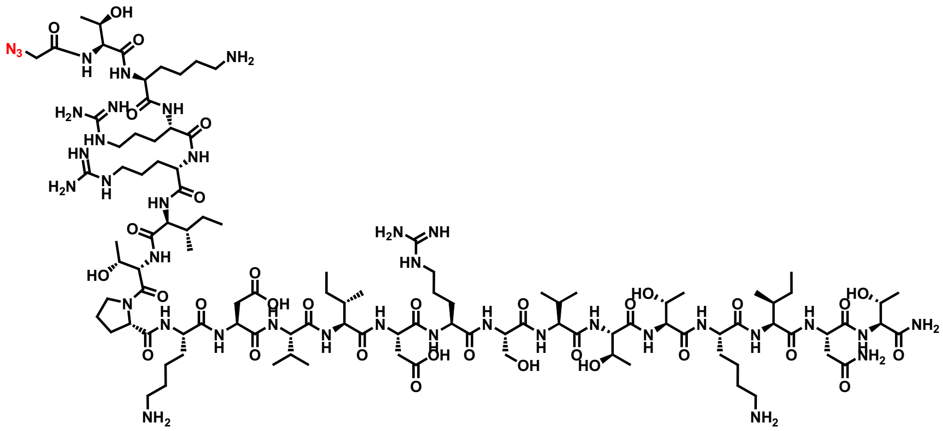

**Figure 1 a** Schematic representation of Barriers to cytosolic delivery of CPPs in bacterial and mammalian cells. **b**. General Scheme of Principle of CHAMP. Cells express HaloTag (HT) in the cytosol (top). Treatment with a DBCO-linked chloroalkane anchors a strained alkyne to HaloTag (middle). Azide-tagged test molecules that reach the cytosol react with the anchored DBCO by strain-promoted azide–alkyne cycloaddition and block it, and a subsequent azide fluorophore (green) labels the remaining DBCO sites (bottom). Fluorescence therefore decreases as cytosolic accumulation increases.

**Figure 2 a**. Chemical structures of the amphipathic (A1–A21), cationic (C1–C13) and hydrophobic (H1–H6) CPPs. The C-terminal azide tag is [highlighted in red]. **b**. Codes, names and amino acid sequences (one-letter code, N to C). Peptides were synthesized by solid-phase peptide synthesis [with free N-terminus and C-terminal azide].

**Figure 3 a**. Apparent accumulation measured by CHAMP in HaloTag-expressing HeLa cells for cationic (C1–C13), amphipathic (A1–A21) and hydrophobic (H1–H6) CPPs. HaloTag were treated with 50 μM of DBCOcl for 30 min, pulsed with 50 μM of azide-tagged CPPs for 1 h, chased with 50 μM TMRaz for 15 min. **b**. Viability of HeLa cells after treatment with each peptide at 50 μM for 2 h, measured by MTT assay and expressed as a percentage of untreated cells.

**Figure 4 a**. Measurement of cytosolic accumulation in *E.coli*. Apparent accumulation measured by CHAMP in HaloTag-expressing *E. coli* for cationic (C1–C13), amphipathic (A1–A21) and hydrophobic (H1–H6) CPPs. Cells were treated with 50 μM chloroalkane–DBCO (30 min), pulsed with 50 μM azide-tagged CPPs (1 h) and chased with 50 μM TMRaz (1 h). Apparent accumulation is 1 − FC, where FC is the fluorescence of peptide-treated cells divided by that of untreated cells; values below zero indicate fluorescence above the untreated control and are not interpreted as accumulation. **b**. Membrane integrity assessed by SYTO9 fluorescence of peptide-treated cells at 10 μM for 1 h shown as fold change over untreated cells. Polymyxin B. is used as positive control
